# Retrograde mitochondrial signaling governs the identity and maturity of metabolic tissues

**DOI:** 10.1101/2022.08.02.502357

**Authors:** Gemma L. Pearson, Emily M. Walker, Nathan Lawlor, Anne Lietzke, Vaibhav Sidarala, Jie Zhu, Tracy Stromer, Emma C. Reck, Ava M. Stendahl, Jin Li, Elena Levi-D’Ancona, Mabelle B. Pasmooij, Dre L. Hubers, Aaron Renberg, Kawthar Mohamed, Vishal S. Parekh, Irina X. Zhang, Benjamin Thompson, Deqiang Zhang, Sarah A. Ware, Leena Haataja, Stephen C.J. Parker, Peter Arvan, Lei Yin, Brett A. Kaufman, Leslie S. Satin, Lori Sussel, Michael L. Stitzel, Scott A. Soleimanpour

## Abstract

Mitochondrial damage is a hallmark of metabolic diseases, including diabetes and metabolic dysfunction-associated steatotic liver disease, yet the consequences of impaired mitochondria in metabolic tissues are often unclear. Here, we report that dysfunctional mitochondrial quality control engages a retrograde (mitonuclear) signaling program that impairs cellular identity and maturity across multiple metabolic tissues. Surprisingly, we demonstrate that defects in the mitochondrial quality control machinery, which we observe in pancreatic β cells of humans with type 2 diabetes, cause reductions of β cell mass due to dedifferentiation, rather than apoptosis. Utilizing transcriptomic profiling, lineage tracing, and assessments of chromatin accessibility, we find that targeted deficiency anywhere in the mitochondrial quality control pathway (*e.g.*, genome integrity, dynamics, or turnover) activate the mitochondrial integrated stress response and promote cellular immaturity in β cells, hepatocytes, and brown adipocytes. Intriguingly, pharmacologic blockade of mitochondrial retrograde signaling *in vivo* restores β cell mass and identity to ameliorate hyperglycemia following mitochondrial damage. Thus, we observe that a shared mitochondrial retrograde response controls cellular identity across metabolic tissues and may be a promising target to treat or prevent metabolic disorders.

## INTRODUCTION

Mitochondria are vital to cellular bioenergetics in eukaryotes, and mitochondrial defects in numerous tissues are associated with the development of metabolic diseases, such as type 2 diabetes (T2D), metabolic dysfunction-associated steatotic liver disease (MASLD), and metabolic dysfunction-associated steatohepatitis (MASH) (*1*). Insulin-producing pancreatic β cells from islet donors with T2D have dysmorphic mitochondrial ultrastructure as well as reductions in the ratio of ATP/ADP following glucose stimulation (*2, 3*). Skeletal muscle from individuals with T2D exhibits impaired mitochondrial ATP production in response to insulin, reduced mitochondrial respiratory capacity, and decreased mitochondrial DNA content (*4-8*). Visceral adipose tissue in insulin resistant individuals have significantly lower maximal mitochondrial respiration (*9*). Hepatic mitochondrial uncoupling and proton leak in T2D results in decreases in mitochondrial respiration as well as ATP synthesis and turnover (*9-12*). Numerous studies have highlighted the central role of broad mitochondrial impairments in the development of MASLD and MASH (reviewed in (*13-15*)), including increased mitochondrial membrane permeability, reduced ATP synthesis and fatty acid oxidation, increased mitochondrial (mt)DNA mutations, reduced mitochondrial quality control, and increased mitochondrial reactive oxygen species (mtROS). Pharmacologic agents to augment diverse aspects of mitochondrial health have also shown therapeutic potential for MASLD in clinical trials (*14*).

Lowell and Shulman first proposed mitochondrial dysfunction as a broad unifying mechanism in T2D, and features of numerous mitochondrial defects have since been described in myocardium, vascular endothelium, adipose tissue, and the central and peripheral nervous systems (*1, 16*). Importantly, the impact of mitochondria in T2D pathogenesis has been further bolstered by recent work determining that β cell mitochondrial gene expression and oxidative phosphorylation defects precede the development of T2D in humans (*17*). Moreover, several human genetic studies support links between mitochondria and T2D (*18-23*).

Loss of terminal cell identity has recently been observed in several tissues in T2D. Whereas the maintenance of pancreatic β cell mass was previously believed to depend solely on a balance of β cell replication and apoptosis (*24*), the loss of β cell differentiation or identity is now recognized as a key factor driving β cell failure in T2D (*25-27*). Dedifferentiated or immature β cells lack factors important in fully mature adult β cells, may acquire endocrine progenitor markers, and often express other islet cell hormones (*25-27*). Impairments in differentiation have also been observed in other metabolic tissues during T2D (*28-30*). However, despite the central importance of mitochondrial dysfunction to T2D, whether mitochondrial defects directly promote dedifferentiation or immaturity in metabolic tissues is unknown.

Here, we find that abnormalities in mitochondrial quality control engage a retrograde mitonuclear signaling program that promotes immaturity in metabolic tissues. Utilizing multiple independent mouse genetic models, metabolic reporters, lineage tracing approaches, and deep sequencing techniques, we observe that loss of mitophagy, mitochondrial genome stability, or mitochondrial fusion each elicit dedifferentiation in pancreatic β cells, ultimately leading to loss of β cell mass and hyperglycemia. Importantly, in multiple metabolic tissues—mouse β cells, hepatocytes, brown adipocytes, as well as in primary human islets—mitochondrial quality control defects result in a shared signature including loss of terminal maturity markers, induction of progenitor markers, and activation of the mitochondrial integrated stress response (ISR). Inhibition of the ISR *in vivo* restores β cell mass and maturity and ameliorates glucose intolerance following impaired mitochondrial quality control, illustrating that induction of retrograde mitonuclear communication directly promotes dedifferentiation of key metabolic tissues that may be essential to the development of metabolic disorders, including, but not limited to, T2D, MASLD, or MASH.

## RESULTS

### Mitochondrial quality control is impaired in β-cells in T2D

Abnormalities in mitochondrial structure, gene expression, and energetics have been described in several metabolic tissues in T2D, including pancreatic β-cells (*2, 3*),(*17*). Defects in mitochondrial structure, gene expression, and energetics in T2D could arise due to impairments in mitochondrial quality control or the mitochondrial life cycle, which tightly regulate mitochondrial genome integrity, biogenesis, dynamics, and turnover by mitophagy (*31, 32*). To investigate these mechanisms, we analyzed human islets from donors with T2D. Human islets from donors with T2D exhibited significant reductions in mtDNA content compared to non-diabetic donors (Fig. 1A). Single cell RNA sequencing (RNAseq) of the β-cells from human islets revealed significantly reduced expression of 11 of the 13 mitochondrial-encoded RNAs (mtRNAs) in donors with T2D (Fig. 1B). Of note, reduced mtRNA expression was β-cell specific as there were no differences in mtRNA levels in α-cells or other non β islet cells in donors with T2D (fig. S1A and (*33*)). We also did not observe differences in mitochondrial mass in islets between donors with or without T2D, as measured by citrate synthase activity as well as western blot analysis for OXPHOS subunit proteins and the outer mitochondrial membrane protein TOM20, suggesting that reduced mtDNA content and mitochondrial gene expression were not secondary to a decline in mitochondrial mass (fig. S1B-C).

**Figure 1.**
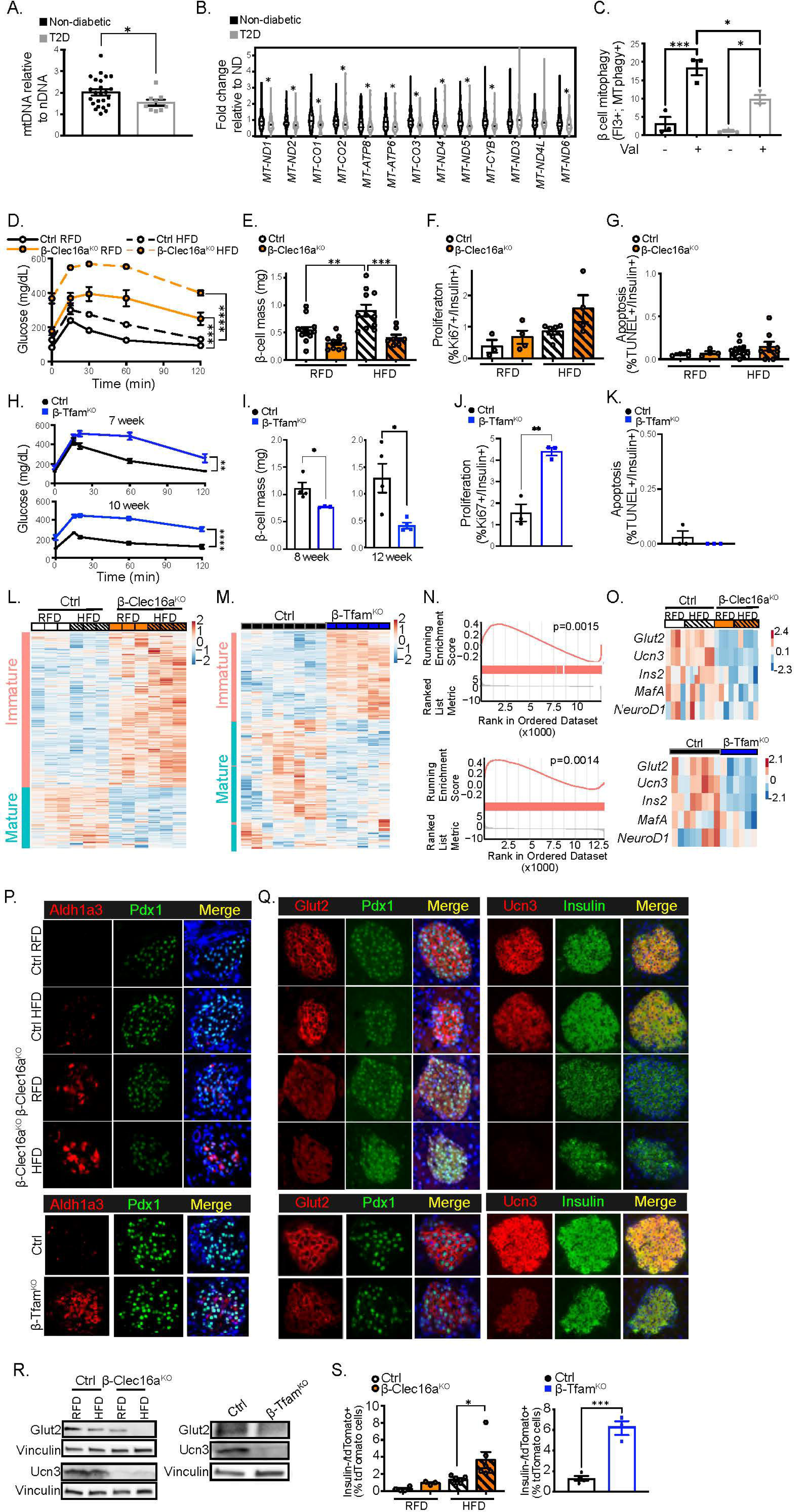
Mitochondrial quality control is impaired in β-cells in T2D and is required for maintenance of glucose homeostasis and β-cell mass by maintaining β-cell maturity. (A) mtDNA:nuclear DNA ratio analysis of human islets from non-diabetic donors (n=24) or those with T2D (*n* = 9). \**P* < 0.05 by unpaired Mann-Whitney test. (B) Violin plot showing levels of mitochondrially encoded RNA from single β-cell analysis (*33*) of human non-diabetic donors (ND) or those with T2D (*n* = 3-5/group). \**P* < 0.05 by unpaired Mann-Whitney test. (C) Flow cytometry quantification of percentage of β-cells (Fluozin-3+) containing MTPhagy+ mitochondria from human islets of non-diabetic donors or those with T2D (*n* = 3-4/group). \**P* < 0.05, \*\*\**P* < 0.001 by one-way ANOVA Tukey’s multiple comparison post-test. At least 2500 Fluozin-3+/DAPI-events quantified per sample. (D) Blood glucose concentration measured during an IPGTT of Ctrl or β-Clec16a^KO^ mice following 12-weeks regular fat diet (RFD) or high fat diet (HFD). *n* = 10-18mice per group. \*\*\**P* < 0.001 Ctrl RFD vs β-Clec16a^KO^ RFD, \*\*\*\**P* < 0.0001 Ctrl HFD vs β-Clec16a^KO^ HFD by two-way ANOVA. (E) Pancreatic β-cell mass from 12-week diet fed (16-week-old) Ctrl or β-Clec16a^KO^ mice. *n* = 8-13 mice per group. \*\**P* < 0.01; \*\*\**P* < 0.001 by one-way ANOVA, Tukey’s multiple comparison post-test. (F) β-cell proliferation measured as the % of Ki67+/Insulin+ cells. *n* = 3-6 mice per group. (G) β-cell apoptosis measured as the % of TUNEL+/Insulin+ cells. n=4-11 mice per group. (H) Blood glucose concentrations during an IPGTT of Ctrl or β-Tfam^KO^ mice at 7- or 10-weeks of age. *n* = 8-14 animals per group. \*\**P* < 0.01, \*\*\*\**P* < 0.0001 by two-way ANOVA effect of genotype. (I) Pancreatic β-cell mass from 8- or 12-week-old Ctrl or β-Tfam^KO^ mice. *n* = 3-4 mice per group. \**P* < 0.05 by Student’s unpaired *t*-test. (J) β-cell replication measured as the % of Ki67+/Insulin+ cells in 8-week-old mice. *n* = 3 mice per group. \*\**P* < 0.01 by Student’s unpaired *t*-test. (K) β-cell apoptosis measured as the % of TUNEL+/Insulin+ cells. *n* = 3 mice per group. \**P* < 0.05 by Student’s unpaired *t*-test. Expression heatmap of islet RNAseq from (L) Ctrl versus β-Clec16a^KO^ islets or (M) Ctrl versus β-Tfam^KO^ islets demonstrating expression of immature or mature gene markers (gene sets generated from (*50*)). (N) Gene set enrichment analysis (GSEA; gene set generated from (*50*)) for expression of immature genes in β-Clec16a^KO^ versus Ctrl islets (top) or β-Tfam^KO^ versus Ctrl islets (bottom). (O) Heatmap displaying expression of core β-cell identity markers from Ctrl versus β-Clec16a^KO^ islets (top) and Ctrl versus β-Tfam^KO^ mice (bottom). (P) Representative immunofluorescence images for key β-cell dedifferentiation marker Aldh1a3 (red), Pdx1 (green) from Ctrl and β-Clec16a^KO^ islets (top) and Ctrl and β-Tfam^KO^ mice (bottom). *n* = 3-6/group. (Q) Representative immunofluorescence images for key β-cell maturity markers. Glut2 (red), Pdx1 (green); Ucn3 (red), Insulin (green) in Ctrl and β-Clec16a^KO^ islets (top) and Ctrl and β-Tfam^KO^ mice (bottom). *N* = 3-6/group. (R) Protein expression of Glut2 and Ucn3 by WB in Ctrl and β-Clec16a^KO^ islets (left) and Ctrl and β-Tfam^KO^ mice (right). *n* = 3-4/group. (S) Quantification of Insulin-negative tdTomato-positive cells from *Ins1*-Cre;Rosa26-tdTomato and β-Clec16a^KO^;Rosa26-tdTomato and *Ins1*-Cre;Rosa26-tdTomato (left) and β-Tfam^KO^;Rosa26-tdTomato mice (right). *n* = 3-4 mice per group; \**P* < 0.05; \*\*\**P* < 0.001 by Student’s unpaired *t*-test.

Previously published ultrastructural studies have already established that defects in mitochondrial architecture are apparent in T2D, suggestive of alterations in mitochondrial dynamics in T2D (*2, 3*). Thus, we next turned our focus to an evaluation of mitophagic flux in single cells by flow cytometry analysis as per our previous approaches (*34-36*). In β-cells from non-diabetic donors, the potassium ionophore valinomycin induced the expected increase in mitophagic flux, defined by cells with increases in depolarized mitochondria within the acidic lysosome compartment (Fig. 1C). T2D β cells exhibited lower mitophagic flux following valinomycin exposure, consistent with impairments in mitophagy (Fig. 1C). Notably, defects in mitophagy were not observed in non-β cells from T2D donors (fig. S1D), again corroborative of β-cell specific mitochondrial defects in T2D. Taken together, these results indicate that islets in T2D exhibit specific defects in mtDNA content, β cell mtRNA levels, and β cell mitophagy, in addition to previously published mitochondrial structural defects, consistent with comprehensive impairments in mitochondrial quality control.

To determine whether changes in β-cell mitochondrial quality control in T2D are secondary to obesity, we measured mtDNA content and mitophagy in transgenic mice expressing the mt-Keima pH sensitive mitophagy reporter (*37, 38*) fed a regular fat diet (RFD) or high fat diet (HFD) for up to one year of age. There were no significant differences in islet mtDNA content secondary to age or diet (fig. S1E). We did observe a small, yet significant reduction in the rates of mitophagy in β-cells following diet-induced obesity (DIO) that were not observed in non-β-cells, as measured by flow cytometry (fig. S1F-G). Further, a previously published report demonstrated that obesity did not induce changes in β-cell mitochondrial architecture in mice (*35*). Taken together, these findings suggest the peripheral effects of obesity or insulin resistance alone do not recapitulate the changes in β-cell mitochondrial quality control found in T2D.

### Mitochondrial quality control is necessary to maintain β-cell mass and function

To determine whether impaired mitochondrial quality control is sufficient to lead to β-cell failure, as observed in T2D, we generated mouse genetic models deficient in components of the mitochondrial quality control machinery (though not necessarily to model T2D *per se*). We hypothesized that using specific models of impaired mitophagy or mitochondrial genome integrity in isolation might allow us to parse the individual contribution of each mitochondrial quality control defect to β-cell failure in T2D. We first generated mice bearing β-cell-specific deletion of Clec16a (*Clec16a*^f/f^; *Ins1*-Cre, hereafter known as β-Clec16a^KO^; fig. S2A), a critical regulator of mitophagy that is essential to maintain proper mitochondrial clearance and respiratory function (*39*). Notably, expression of *Clec16a* is reduced in islets from donors with T2D, and *Clec16a* is also a direct transcriptional target of the T2D gene *Pdx1* (*40-42*). β-Clec16a^KO^ mice as well as controls were fed RFD or HFD for 12 weeks prior to physiologic or morphometric assays, and we did not observe differences in insulin sensitivity between genotypes (fig. S2B). Of note, *Ins1*-Cre and floxed-only controls were phenotypically indistinguishable from each other, exhibiting no differences in glucose tolerance or body weight following DIO (fig. S2C-D), and thus were pooled as controls for subsequent analyses, consistent with previous studies from our group and others (*35, 43*). RFD-fed β-Clec16a^KO^ mice developed glucose intolerance consistent with our previous studies (Fig. 1D and (*39,77*)). We found in the setting of DIO that this phenotype was markedly exacerbated (Fig. 1D). β-Clec16a^KO^ animals had reductions in serum insulin concentrations upon HFD feeding and lower insulin release *in vivo* after a glucose challenge compared to HFD-fed controls, as well as modest changes in OXPHOS subunit expression (fig. S2E-G). We next evaluated mitochondrial ultrastructure by transmission electron microscopy (*44*) and mitochondrial morphology and network complexity by immunostaining for the mitochondrial marker SDHA, utilizing previously validated algorithms (*22,23)* applied as per our previous work (*35*). Following β-Clec16a-deficiency, we observed abnormal mitochondrial ultrastructure with distorted cristae and inclusions (fig. S2H). Further, in HFD β-Clec16a^KO^ mice we observed increases in mitochondrial sphericity as well as reductions in network/branch complexity that are consistent with previous observations of impaired mitochondrial clearance (fig. S2I-N and (*36*)).

We next evaluated β-cell mass in mitophagy-deficient mice and observed that β-Clec16a^KO^ animals failed to expand their β-cell mass to compensate for diet-induced insulin resistance (Fig. 1E). We did not observe differences in α-cell mass between the groups (fig. S2O). To understand the etiology of reduced β-cell mass in HFD-fed β-Clec16a^KO^ mice, we assessed markers of β-cell proliferation and apoptosis. Surprisingly, we observed a trend toward increased β-cell replication with no changes in apoptosis in HFD-fed β-Clec16a^KO^ mice (Fig. 1F-G). Thus, our data led us to speculate that another etiology may contribute to the loss of β-cell mass in this model.

As aging is also a risk factor for the development of T2D (*45*), we evaluated β-Clec16a^KO^ mice at one year of age. β-Clec16a^KO^ mice developed severe glucose intolerance and reduced β-cell mass with age with no change in proliferation or apoptosis, similar to the effect of HFD feeding (fig. S2P-S). To ensure the physiologic and histologic changes we observed were not attributable to developmental defects, we generated inducible β-cell specific Clec16a knockout animals by intercrossing our mice bearing the *Clec16a* conditional allele with the tamoxifen-inducible MIP-CreERT strain (*Clec16a*^f/f^; *MIP*-CreERT, hereafter known as iβ-Clec16a^KO^ mice). Following tamoxifen-mediated recombination at 7 weeks of age, iβ-Clec16a^KO^ mice exhibited mild glucose intolerance on RFD that was exacerbated upon HFD feeding, reductions in β-cell mass on HFD, and no differences in β-cell replication or survival when compared to *MIP*-CreERT controls (fig. S2T-W), phenocopying constitutive β-cell Clec16a knockouts. Taken together, these data indicate that mitophagy is vital for the preservation of β-cell mass following aging or obesity, independent of changes in cell replication or survival and not attributable to defects in β-cell development.

Next, we generated mice bearing reductions of mtDNA content following loss of Tfam, a crucial regulator of mitochondrial genome integrity, mtDNA copy number control, and mtRNA transcription, by intercrossing *Ins1*-Cre and *Tfam*^f/f^ mice (hereafter noted as β-Tfam^KO^; fig. S3A). We chose the *Ins1*-*Cre* knockin strain to alleviate concerns related to off-target and non-specific effects of the transgenic *RIP2*-Cre strain previously employed to knockout *Tfam* (*46, 47*). Of note, reductions in *Tfam* expression were previously observed in the β-cells of donors with T2D, and *Tfam* has also been reported to be a direct transcriptional target of the T2D gene *Pdx1* (*33, 48*).

We first observed the expected reductions of mtDNA in β-Tfam^KO^ islets (fig. S3B). To confirm that mtDNA depletion was specific to β-cells, we additionally intercrossed β-Tfam^KO^ mice with the ROSA26-lox-STOP-lox-tdTomato reporter strain, such that β-cells were irreversibly labeled with the tdTomato reporter following Cre-mediated recombination (*49*). Following fluorescence-activated cell sorting of tdTomato-positive β-cells, we observed near complete depletion of mtDNA content in β-Tfam^KO^ mice, while tdTomato-negative cells were unaffected (fig. S3C). Additionally, β-Tfam^KO^ islets had significant changes in OXPHOS protein expression (fig. S3D). An evaluation of mitochondrial morphology and network complexity in TFAM-deficient β-cells revealed increases in mitochondrial volume, decreased sphericity, and increases in branch number, branch length, and branch junctions (fig. S3E-J). These data bear resemblance to the increases in mitochondrial density previously reported in β-cells from islet donors with T2D (*2*) and also could suggest a connection between alterations in mitochondrial density and reduced mtDNA content or mitochondrial gene expression in T2D (Fig. 1A-B).

Similar to Clec16a-deficient mice, β-Tfam^KO^ mice showed progressive glucose intolerance with age, decreased insulin secretion following glucose administration, and reduced β-cell mass without changes in α-cell mass (Fig. 1H-I and fig. S3K-M). Interestingly, β-Tfam^KO^ mice also exhibited a significant increase in β-cell replication at 8 and 12 weeks of age (Fig. 1J and fig. S3N) with no change in apoptosis at 8 weeks and only a small increase at 12 weeks of age (Fig. 1K and fig. S3O). Further, no increases in β-cell apoptosis were previously observed following *RIP2*-Cre mediated deletion of *Tfam* (*46*). Analogous to our findings in Clec16a-deficient mice, these results suggest that loss of Tfam reduces β-cell mass, and that this reduction is unlikely to be primarily mediated by β-cell replication or survival. Our findings indicate that defects in mitochondrial quality control promote the progressive loss of β-cell mass and function, which bear resemblance to what is observed in T2D.

### Defects in mitochondrial quality control induce loss of β-cell identity

We next asked whether a shared mechanism explained the similarities in phenotypic responses to the orthogonal approaches we used to impair mitochondrial quality control. First, we performed RNAseq on isolated islets from RFD- and HFD-fed β-*Clec16a*^KO^ mice as well as β-Tfam^KO^ mice and their respective controls. Overlaying our results with a previously validated β-cell maturity/immaturity gene set (*50*) revealed significant upregulation of genes associated with β-cell immaturity and corresponding downregulation of genes associated with mature β-cells in both RFD and HFD-fed β-Clec16a^KO^ mice as well as islets from β-Tfam^KO^ mice (Fig. 1L-M). Gene set enrichment analysis (GSEA) confirmed that genes induced in Clec16a- or Tfam-deficient mice were significantly enriched for markers of β-cell immaturity (Fig. 1N). Additionally, pathway analysis identified specific divergences between islets from control and β-Clec16a^KO^ mice. Clec16a-deficient islets displayed downregulation of genes related to insulin secretion and mitochondrial lipid metabolism (β-oxidation and mitochondrial long chain fatty acid oxidation), suggestive of maladaptive β-cell metabolic function. Clec16a-deficient islets also demonstrated increased enrichment for developmental or morphogenetic pathways, including Notch and Id signaling (fig. S4A-B; (*51-54*)). Similarly, pathway analysis of β-Tfam^KO^ islets displayed downregulation of insulin secretion and key β-cell signature pathways (insulin secretion, maturity onset diabetes of the young, and type 2 diabetes mellitus) when compared to controls (fig. S5A). Moreover, reduced expression of core β-cell maturity/identity markers, including *Ins2*, *Ucn3*, *Glut2*, and *MafA*, and upregulation of immaturity markers (*50, 55, 56*) were commonly observed in both genetic models (Fig. 1O and fig. S5B-C).

Reduced expression of core β-cell maturity markers and increased markers of immaturity were also apparent in islets from mice bearing β-cell-specific deletion of Mitofusins 1 and 2, a recently described model of impaired mitochondrial quality control that abrogates mitochondrial fusion (β-Mfn1/2^DKO^ mice; fig. S5D-E; (*35*)). Loss of Mfn1/2 also leads to modest reductions in β-cell mass with age (*57*). We extended our understanding of the relevance of mitochondrial fusion on β-cell mass following DIO, first noting that β-Mfn1/2^DKO^ mice exhibited substantial defects in glucose tolerance and insulin secretion, as well as expected decreases in expression of OXPHOS subunits within complexes I, III, and IV (fig. S5F-H and (*35)*). Importantly, consistent with Clec16a and Tfam-deficient animals, HFD-fed β-Mfn1/2^DKO^ mice displayed a loss of β-cell mass, with increased β-cell proliferation without increases in β-cell apoptosis (fig. S5I-K).

To verify changes in β-cell maturity, we first performed immunostaining on pancreatic sections. The dedifferentiation marker Aldh1a3, which is not observed in terminally differentiated β-cells but has been found in islets of donors with T2D (*25, 58*), was detectable in RFD-fed β-Clec16a^KO^ mice and further increased upon HFD feeding or aging (Fig. 1P and fig. S6A). Aldh1a3 was also upregulated in β-Tfam^KO^, iβ-Clec16a^KO^, and β-Mfn1/2^DKO^ β-cells (Fig. 1P and fig. S6B-C). Further, key β-cell maturation markers Glut2 and Ucn3 were markedly decreased in each of these models of impaired mitochondrial quality control by immunostaining as well as western blot compared with littermate controls (Fig. 1Q-R and fig. S6D-H). Stem cell or early endocrine progenitor cell markers such as *Oct4*, *Sox2*, *Nanog*, and *Ngn3* were not upregulated in islets from β-Clec16a^KO^, β-Tfam^KO^, or β-Mfn1/2^DKO^ mice (data not shown). Genes related to the Parkin or mTOR pathways were also unaffected in β-Clec16a^KO^, β-Tfam^KO^, or β-Mfn1/2^DKO^ islets (fig. S7). These findings demonstrate that loss of mitochondrial quality control induces markers of immaturity and loss of terminal β-cell identity without a signature associated with regression to early developmental precursors.

We next employed genetic lineage tracing to confirm the loss of β-cell identity in Clec16a and Tfam-deficient mice. As noted above, both β-*Clec16a*^KO^ and β-Tfam^KO^ mice (and *Ins1*-*Cre* control littermates) were intercrossed with the ROSA26-lox-STOP-lox-tdTomato reporter strain (*49*), such that β-cells were irreversibly labeled with the tdTomato reporter following Cre-mediated recombination. By this lineage tracing strategy, loss of β-cell identity is predicted to result in insulin-negative, tdTomato-positive cells. As expected, >95% of β-cells were positive for both insulin and tdTomato in *Ins1*-Cre control mice, consistent with previous reports (fig. S8A-B and (*43*)). No changes in tdTomato lineage allocation were observed between RFD-fed β-*Clec16a*^KO^ mice and controls (Fig. 1S). However, HFD-fed β-*Clec16a*^KO^ mice had significantly more insulin-negative tdTomato-positive cells compared to HFD-fed controls (Fig. 1S and fig. S8A). We also observed a robust increase in insulin-negative tdTomato-positive cells in Tfam-deficient mice compared to controls (Fig. 1S and fig. 8SB). Indeed, the frequency of altered β-cell identity in Tfam-deficient mice was similar to the frequency previously reported following loss of the critical β-cell transcription factor Nkx2.2 (*59*), supportive of a profound impact of mtDNA depletion on β-cell identity. Surprisingly, we also observed a significant increase in glucagon-positive, tdTomato-positive cells in β-Tfam^KO^ mice, and a similar trend in HFD-fed β-*Clec16a*^KO^ mice, which could indicate that dedifferentiated β-cells acquire glucagon expression (fig. S8C-D). These complementary studies in several models resembling discrete mitochondrial quality control defects found in T2D, converge to indicate that impaired mitochondrial quality control leads to loss of β-cell identity.

### Defective mitochondrial quality control induces the ISR

We hypothesized that impairments in mitochondrial quality control may induce a common mitochondrial-nuclear signaling circuit leading to a transcriptional response consistent with the induction of β-cell dedifferentiation/immaturity. To determine if a common transcriptional signature is activated by impaired mitochondrial quality control, we performed a comparative analysis of the top 500 up- and down-regulated genes by RNAseq across our models of mitophagy deficiency (β-Clec16a^KO^), mtDNA depletion (β-Tfam^KO^) and defective fusion (β-Mfn1/2^DKO^). Interestingly, we found 74 upregulated and 28 downregulated genes in common across all three models (Fig. 2A and table S1). To search for connections between these targets, we next analyzed these 102 commonly differentially expressed genes by STRING (Search Tool for the Retrieval of Interacting Genes/Proteins; (*60*)), which revealed an unexpected interaction relating to key targets in the β-cell maturity (*Ins2*, *Ucn3*, *MafA*, and *Slc2a2*, which encodes Glut2) and ISR pathways (*Trib3*, *Ddit3*, *Atf3*, *Fgf21*, *Cebpβ*, and *Gdf15*) (Fig. 2B).

**Figure 2.**
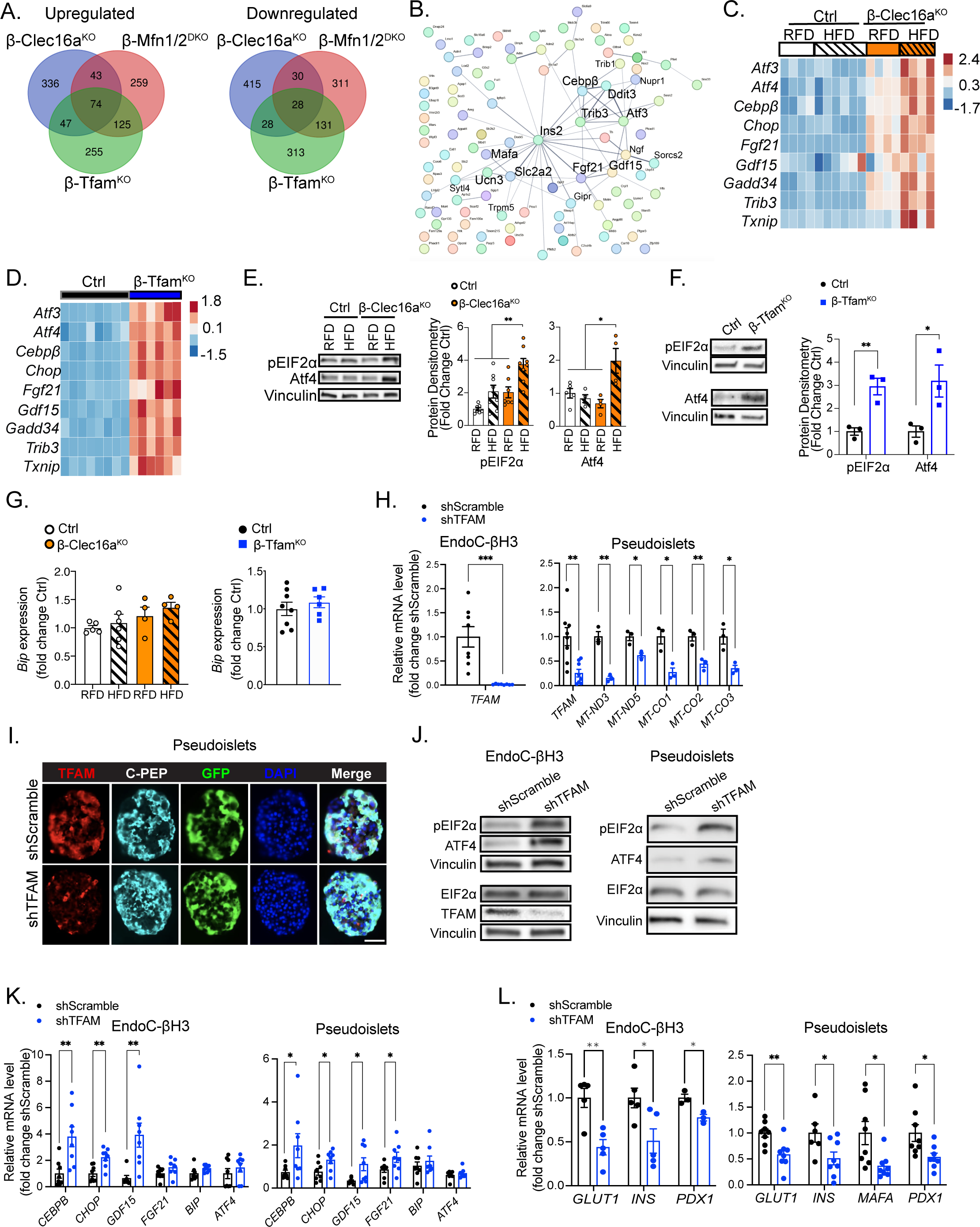
Loss of mitophagy, mtDNA content, or mitochondrial fusion activates a shared integrated stress transcriptional response in mouse and human islets. Comparative analysis of the top 500 upregulated (left) and the top 500 downregulated (right) genes across 3 models of β-cell-specific mitochondrial perturbations; mitophagy deficiency (β-Clec16a^KO^), mtDNA depletion (β-Tfam^KO^), and fusion deficiency (β-Mfn1/2^DKO^). (B) STRING analysis of the 74 commonly upregulated and 28 commonly downregulated genes across all models. (C) Heatmap generated from bulk RNA sequencing of Ctrl and β-Clec16a^KO^ islets after RFD or HFD displaying ISR gene expression. *n* = 4-6 mice per group. (D) Heatmap generated from bulk RNA sequencing of Ctrl and β-Tfam^KO^ islets displaying ISR gene expression. *n* = 6-8 mice per group. (E) Representative WB and quantitation demonstrating expression of pEIF2α and Atf4 protein from Ctrl or β-Clec16a^KO^ islets fed RFD or HFD, Vinculin serves a loading control. *n* = 4-6/group. \*\**P* < 0.01 by one-way ANOVA, Tukey’s multiple comparison post-test. (F) Representative WB and quantitation demonstrating expression of pEIF2α and Atf4 protein from Ctrl or β-Tfam^KO^ islets, Vinculin serves a loading control. *n* = 3 group. \**P* < 0.05, \*\**P* < 0.01 by Student’s unpaired *t*-test. (G) *Bip* expression from bulk RNA sequencing of Ctrl or β-Clec16a^KO^ islets after RFD and HFD (left) and Ctrl and β-Tfam^KO^ islets (right). *n* = 4-8 mice per group. (H) qPCR analysis of *TFAM* and mtRNAs in shScramble or shTFAM-expressing EndoC-βH3 cells and human pseudoislets. *n* = 8/group. \**P* < 0.05, \*\**P* < 0.01, \*\*\**P* < 0.001 by Student’s unpaired *t*-test. (I) TFAM protein expression by immunostaining in shTFAM compared with shScramble human pseudoislets. C-peptide marks β cells. Scale bar = 50*μ*m, *n* = 4/group. (J) Expression of pEIF2α, total EIF2α and ATF4 protein by WB from shScramble or shTFAM EndoC-βH3 cells and human pseudoislets. Vinculin serves a loading control. *n* = 3-4/group. (K-L) qPCR analysis of ISR target genes and *BIP* expression (K) and β cell markers (L) in shScramble or shTFAM EndoC-βH3 cells and human pseudoislets. *n* = 5-8/group. \**P* < 0.05, \*\**P* < 0.01 by Student’s unpaired *t*-test.

While anterograde signaling from the nucleus to mitochondria has been described in β-cells (*40, 48*), mitochondrial signals that direct nuclear gene expression (*i.e.*, retrograde signaling) have not been observed in β-cells to date. A pathway observed to mediate mitochondrial retrograde signaling is the ISR, which coordinates mitochondrial or ER damage responses into a single pathway (*61*), triggered by phosphorylation of the eukaryotic translation initiation factor Eif2α, which then stabilizes Atf4 to promote downstream transcriptional responses (*62*). Indeed, we observed upregulation of canonical ISR transcriptional targets in RFD-fed β-Clec16a^KO^ mice, which increased to a greater degree upon HFD feeding (Fig. 2C). Canonical ISR transcriptional targets were similarly upregulated in β-Tfam^KO^ islets as well as in β-Mfn1/2^DKO^ islets (Fig. 2D and fig. S9A) Also overexpressed in these models was *Txnip*, which connects glucotoxicity with mitochondrial damage (*63*). Eif2α phosphorylation and Atf4 protein were also significantly increased in β-Clec16a^KO^, β-Tfam^KO^, and β-Mfn1/2^DKO^ islets (Fig. 2E-F and fig. S9B). Notably, we did not observe signs of ER stress, as we found no increases in expression of the ER chaperone *Bip* in β-Clec16a^KO^, β-Tfam^KO^, or β-Mfn1/2^DKO^ islets (Fig. 2G and fig. S9A), nor did we observe overt changes in ER morphology in RFD or HFD β-Clec16a^KO^ islets (fig. S10A-D) or β-Mfn1/2^DKO^ islets (*35*) by TEM.

To determine if impaired mitochondrial quality control elicits the ISR and loss of β-cell maturity in humans, we next generated TFAM-deficient EndoC-βH3 human β cells as well as primary human islets using the recently described pseudoislet approach (*64*). Briefly, primary human islets were dispersed, transduced with adenoviral particles encoding shRNA targeting TFAM (or scramble controls), and then re-aggregated into pseudoislets. We first confirmed highly efficient TFAM knockdown in EndoC-βH3 cells and human pseudoislets, and reductions in mtRNA expression following knockdown of TFAM, consistent with impairments in TFAM function (Fig. 2H-I). Similar to our observation in mouse models, loss of TFAM activated the mitochondrial ISR in human EndoC-βH3 cells and pseudoislets, evidenced by EIF2α phosphorylation and ATF4 protein stabilization, induction of ISR transcriptional targets *CEBPβ*, *CHOP*, *GDF15*, and *FGF21*, and no alterations in *BIP* expression (Fig. 2J-K). Further, we observed evidence of loss of β cell maturity following TFAM deficiency in shTFAM-treated human EndoC-βH3 cells and pseudoislets (Fig. 2L). Importantly, these results confirm that defects in mitochondrial quality control induce β cell immaturity and the ISR in both mouse and human islets.

### Examination of retrograde activating signals following defective mitochondrial quality control

We next assessed the upstream mitochondrial signals that promote the ISR following β-cell mitochondrial quality control defects. The mitochondrial ISR can be activated via excess mtROS, accumulation of misfolded mitochondrial proteins, or ETC/OXPHOS defects (*61*) (*65-67*). Thus, we first tested if excess mtROS led to β-cell dysfunction following mitophagy deficiency by intercrossing β-*Clec16a*^KO^ mice (or *Ins1*-Cre controls) with Cre-inducible mitochondrial-targeted catalase (mCAT) overexpressor mice to selectively scavenge β-cell mtROS (*68, 69*) (fig. S11A). However, mCAT overexpression, which reduces H_2_O_2_ as well as superoxide in β-cells (*68*) (*69*), did not improve glycemic control or insulin secretion in HFD-fed β-*Clec16a*^KO^ mice (fig. S11B-D). Next, we measured markers of the mitochondrial unfolded protein response (UPR^mt^) as a screen for transcriptional responses to misfolded mitochondrial proteins. However, we did not observe a consistent increase in markers of the UPR^mt^ across our models of impaired mitochondrial quality control (fig. S11E-G), suggesting that mitochondrial protein misfolding as well as mtROS are unlikely to be responsible here for induction of the ISR.

We next measured the ATP/ADP ratio with the Perceval biosensor in live β-cells as an initial assessment of energetics following loss of mitochondrial quality control (*70*). We observed reduced glucose-stimulated ATP/ADP ratio in islets from β-Clec16a^KO^ mice compared to controls on RFD, which worsened upon HFD-feeding (fig. S12A). These differences occurred without defects in total energetic reserve as measured after exposure to NaN_3_ to quench all mitochondrial function (fig. S12A). In islets from β-Tfam^KO^ mice, we observed stark reductions in both glucose-stimulated energetic output and total energetic reserve compared to controls, consistent with a substantial energetic deficit (fig. S12B).

Connections between the ETC/OXPHOS system and the ISR are well known yet can be challenging to resolve due to the multiple biochemical repercussions on the ETC/OXPHOS system that might be anticipated following loss of mitochondrial quality control. Indeed, we observed disruption of OXPHOS subunits at the protein level in our models of impaired mitochondrial quality control (fig. S2E, S3D, and S5H). To disentangle the multiple ETC/OXPHOS defects expected following mitochondrial genome instability, Mick, et al. previously induced the ISR via specific inhibition of complexes I, III, or V in myoblasts and differentiated myotubes, concluding that a complex network of metabolic defects downstream of energetic disruption activate the ISR (*71*). With these challenges in mind, we attempted several approaches to corroborate if damage to the ETC/OXPHOS system might contribute to ISR activation following defects in β cell mitochondrial quality control. To this end, we induced activation of the ISR following TFAM-deficiency in EndoC-βH3 human β cells after transduction with an adenovirus encoding an shRNA directed against TFAM (AdV-shTFAM). We then treated AdV-shTFAM (and AdV-shScramble) human β cells with the flavonoid (-)-Epicatechin, which has been previously demonstrated to improve respiratory function as well as OXPHOS expression and activity in β cells (*72-74*). Interestingly, we observed that EPI exposure attenuated activation of the ISR in AdV-shTFAM human β cells (fig. S12C-D).

Recent studies have shown that overexpression of bacterial or yeast proteins to facilitate continued NADH oxidation following loss of complex I *in vivo* or inhibition of complexes I, III, or V *in vitro* can attenuate ISR activation (*71, 75*). To evaluate if we would observe a similar phenomenon following mitochondrial genome instability, we overexpressed (1) LbNOX, a bacterial water-forming NADH oxidase which converts NADH to NAD+ independent of the ETC, (2) mitoLbNOX, a mitochondrial matrix-targeted LbNOX variant, or (3) NDI1, a single-subunit alternative internal NADH dehydrogenase that passes electrons to ubiquinone to facilitate continued NADH oxidation and ATP synthesis by OXPHOS, in EndoC-βH3 human β cells after transient knockdown of TFAM. While these approaches are distinct from the actions of EPI, we intriguingly observed a similar effect as EPI exposure, such that expression of LbNox, mitoLbNox or NDI1 diminished induction of the ISR in AdV-shTFAM-treated human β cells (fig. S12E). These complementary studies suggest that defects in the ETC/OXPHOS system following mitochondrial genome instability contribute to the induction of the ISR in human pancreatic β cells.

### Loss of mitochondrial quality control promotes β cell chromatin remodeling

To resolve changes in the transcriptional landscape of β cells following loss of mitochondrial quality control, we applied both single-nucleus RNAseq and ATACseq to assess chromatin remodeling that may be associated with mitochondrial retrograde signaling. UMAP plots of snRNAseq and snATACseq data revealed distinct populations of β cells of HFD-fed β-Clec16a^KO^ mice compared to controls (Fig. 3A). These results were supported by pathway analyses, which found significant differences between biological processes in β cells from control and β-Clec16a^KO^ mice (fig. S13A-B). Indeed, the snRNAseq results revealed enriched expression of targets associated with insulin secretion in control β cells when compared to Clec16a-deficient β cells (fig. S13A-B). Analysis of snATACseq data revealed 25,016 putative *cis-*regulatory elements uniquely detected as chromatin accessibility peaks unique to Clec16a-deficient β cells, with 3,567 peaks that were absent when compared to control β cells (Fig. 3B). These changes included loss of chromatin accessibility at multiple sites in the *Ins2* locus and accessibility gains in the *Atf3* and *Gdf15* loci in Clec16a-deficient β cells (Fig. 3C).

**Figure 3.**
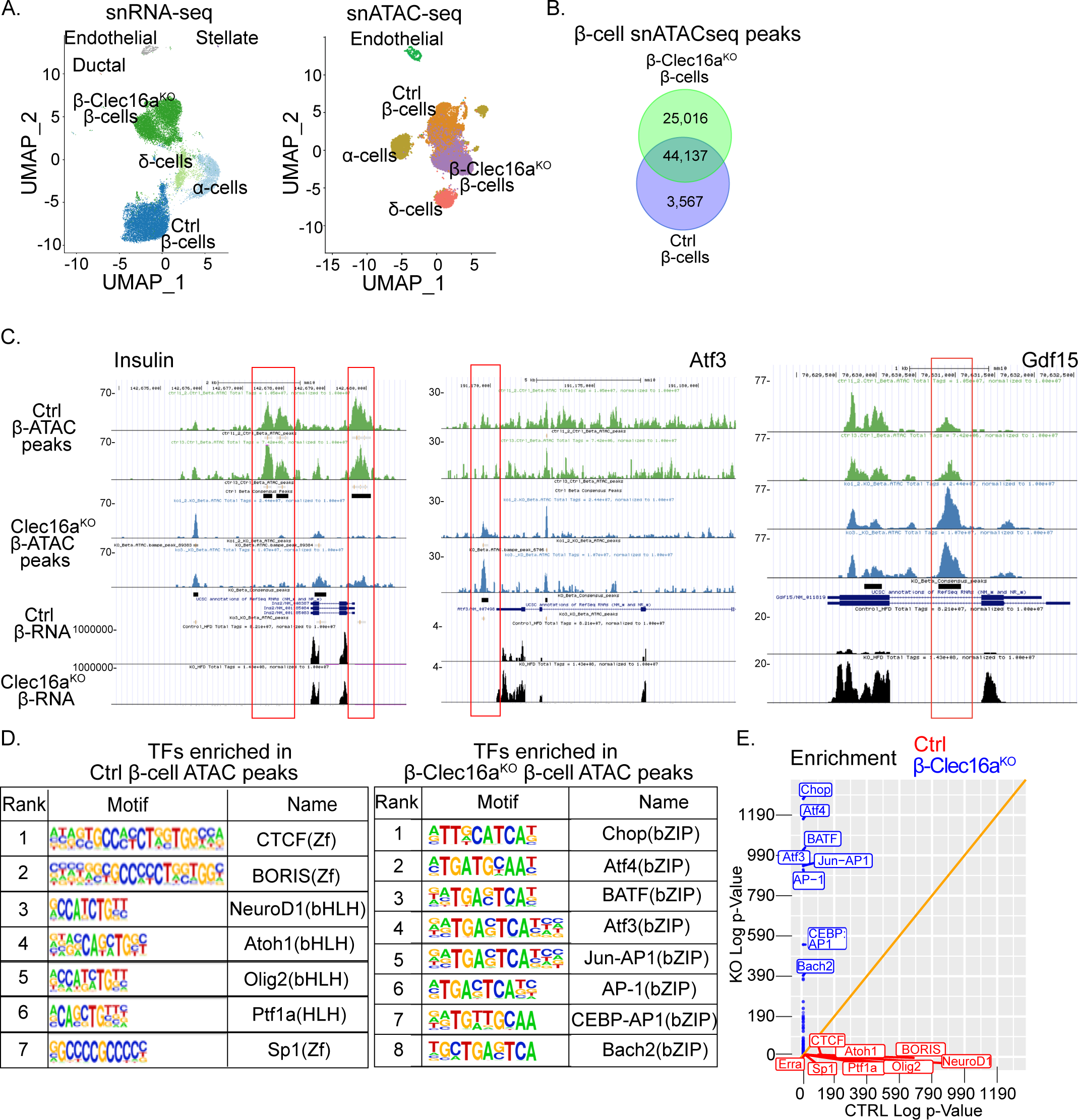
Loss of mitochondrial quality control induces chromatin remodeling. (A) UMAP plots of snRNA-seq from Ctrl and β-Clec16a^KO^ islet cell nuclei (left) and snATAC-seq (right) from Ctrl and β-Clec16a^KO^ islet cell nuclei. *n* = 4. (B) Venn diagram of common and unique open chromatin regions from snATAC-seq of Ctrl and β-Clec16a^KO^ β-cell nuclei. (C) Representative *Ins2, Atf3,* and *Gdf15* snATAC accessibility peaks from β-cells of HFD-fed Ctrl (1^st^ and 2^nd^ tracks shown in green from 2 independent mice) and β-Clec16a^KO^ (3^rd^ and 4^th^ tracks shown in blue from 2 independent mice) islets as well as bulk RNA expression-peaks from HFD-fed Ctrl and β-Clec16a^KO^ islets (5^th^ and 6^th^ tracks, respectively, shown in black). (D) HOMER analysis of Ctrl β-Cell ATAC peaks showing enrichment of motifs corresponding to zinc finger (Zf) and beta-Helix Loop Helix (bHLH) identity factors (left) and β-Clec16a^KO^ β-Cell ATAC peaks showing enrichment of motifs corresponding to bZIP stress response factors (right). *n* = 4. (E) Visualization of the top 8 enriched TF motifs from Ctrl β-cell specific ATAC peaks (red) and β-Clec16a^KO^ β-cell specific ATAC peaks (blue). *n* = 4.

Enrichment analysis of differentially accessible peaks in *Clec16a*-deficient β cells using the Genomic Regions Enrichment of Annotations Tool (GREAT; (*76*)) revealed the loss of peaks in regions related to the regulation of mitophagy, lysosomal transport, and glucose metabolic processes, consistent with known functions for Clec16a ((*39, 77*); fig. S13C). In contrast, GREAT peaks uniquely accessible in *Clec16a*-deficient β cells were enriched in regions related to cellular responses to carbohydrate stimulus, abnormal pancreas development, and abnormal pancreatic β cell differentiation (fig. S13D). Importantly, we observed significant enrichment of transcription factor binding site motifs for key ISR transcription factors in Clec16a-deficient β cells by HOMER (*78*), with bZIP factors, including Atf4, Atf3, and CHOP, the most enriched in β-Clec16a^KO^ mice (Fig. 3D-E). Increased binding sites for Bach2, a transcription factor which is associated with β cell immaturity in T2D (*79*), were also found in Clec16a-deficient β cells (Fig. 3D-E). We additionally observed loss of binding sites for transcription factors known to be vital for β cell maturity, identity, and function in Clec16a-deficient β cells, most notably the critical β cell transcription factor and maturity regulator NeuroD1, as well as more broadly expressed transcription factors Sp1 and CTCF (Fig. 3D-E; (*80-82*)). These results suggest that mitochondrial retrograde signals are associated with modification of chromatin accessibility favoring transcriptional activation of the ISR and loss of cell identity and maturity.

### Loss of mitochondrial quality control induces immaturity and the ISR in hepatocytes and brown adipocytes

Given the importance of mitochondrial damage in metabolic disorders, including T2D, MASLD, and MASH, we next asked if defects in mitochondrial quality control induce comparable impairments in cell maturity in other metabolic tissues. To this end, we generated mice bearing liver-specific deletion of Mfn1/2 or Tfam, by intraperitoneal administration of adeno-associated virus 8 expressing Cre-recombinase under control of the hepatocyte-specific *Tbg* promoter (AAV8-Tbg^cre^) to 12-week-old HFD-fed control mice, and *Mfn1/2*- or *Tfam*-floxed animals (hereafter known as L-Mfn1/2^DKO^ or L-Tfam^KO^ mice, respectively; Fig. 4A-C). In both models, we observed expected decreases in mtDNA content and alterations of expression of OXPHOS subunits, including reduced Ndufb8 and mt-Co1 and increased Sdhb and Atp5a, without overt reductions in the mitochondrial mass marker Tom20 (Fig. 4D-G). L-Mfn1/2^DKO^ animals developed alterations in glucose tolerance and insulin sensitivity, whereas L-Tfam^KO^ mice developed slight alterations in insulin sensitivity with notable histological defects in liver morphology, including hepatic steatosis and ballooning (fig. S14A-E). However, we did not observe signs of liver fibrosis in L-Mfn1/2^DKO^ and L-Tfam^KO^ mice (fig. S14F). Signs of hepatic dysfunction were also evident following loss of mitochondrial quality control, with significant increases in serum ALT and LDH in L-Tfam^KO^ mice and decreases in serum cholesterol in L-Mfn1/2^DKO^ mice, and decreased liver glycogen in both L-Mfn1/2^DKO^ and L-Tfam^KO^ mice (Fig. 4H-K). Consistent with our observations in β cells, loss of mitochondrial quality control also did not induce apoptosis in hepatocytes (fig. S14G).

**Figure 4.**
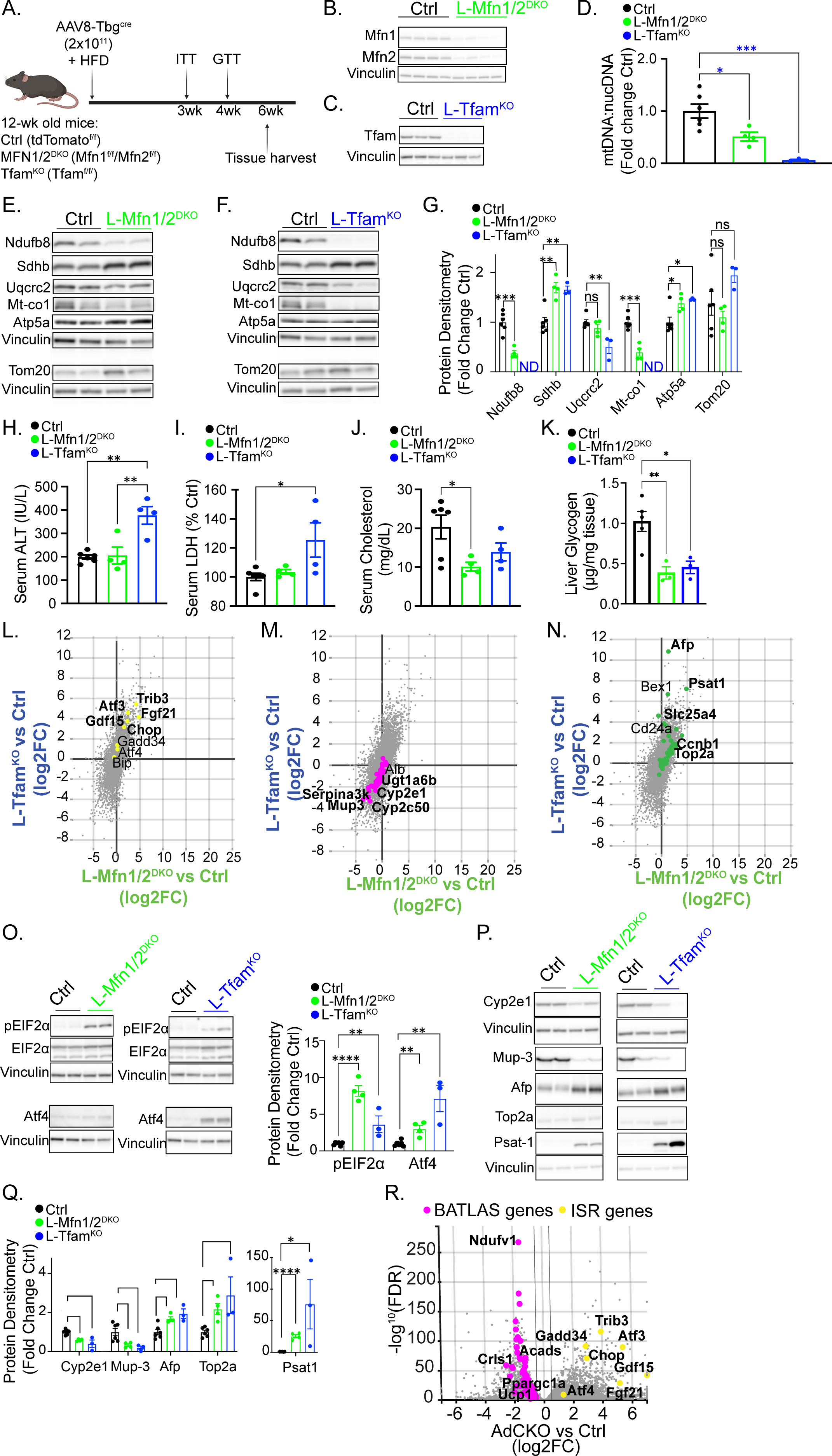
Impaired mitochondrial quality control commonly induces the ISR and immaturity across several metabolic tissues. (A) Schematic diagram illustrating the experimental design of AAV8-Tbg^cre^ delivery and analysis of liver-specific loss of mitochondrial fusion (L-Mfn1/2^DKO^) and mtDNA copy number control (L-Tfam^KO^). (B) Protein expression of Mfn1 and Mfn2 by WB in liver from Ctrl (AAV8-Tbg^cre^) and L-Mfn1/2^DKO^ mice. (C) Protein expression of Tfam by WB in liver from Ctrl (AAV8-Tbg^cre^) and L-Tfam^KO^ mice. (D) Quantification of mtDNA to nuclear DNA ratio in liver by qPCR. *n* = 4-6 animals per group. \**P* < 0.05, \*\*\**P* < 0.001 by one-way ANOVA, Tukey’s multiple comparison post-test. (E-F) Protein expression of murine OXPHOS complex subunits and Tom20 in liver from Ctrl and L-Mfn1/2^DKO^ (E) and Ctrl and L-Tfam^KO^ (F) mice by WB. Vinculin serves as a loading control. *n* = 3-6 mice per group. (G) Densitometry quantification of OXPHOS subunits and Tom20 from (E-F). *n* = 3-6 mice per group. \**P* < 0.05, \*\**P* < 0.01, \*\*\**P* < 0.001 by Student’s unpaired *t*-test. ND=not detected. (H-K) Liver function analyses from AAV8-Tbg^cre^ Ctrl, L-Mfn1/2^DKO^ and L-Tfam^KO^. (H) Serum ALT; \*\**P* < 0.01 one-way ANOVA, Tukey’s multiple comparison post-test. (I) Circulating serum LDH levels; \**P* < 0.05 one-way ANOVA, Tukey’s multiple comparison post-test. (J) Circulating serum cholesterol levels; \**P* < 0.05 one-way ANOVA, Tukey’s multiple comparison post-test. (K) Liver glycogen levels; \**P* < 0.05, \*\**P* < 0.01 one-way ANOVA, Tukey’s multiple comparison post-test. (L-N) Differential gene expression comparison plots of Ctrl vs either L-Tfam^KO^ (y-axis) or L-Mfn1/2^DKO^ (x-axis) showing similarly upregulated ISR genes (L), immaturity genes (*83*) (M), and similarly downregulated maturity genes (*83*) (N). Select gene expression changes reaching statistical significance in both L-Mfn1/2^DKO^ and L-Tfam^KO^ mice are noted in bold font. (O) Protein expression (left) and densitometry (right) of pEIF2α, EIF2α, or Atf4 in Ctrl, L-Mfn1/2^DKO^ and L-Tfam^KO^ liver. Vinculin serves as a loading control. *n* = 3-6 mice per group. \*\**P* < 0.01, \*\*\*\**P* < 0.0001 by Student’s unpaired *t*-test. (P) Protein expression of mature (Mup-3, Cyp2e1) and immature (Top2a, AFP and Psat-1) hepatocyte markers from liver of Ctrl, L-Mfn1/2^DKO^ and L-Tfam^KO^ mice. Vinculin serves as a loading control. (Q) Densitometry of immature and mature protein levels shown in (P). *n* = 3-6 mice per group. \**P* < 0.05, \*\**P* < 0.01, \*\*\**P* < 0.001, \*\*\*\**P* < 0.0001 by Student’s unpaired *t*-test. (R) Volcano plot of differential gene expression in Ctrl vs AdCKO (adiponectin Cre-driven Crls1 knock-out) brown fat showing upregulated ISR genes (yellow circles) and downregulated BATLAS brown fat maturity genes (pink circles; generated from (*89*)).

We next used RNAseq to begin to determine if loss of mitochondrial quality control promoted a common signature of activation of the mitochondrial ISR, induction of cell immaturity, and loss of terminal cell identity markers in hepatocytes. As observed in β cells, L-Mfn1/2^DKO^ or L-Tfam^KO^ mice had significantly increased expression of canonical ISR targets without upregulation of *Bip* expression (Fig. 4L). To determine if defects in mitochondrial quality control altered hepatocyte maturity, we overlaid the RNAseq data with a curated list of key hepatocyte terminal identity and developmental markers (*83*). We found decreased expression of terminal hepatocyte markers including genes associated with cytochrome P450 activity and bilirubin glucuronidation, and increases in several markers of hepatocyte immaturity, including alpha fetoprotein (*Afp*), *Top2a*, and *Psat1* ((*83*); Fig. 4M-N). Moreover, western blot analysis confirmed induction of the ISR by detection of increased Atf4 protein levels and Eif2α phosphorylation (Fig. 4O), reduced levels of mature hepatocyte proteins (Cyp2e1, Mup-3), and increased levels of markers of hepatocyte immaturity (Afp, Top2a, Psat-1) in both L-Mfn1/2^DKO^ and L-Tfam^KO^ mice (Fig. 4P-Q).

Due to the broad importance of mitochondria for brown adipocyte tissue function (*84*) (*85*), we next surveyed expression of terminal identity markers and the ISR from mice bearing adipocyte-specific deletion of cardiolipin synthase (Crls1), an enzyme essential for biosynthesis of the key mitochondrial phospholipid cardiolipin (*86*). Cardiolipin plays several crucial roles in mitochondria, including regulation of mitochondrial quality control through cardiolipin-dependent mitophagy (*87, 88*). To this end, we analyzed publicly available RNAseq data from BAT depots isolated from mice bearing adipocyte-specific deletion of *Crls1* generated by intercrossing *Crls1* floxed animals with the adiponectin Cre-recombinase strain (hereafter known as AdCKO mice; (*86*)). AdCKO mice were previously reported to develop reduced uncoupled mitochondrial respiration, glucose uptake, and thermogenesis in BAT depots, however, a link between cardiolipin metabolism and BAT maturity was not assessed (*86*). To address this question, we evaluated differentially expressed genes between AdCKO and control BAT depots for targets from the BATLAS database, which includes numerous key terminal BAT identity/maturity markers (*89*). We found a striking downregulation of numerous BATLAS genes in AdCKO BAT depots, including a loss of BAT-specific genes associated with cellular maturity/identity, such as *Ppargc1α* and *Ucp1* (Fig. 4R). Further, we observed an induction of canonical ISR transcriptional targets in AdCKO BAT depots (Fig. 4R). Taken together, these results highlight a vital role for mitochondrial quality control in promoting terminal cell identity by restraining mitochondrial retrograde signaling in numerous metabolic tissues.

### Blockade of retrograde signaling restores β-cell mass and maturity *in vivo*

Our results in multiple metabolic tissues suggest that defects in mitochondrial quality control engage the ISR and induces loss of terminal cell maturity/identity. We, therefore, hypothesized that induction of retrograde mitochondrial signaling through the ISR directly leads to loss of cellular maturity. To test the importance of the ISR in β-cell maturity, we treated HFD-fed β-Clec16a^KO^ mice with ISRIB, a well-established pharmacologic inhibitor of the ISR (*90*). We first treated isolated islets from HFD-fed β-Clec16a^KO^ mice and controls with ISRIB for 24 hours, observing the expected reduction of ISR transcriptional targets, including *Cebpβ* and *Trib3*, in islets from HFD-fed β-Clec16a^KO^ mice (Fig. 5A-B). Concordantly, we found that ISRIB restored expression of the terminal β-cell identity marker *Ucn3* and reduced expression of the dedifferentiation marker Aldh1a3 in islets from HFD-fed β-Clec16a^KO^ mice (Fig. 5C-D). To determine if the restoration of β-cell identity was related to an unanticipated upregulation of mitophagy in Clec16a-deficient β-cells, we intercrossed β-Clec16a^KO^ mice with the mt-Keima mitophagy reporter strain to assess mitophagy flux. Importantly, ISRIB did not ameliorate the well-known defects in mitophagy flux associated with Clec16a-deficiency (Fig. 5E), suggesting the restoration of β-cell identity markers by ISRIB was not due to improved mitophagy but rather related to blockade of retrograde signaling through the ISR.

**Figure 5.**
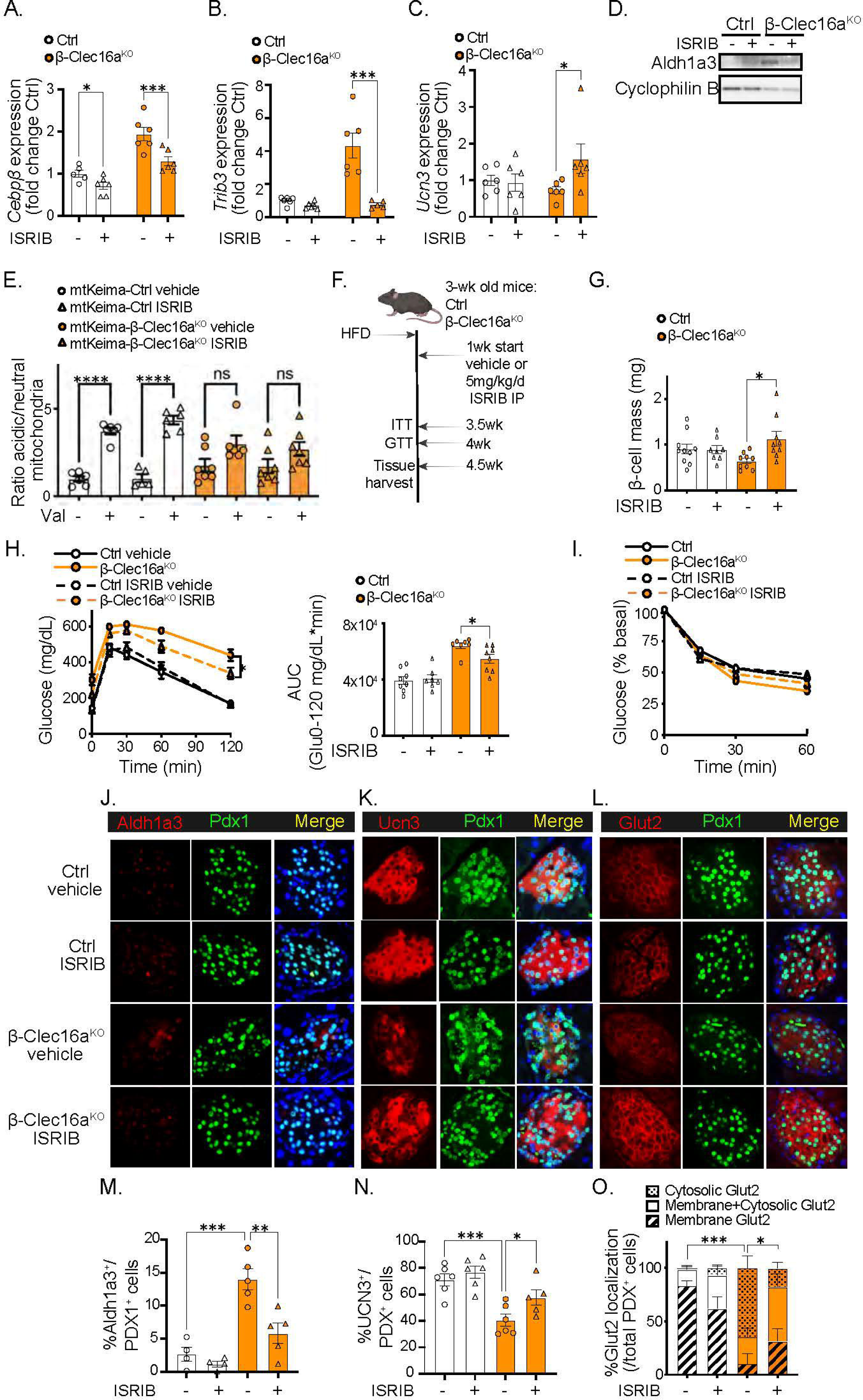
Pharmacologic blockade of mitochondrial retrograde signaling *in vivo* restores β-cell mass and maturity. (A-C) qPCR of ISR targets *Cebpβ* and *Trib3* (A-B) and β-cell maturity marker Ucn3 (C) from Ctrl or β-Clec16a^KO^ islets treated *ex vivo* with vehicle or ISRIB. *n* = 6/mice per group. \**P* < 0.05, \*\*\**P* < 0.01 by two-way ANOVA, Sidak’s multiple comparison posttest effect of drug within genotype. (D) Protein expression of Aldh1a3 by WB from Ctrl or β-Clec16a^KO^ islets treated *ex vivo* with ISRIB. Cyclophilin B was used as a loading control. (E) Ratio of acidic to neutral compartment localized mitochondria from β-cells from mtKeima-Ctrl or mtKeima-β-Clec16a^KO^ mice treated with vehicle or ISRIB *ex vivo* for 24hr. *n* = 5-7mice per group. \*\*\*\**P* < 0.0001 by one-way ANOVA, Tukey’s multiple comparison post-test. (F) Schematic diagram illustrating the experimental design of *in vivo* ISRIB treatment and analysis of Ctrl or β-Clec16a^KO^ mice. (G-I) Results from Ctrl or β-Clec16a^KO^ mice treated with vehicle or ISRIB once daily for 4-weeks. *n* = 8-10 mice per group. (G) Pancreatic β-cell mass; \**P* < 0.05 by two-way ANOVA effect of ISRIB within genotype. (H) Blood glucose concentrations (left) and AUC (right) during an IPGTT. \**P* < 0.05 by Student’s unpaired *t*-test. (I) Blood glucose levels during an ITT shown as % basal glucose. (J-L) Representative pancreatic immunofluorescence images from Ctrl or β-Clec16a^KO^ mice treated *in vivo* with vehicle or ISRIB. (J) Sections stained with antibodies against Aldh1a3 (red), Pdx1 (green), and DAPI (blue). (K) Sections stained with antibodies against Ucn3 (red), Pdx1 (green), and DAPI (blue). (L) Sections stained with antibodies against Glut2 (red), Pdx1 (green), and DAPI (blue). (M-O) Quantification of staining for (J) % cytosolic Aldh1a3^+^ / Pdx1^+^ cells. *n* = 4-5 mice per group. \*\**P* < 0.01 by one-way ANOVA, Tukey’s multiple comparison post-test; (K) % Ucn3^+^/Pdx1^+^ cells. *n* = 5-6 mice/group. \**P* < 0.05, \*\*\**P* < 0.01 by one-way ANOVA, Tukey’s multiple comparison post-test; and (O) Glut2 localization classified as all cytosolic (Cytosolic), mixed cytosolic and plasma membrane (Membrane+Cytosolic) and all plasma membrane (Membrane) as a % of all Pdx1^+^ cells. *n* = 6 mice/group. \**P* < 0.05, \*\*\**P* < 0.01 by one-way ANOVA, Tukey’s multiple comparison post-test.

To next determine if retrograde signaling through the ISR leads to loss of β-cell mass and identity *in vivo*, we treated HFD-fed β-Clec16a^KO^ mice and littermate controls with ISRIB by daily intraperitoneal administration for 4 weeks (Fig. 5F). Notably, extended treatments with ISRIB in control animals *in vivo* did not impair β-cell mass or glucose tolerance (Fig. 5G-I), consistent with a previous report (*91*). Further, we performed hyperinsulinemic-euglycemic clamp studies in ISRIB treated mice, which revealed no significant changes in glucose infusion rate, tissue glucose uptake, or glucose clearance, no significant changes in basal insulin levels and insulin clearance, and no significant differences in hepatic glucose production (fig. S15A-I), suggesting that ISRIB treatment did not affect peripheral insulin sensitivity.

ISR inhibition rescued the loss of β-cell mass in HFD-fed β-Clec16a^KO^ mice (Fig. 5G). ISRIB treatment also ameliorated glucose intolerance in HFD-fed β-Clec16a^KO^ mice, while insulin tolerance tests remained unchanged between groups (Fig. 5H-I). ISRIB treatment reduced expression of Aldh1a3 in HFD-fed β-Clec16a^KO^ mice and increased the frequency of Ucn3-positive β cells bearing proper expression/localization of Glut2, consistent with restoration in β cell maturity and identity (Fig. 5J-O). Notably, the benefits of ISRIB on β-cell mass and identity are unlikely to be due to an indirect mitigation of chronic hyperglycemia, as treatment with the SGLT1/2 inhibitor phlorizin to improve glucose homeostasis by increased glycosuria did not rescue β-cell mass or function in mitophagy-deficient mice (fig. S16A-E). Taken altogether, these studies position the induction of mitochondrial retrograde signaling through the ISR as a direct regulator of cell maturity and identity in metabolic tissues.

## DISCUSSION

Mitochondria are pivotal cellular organelles, which not only provide the energy required for cellular function, growth, division, and survival, but also carry vast complexity, importance, and reach beyond energy production (*92*). Here, we demonstrate that loss of mitochondrial quality control engages a retrograde signaling response shared across metabolic tissues, which ultimately impairs cellular identity and maturity. Perturbations at any point within the mitochondrial quality control machinery converge to activate the ISR, eliciting a transcriptional signature associated with induction of cellular immaturity and modulation of chromatin architecture. Intriguingly, changes in terminal cell identity are reversible, as inhibition of the mitochondrial ISR restores cellular maturity, mass, and function. Together, our results demonstrate that mitochondrial quality control plays a central role in the maintenance of cell identity and maturity in metabolic tissues.

Our results were surprising as mitochondrial defects have been classically associated with a decline in proliferation or survival. Retrograde signals generated from the mitochondria in yeast to activate the transcriptional regulators Rtg1, 2, and 3 have been well characterized (*93-95*), yet mammalian orthologs of these factors have proved elusive to date. Indeed, retrograde signaling has unclear roles in the regulation of mammalian cellular homeostasis (*61, 96, 97*). Interestingly, the Rtg family has been implicated in the modulation of TCA cycle enzymes, glutamate biosynthesis and mtDNA maintenance, but not in cellular identity (*95*). Intriguingly, our studies position an unexpected role for mitochondrial quality control defects to impair cellular maturity and identity via retrograde signaling, ultimately leading to the deterioration of metabolic tissues.

Our results identifying the crucial importance for mitochondria to direct cellular maturity and identity additionally resolve confusion regarding anterograde (*i.e.*, nuclear to mitochondrial) control of cell fate across metabolic tissues. Instances of anterograde control of cellular maturity and identity have been reported; however these transcriptional regulators, including Foxo1, Prdm16, Pgc1α, ERRγ, and Pdx1, possess unique tissue-specific effects and have not been demonstrated to directly regulate both cellular maturity and mitochondrial metabolism uniformly across metabolic tissues (*40, 98-102*). For instance, Foxo1 deficiency promotes β cell dedifferentiation via control of the NADH-dependent oxidoreductase Cyb5r3, which localizes to membranes in the ER, mitochondria, and plasma membrane (*26, 103, 104*). In contrast, Foxo1 deficiency promotes BAT differentiation and has not been shown to lead to dedifferentiation in adult hepatocytes (*105-107*). Further, Pgc1α, while crucial for BAT maturity and mitochondrial function, is dispensable for β cell maturity and mitochondrial function (*99, 108-110*). ERRγ induces postnatal maturation of β cells and BAT while activating expression of numerous mitochondrial genes; however, ERRγ deficiency has not been previously implicated in the development of hepatocyte immaturity via mitochondrial alterations (*102, 111, 112*). Due to their tissue-specific effects, many of these transcriptional regulators regulate additional mitochondria-independent processes, rendering it challenging to conclude that regulation of mitochondrial function by these nuclear factors is solely sufficient to maintain cellular maturity and identity across metabolic tissues. Thus, our studies obviate these concerns by establishing the central importance of mitochondrial health to maintain cellular maturity and identity across metabolic tissues.

We observe changes in chromatin accessibility following the induction of mitochondrial quality control defects and activation of the ISR. The increased frequency of binding motifs related to the central ISR transcriptional regulator ATF4, as well as ATF4 targets, is suggestive that retrograde signaling might have the potential to alter chromatin structure. Indeed, ATF4 is a pioneer factor capable of altering chromatin accessibility (*113, 114*), which could be consistent with the transcriptional signatures observed in metabolic tissues with mitochondrial damage. However, it is unclear how the induction of ATF4 remodels sites of chromatin accessibility that are lost at key markers of β cell maturity, such as NeuroD1 (*80*). Loss of binding regions essential for β cell maturity following mitochondrial defects in quality control could also be secondary to impaired substrate availability and metabolism, eventually affecting epigenetic modifications, similar to metabolic control of epigenetics reported in the cancer biology literature (*115*). Thus, we speculate that defects in bioenergetic capacity following mitochondrial quality control defects could have a cascade of effects, initiated by activation of the ISR, to remodel chromatin architecture, and drive dedifferentiation to an immature state. How retrograde metabolic signals may be tuned by sensors of the ISR, including the kinases PERK, GCN2, HRI, and PKR (*116*), and their subsequent effects on chromatin accessibility, will be intriguing avenues for future study.

Mitochondria are broadly vital to the function of metabolic tissues, yet mitochondria are frequently viewed as client organelles rather than as primary drivers of cell fate. Indeed, the appearance of decreased mtDNA content, altered mitochondrial architecture, or mitophagy in metabolic tissues during T2D is often considered an indirect consequence of other etiologies, including transcriptional defects, ER stress, or nutritional excess (*117-119*). Relatedly, reduced ATP production and increased mtROS in hepatocytes triggering inflammation and fibrosis are well established findings in MASLD and MASH (*1, 120-122*). Moreover, hepatic mitochondrial morphological defects and impaired mitophagy have been reported (*15)*. Here, we observed key defects across the spectrum of the mitochondrial quality control pathway in human β-cells during T2D that are each sufficient to elicit β-cell immaturity in isolation in genetic mouse models or primary human islets. The remarkable consistency of these connections between mitochondrial quality control defects and cellular immaturity across human and mouse metabolic tissues further supports the key importance of mitochondria in maintaining terminal cell identity. Interestingly, our results suggest the loss of mitochondrial quality control across metabolic tissues lead to distinct effects from other cell types, such as renal epithelial cells, which develop activation of the cytosolic cGAS-stimulator of interferon genes (STING) DNA sensing pathway, cytokine expression, and immune cell recruitment upon Tfam-deficiency (*123*). We did not observe evidence for activation of cGAS-STING or cytokine expression across our models of mitochondrial mtDNA loss, including both β-Tfam^KO^ and L-Tfam^KO^ mice, suggesting the development of immaturity upon decreased mitochondrial quality control pathways may be distinct in metabolic tissues and unlikely to involve a type I interferon response. Additionally, mitophagy-deficiency in lung alveolar cells induces pulmonary fibrosis with aging (*124*); however, we did not observe increases in liver fibrosis or pro-fibrotic or pro-inflammatory markers (including *Fgf2*, *Il6*, or *Il10*) in our hepatocyte or β-cell models of impaired mitochondrial quality control, respectively. Studying these cell type-specific responses will be a captivating topic for future investigation.

Our data suggest that blockade of retrograde signaling can prevent the loss of β-cell mass and glycemic control following mitochondrial quality control defects. Our findings reinforce the importance of signatures of mitochondrial gene expression and OXPHOS defects in human β-cells of pre-diabetic donors with impaired glucose tolerance, potentially indicating that β-cell mitochondrial bioenergetic or quality control defects preceding disease may be an etiopathological contributor to T2D that possibly extends to other tissues. While the primary focus of this work was to define a central mechanism underlying mitochondrial quality control regulation of cell fate in metabolic tissues, future studies will be of keen interest to explore if ISRIB treatment can restore β-cell maturation and glycemic control in T2D as well as in T1D, where blockade of β-cell stress responses are capable of diabetes prevention (*125*) (*126*). Relatedly, despite reduced *Clec16a* expression in islets of donors with T2D and control by the T2D gene *Pdx1* (*40-42*), *Clec16a* is more commonly known as a gene associated with T1D in humans (*39, 127*). We previously observed the importance of Clec16a-mediated mitophagy in both rodent and human β-cells for the maintenance of insulin secretion and glucose homeostasis (*36, 39, 77, 128*). A limitation of our study was that it is unclear how Clec16a functions in human β-cells specifically in T2D. Thus, studies to clarify the function of Clec16a, mitophagy, and the ISR within human β-cells across the spectrum of T1D and T2D will be insightful in the future. Finally, it will be fascinating to determine if preemptively targeting mitochondrial quality control or retrograde signaling in specific metabolic tissues could prevent the onset of T2D, MASLD, or MASH.

## RESOURCE AVAILABILITY

### Lead Contact

Further information and request for resources and reagents should be directed to and will be fulfilled by the lead contact, Scott Soleimanpour

### Materials Availability

Rabbit polyclonal Clec16a antisera is available upon request from the lead author. mtKeima mice were maintained on the C57BL/6N background and intercrossed to Clec16a^f/f^ mice to generate mtKeima-Ctrl and mtKeima-β-Clec16a^KO^ mice; these mice are available on request from the lead author.

### Data Availability

Bulk RNA-seq, single nuclei RNA-seq and single nuclei ATAC-seq data have been deposited at GEO (GSE262023) and are publicly available at the date of publication. This paper also analyzed existing publicly available data. Microscopy data will be shared by lead contact on request.

Any additional information required to reanalyze the data reported in this paper is available from the lead author on request.

## Acknowledgements

S.A.S. acknowledges support from the JDRF (CDA-2016-189, COE-2019-861, SRA-2023-1392), the NIH (R01 DK108921, R01 DK135032, R01 DK135268, R01 DK136671, R01 DK127270, U01 DK127747, P30 DK020572), the Department of Veterans Affairs (I01 BX004444), the Brehm family, and the Anthony family. G.L.P. was supported by the American Diabetes Association (19-PDF-063). E.M.W. was supported by the NIH (5K01DK133533). B.A.K. was supported by the Department of Veterans Affairs (I01 BX004444). L.S.S. was supported by the NIH (R01 DK46409). The JDRF Career Development Award to S.A.S. is partly supported by the Danish Diabetes Academy and the Novo Nordisk Foundation. The Animal Metabolic, Physiological and Behavioral Phenotyping Core through the Michigan NIH Mouse Metabolic Phenotyping Center (MMPC-Live) performed clamp studies (U2CDK135066). We acknowledge the Microscopy, Imaging and Cellular Physiology Core and Islet Core of the University of Michigan DRC (P30 DK020572) for assistance with imaging studies and pseudoislet studies, respectively. We thank the University of Michigan Flow Cytometry Core for assistance with flow cytometry studies. Next generation sequencing was carried out in the Advanced Genomics Core at the University of Michigan. Human pancreatic islets and/or other resources were provided by the NIDDK-funded Integrated Islet Distribution Program (IIDP) (RRID:SCR _014387) at City of Hope, NIH Grant # U24DK098085. Human islets were also provided by the Alberta Diabetes Institute IsletCore at the University of Alberta in Edmonton (http://www.bcell.org/adi-isletcore.html) with the assistance of the Human Organ Procurement and Exchange (HOPE) program, Trillium Gift of Life Network (TGLN), and other Canadian organ procurement organizations. Islet isolation was approved by the Human Research Ethics Board at the University of Alberta (Pro00013094). All donors’ families gave informed consent for the use of pancreatic tissue in research. We thank Dr. K. Claiborn and members of the Soleimanpour laboratory for helpful advice.

## Author Contributions

G.L.P. and E.M.W. conceived, designed, and performed experiments, interpreted results, drafted and reviewed the manuscript. N.L. designed and performed experiments, interpreted results, and reviewed the manuscript. A.L., V.S., J.Z., T.S., E.C.R., A.M.S., E.L-D., M.B.P., D.L.H., A.R., K.M., V.S.P., I.X.Z., B.T., D.Z., S.A.W., and L.H. designed and performed experiments and interpreted results. S.C.J.P., P.A., L.Y., B.A.K., L.S.S., and L.S. designed studies, interpreted results, and reviewed the manuscript. M.L.S. designed studies, interpreted results, edited, and reviewed the manuscript. S.A.S. conceived and designed the studies, interpreted results, drafted, edited, and reviewed the manuscript.

