## Supplementary material for "Retrograde mitochondrial signaling governs the identity and maturity of metabolic tissues"

**Supplementary Materials**

Materials and Methods

Figs. S1 to S17

Tables S1 to S6

References (1-27)

### Materials and Methods

**Genetically modified mouse lines*:*** All mice were maintained in accordance with the University of Michigan's Institutional Animal Care and Use Committee under specific pathogen-free conditions. Up to 5 mice were housed per cage and were maintained on regular chow or high fat diet chow with *ad libitum* access to food on a 12 h light-dark cycle. Floxed *Tfam* (*Tfam*$^{f/f}$ mice (Jackson Laboratories, Stock no. 026123)), *Clec16a* (*Clec16a*$^{f/f}$ mice(*1*)), *Mfn1*/*Mfn2* (*Mfn1*$^{f/f}$/*Mfn2*$^{f/f}$ mice (Jackson Laboratories stock no. 026401 and 026525), Rosa26 lox-STOP-lox tdTomato reporter mice (Jackson Laboratories, Stock no. 007914), and floxed stop mCAT overexpression mice (Jackson Laboratories, Stock no. 030712) were used. All animals were maintained on a 100% C57BL/6N background. To generate β-cell-specific deletion, floxed models were crossed with *Ins1*-Cre mice from Jackson laboratories (JAX Stock No. 026801). *Ins1*-Cre–alone and respective floxed-only controls for each study (*Tfam*$^{f/f}$, *Mfn1*$^{f/f}$/*Mfn2*$^{f/f}$ or *Clec16a*$^{f/f}$ mice) were phenotypically indistinguishable from each other and combined as controls (Ctrl) as noted, with the exception of lineage tracing studies where controls were *Ins1*-Cre only. *Ins1*-Cre–alone were also phenotypically indistinguishable from wild-type C57BL/6N controls, consistent with previous reports from our group and others (*2-5*). For inducible β-cell deletion, *MIP1*-CreERT mice (JAX 024709) were used or crossed to *Clec16a*$^{f/f}$ mice to generate iβ-Clec16a$^{KO}$ mice as previously described (*6*). To induce knockout in this model, 7 wk old mice were injected intraperitoneally with 2 mg of tamoxifen (Cayman Chemicals) in corn oil every other day for 5 days. MIP1-CreERT alone mice injected with tamoxifen were used as controls for these studies.

For liver specific deletion, tdTomato reporter, *Tfam*$^{f/f}$, and *Mfn1*$^{f/f}$/*Mfn2*$^{f/f}$ mice at 12 wk of age were injected intraperitoneally with 2 x $10^{11}$ particles of AAV-Tbg$^{cre}$ (AAV8 serotype; Vector Biolabs; San Francisco, CA, USA) and placed on high fat diet for 4-6 wk. Mixed sex cohorts were used throughout the study. Mice were between 7 weeks and 1 year of age at the time of study, depending on endpoint (information is provided in the text). Mice were randomized into treatment or vehicle control groups and/or regular fat or high fat diet groups, as necessary.

**Human islet samples:** All human samples were procured from de-identified donors with or without diabetes from the Integrated Islet Distribution Program, Alberta IsletCore, or Prodo Laboratories and approved by the University of Michigan Institutional Review Board. Human primary islets were cultured at 37°C with 5% $CO_2$ in PIM(S) media (Prodo Laboratories, Aliso Viejo, CA, USA) supplemented with 10% FBS, 100 U/mL penicillin/streptomycin, 100 U/mL antibiotic/antimycotic, and 1mM PIM(G) (Prodo Laboratories, Aliso Viejo, CA, USA). Islets were used from male and female donors, and donor information is provided in table S2. Comparison of islet donors for mtDNA analyses and medication history for human islet donors with T2D is described in Tables S3 and S4.

**Mouse primary islet cultures:** Mouse primary islets were isolated by perfusing pancreata with a 1 mg/mL solution of Collagenase P (Millipore Sigma; St Louis, MO, USA) in 1 X HBSS into the pancreatic duct. Following excision of the pancreas, pancreata were incubated at 37°C for 13 min, and Collagenase P was deactivated by addition of 1XHBSS + 10% adult bovine serum (Quench buffer). Pancreata were dissociated mechanically by vigorous shaking for 30 sec, the resulting cell suspension was passed through a 70 μM cell strainer (Fisher Scientific, Waltham, MA, USA). Cells were centrifuged at 1000 rpm for 2 min, the pellet was resuspended in 20 mL Quench buffer and gently vortexed to thoroughly mix. Cells were again centrifuged at 1000 rpm, 1 min. This wash step was repeated once more. Following washes, the cell pellet was resuspended in 5 mL

Histopaque (Millipore-Sigma; St Louis, MO, USA) with gentle vortexing. An additional 5 mL Histopaque was layered on the cell suspension, and finally 10 mL Quench buffer was gently layered on top. The cells were spun at 900 x *g* for 30 min at 10°C, with the brake off. The entire Histopaque gradient was pipetted off and passed through an inverted 70 μM filter to trap the islets cells. Islets were washed twice with 10 mL Quench buffer and once with complete islet media (RPMI-1640 supplemented with 100 U/mL penicillin/streptomycin, 10% FBS, 1 mM HEPES, 2 mM L-Glutamine, 100 U/mL antibiotic/antimycotic and 10 mM sodium pyruvate). The filter was inverted into a sterile petri dish and cells were washed into the dish with 4.5 mL complete islet media. Islets were left to recover overnight, and treatments began the next day. Mouse islets were treated with ISRIB (100 nM, 24 h), Valinomycin (250 μM, 3 h), or vehicle controls, and were isolated from both male and female mice.

**High-fat diet feeding:** *Ins1*-Cre, β-Clec16a$^{KO}$, or β-Mfn1/2$^{dKO}$ mice were randomized onto a 10% regular-fat diet or 60% high-fat diet (Research Diets Inc; New Brunswick, NJ, USA) at weaning. Access to food was *ad libitum* for 12 weeks. Mice were then subjected to *in vivo* metabolic analyses as required (detailed in following sections), and pancreatic tissue was harvested at the end of the study for immunohistochemistry, *ex vivo* islet studies or sequencing studies.

**Intraperitoneal Glucose Tolerance Tests (IPGTT):** Mice were fasted for 6 h. Fasting blood glucose measurements were taken by tail nick (Bayer Contour glucometer) before an IP injection of 2 mg/kg glucose was administered. Blood glucose measurements were then taken at 15, 30, 60 and 120 mins. Following the test, mice were returned to housing cages with *ad libitum* access to food.

***In vivo* Glucose Stimulated Insulin Release:** Mice were fasted for 6 h. Fasting blood glucose was measured after tail nick with a glucometer (Bayer Contour) and a 20 μL blood sample was collected using capillary tubes (Fisher Scientific) and stored on ice. Mice were injected with 3mg/kg glucose and blood glucose and blood samples were taken after 3 min. Blood samples were ejected from the capillary tubes into 1.5 mL tubes, spun at 16,000 x *g*, 4$^{o}$C for 10 min, and serum was aliquoted to new 1.5 mL tubes. Serum insulin levels were measured by ELISA (Alpco; Salem, NH). Following the test mice were returned to housing cages with *ad libitum* access to food.

**Intraperitoneal Insulin Tolerance Tests (ITT):** Mice were fasted for 6 h. Fasting blood glucose was measured after tail nick with a glucometer (Bayer Contour). Mice were injected with 0.8 U/kg insulin (Humulin R; Eli Lilly; Indianapolis, IN) and blood glucose measured at 15, 30, and 60 min. Following the test mice were returned to housing cages with *ad libitum* access to food.

**Mini-osmotic pump implantation:** While under inhaled isoflurane anesthesia, all mice were implanted subcutaneously with an Alzet micro-osmotic pump (model 2006, Durect, Cupertino, CA, USA) with either vehicle or phlorizin (0.8 mg/kg/d for 6 weeks) as previously described (7). Mice were treated with meloxicam once prior to pump implantation and following implantation (5 mg/kg once daily for up to 3 days), monitored for wound healing or signs of distress, and treated with topical antibiotics (triple antibiotic ointment). Urine glucose concentrations were measured 5 weeks after pump implantation (Germaine Labs).

***In vivo* ISRIB treatment:** Mice were administered 2.5 µg/g ISRIB (ApexBio), dissolved in 40% saline; 50% polyethylene glycol; 10% DMSO, or vehicle alone intraperitoneally daily for 4 weeks.

**Mitophagy assessment in live mouse islets using mtKeima:** mtKeima-Ctrl or mtKeima-β-Clec16a$^{KO}$ islets were isolated and treated for 24 h with vehicle (DMSO) or 100 nM ISRIB. On the day of analysis, islets were treated with ctrl (DMSO) or 250 µM Valinomycin for 3 h. Islets were dispersed to single cells using 500 µL 0.25% trypsin and incubation at 37°C for 3min followed by gentle pipetting and neutralization of the trypsin with FACS buffer (RPMI1640 phenol free + 1% BSA). Single cells were sedimented by centrifugation at room temperature, 2000 rpm, 1 min, and the pelleted cells were washed twice with 1X PBS. The cells were resuspended in 500 µL FACS buffer, transferred to FACS tubes, and incubated with 500 nM Fluozin-3AM (ThermoFisher) for 30 min at 37°C, to label $Zn^{2+}$ enriched insulin granules in β-cells. Cells were centrifuged at 1400rpm for 3min and resuspended in 500 µL FACS buffer. Samples were analyzed on an LSR Fortessa flow cytometer (BD Biosciences). Single cells were gated using forward scatter and side scatter (FSC and SSC, respectively) plots, DAPI staining was used to exclude dead cells, and Fluozin-3 was used to identify β cells. Mitophagy measurements were made using dual laser excitation at 407 nm and 532 nm with an emission laser of 605 nm, as previously described (*8*), and results were analyzed with FlowJo (Tree Star Inc). A total of 5,000 β cells were analyzed per individual experimental replicate.

**Mitophagy assessment in live human islets:** Human islets from non-diabetic donors or those with T2D donors were cultured for a maximum of 48 h after shipment. On the day of the experiment islets were incubated with 100nM MTphagy dye (Dojindo Molecular Technologies; Rockville, MD, USA) for 30 min. Vehicle (DMSO) or 1mM Valinomycin (Sigma) was then added to the islets for 3 h. Islets were then dispersed to single cells using 500 µL 0.25% trypsin and incubation at 37°C for 3min. This was followed by gentle pipetting up and down approx. 10 times and neutralization of the trypsin with human islet FACS buffer (1X KRBH + 1% BSA). Single cells were sedimented by centrifugation at room temperature, 2000 rpm, 1 min, and the pelleted cells were washed twice with 1X PBS. The cells were resuspended in 500 µL human islet FACS buffer, transferred to FACS tubes, and incubated with 500 nM Fluozin-3 and 2.5 nM TMRE for 30 min at 37°C to label insulin granule positive β cells and mitochondrial membrane potential respectively. Cells were centrifuged at 1400rpm, room temperature, 3min and resuspended in 500µL human islet FACS buffer. Samples were analyzed on an LSR Fortessa flow cytometer (BD Biosciences). Single cells were gated using forward scatter and side scatter (FSC and SSC, respectively) plots, DAPI staining was used to exclude dead cells, and Fluozin-3 was used to identify β-cells (fig. S17A-B). Mtphagy measurements in β-cells were made using 488 nm excitation laser with a 710 nm emission filter and analyzed using FlowJo (Tree Star Inc.). A total of 5,000 β cells was quantified from each independent islet preparation.

**mtDNA assessment in human islets:** 25 human islets per donor were handpicked and washed twice with 1X PBS. Islets were pelleted and DNA extraction was performed using the Blood/Tissue DNeasy kit (Qiagen; Germantown, MD, USA) as per manufacturer's instructions. Relative mtDNA content was quantified by qPCR using mtDNA specific primers ND1, and nuclear DNA specific primers for B2M as previously described (*9*). Primer information is provided in table S5.

**Citrate synthase activity in human islets:** Lysates from human islets were used to determine citrate synthase activity with the MitoCheck Citrate Synthase Activity Assay Kit (Cayman Chemical) per manufacturer's instructions.

**mtDNA assessment in flow sorted mouse islets:** Mouse islets were isolated from 4 individual control or β-Tfam$^{KO}$ mice and left to recover overnight. The following day, islets were dispersed to single cells (as per mt-Keima FACS protocol) and sorted on a FACSAria III cell sorted (BD Biosciences). Single cells were gated using forward scatter and side scatter (FSC and SSC, respectively) plots, DAPI staining was used to exclude dead cells, and β cells were sorted from non-β cells based on high 561 nm excitation, 582 nm emission. Positive and negative cell populations were collected in 1X PBS from each mouse, spun to pellet and DNA was extracted using the Blood/Tissue DNeasy kit (Qiagen). Relative mtDNA content was quantified by qPCR using mtDNA specific primers Mt9 and Mt11, and nuclear DNA specific primers Ndufv1 Forward and Reverse (*10*).

**β-cell mass analysis:** Whole mouse pancreas was excised, weighed, and fixed in 4% paraformaldehyde for 16 h at 4°C. Samples were stored in 70% ethanol, 4°C before being embedded in paraffin and sectioned. 3 independent depths of sections, at least 50 μM apart, were dewaxed, and rehydrated and antigen retrieval was carried out using 10 mM sodium citrate (pH 6.0) in a microwave for 10 min. Sections were washed twice with 1X PBS, blocked for 1 h at room temperature with 5% donkey serum in PBT (1X PBS, 0.1% Triton X-100, 1% BSA). Sections were then incubated in the following primary antisera overnight at 4°C in PBT: guinea pig anti-insulin (Abcam; Waltham, MA, USA), rabbit anti-glucagon (Santa Cruz; Dallas, TX, USA). Sections were then washed twice with PBS and incubated for 2 h at room temperature with species specific Cy2 and Cy3 conjugated secondary antibodies. Nuclear labelling was performed using DAPI (Molecular Probes). Sections were scanned using an Olympus IX81 microscope (Olympus; Center Valley, PA, USA) at 10X magnification, with image stitching for quantification (Olympus). β-cell mass quantification (estimated as total insulin positive area/total pancreatic area multiplied by pancreatic weight) was performed on stitched images of complete pancreatic sections from 3 independent regions.

**Immunofluorescence of paraffin embedded sections:** Whole mouse pancreas was excised, weighed, and fixed in 4% paraformaldehyde for 16 h at 4°C. Samples were stored in 70% ethanol at 4°C, before being embedded in paraffin and sectioned. For Aldh1a3/Pdx1 staining, antigen retrieval was carried out using 1 X HistoVT buffer (Nacalai USA Inc; San Diego, CA, USA) in an antigen retriever pressure cooker (Electron Microscopy Sciences; Hatfield, PA, USA). Sections were washed twice with 1X PBS, blocked for 1 h at room temperature with 1 X Blocking One solution (Nacalai USA Inc; San Diego, CA, USA). Sections were then incubated in the following primary antisera overnight at 4°C in 1 X PBS + 1% tween + 20% Blocking One solution: rabbit anti-Aldh1a3 (Novus Biologicals; Littleton, CO, USA). Sections were then washed twice with PBS and incubated for 2 h at room temperature with species specific Cy2 and Cy3 conjugated secondary antibodies. For Ucn3/Pdx1 staining, antigen retrieval was carried out using 10mM sodium citrate (pH 6.0) in a microwave on high for 10min. For Glut2/Pdx1 staining, antigen retrieval was carried out using 1 X R-Buffer A (Electron Microscopy Sciences; Hatfield, PA, USA) in an antigen retriever pressure cooker (Electron Microscopy Sciences; Hatfield, PA, USA). Sections were then washed twice with 1X PBS, blocked for 1 h at room temperature with 5% donkey serum in PBT. Sections were then incubated in the following antisera overnight at 4°C in PBT: rabbit anti-Ucn3 (*11*), or rabbit anti-Glut2 (Millipore Sigma; St Louis, MO, USA) and goat anti-Pdx1 (Abcam; Waltham, MA, USA). Sections were then washed twice with PBS and incubated for 2 h at room temperature with species specific Cy2 and Cy3 conjugated secondary antibodies. Nuclear labelling was performed using DAPI (Molecular Probes). Images were captured under 100X oil immersion magnification on an Olympus IX81 microscope. Quantification of co-positive cells was carried out by a blinded third-party researcher, and samples were re-identified after quantification to reduce bias. Antibody information is provided in table S6.

**Imaging of mitochondrial morphology in mouse β-cells:** Images of pancreas sections stained with SDHA and Insulin (to identify β-cells) were captured with an IX81 microscope (Olympus) using an ORCA Flash4 CMOS digital camera (Hamamatsu). Immunostained pancreatic sections, were captured with Z-stack images and subjected to deconvolution (CellSens; Olympus). Quantitative 3D assessments of mitochondrial morphology and network were performed on ImageJ using Mitochondria Analyzer plugin (*12*). Co-localization analyses were performed on Z-stack images of immunostained dissociated islet cells using the Coloc2 plugin on ImageJ.

**Lineage tracing immunofluorescence analysis:** Whole pancreas was excised, weighed, and fixed in 4% paraformaldehyde for 16 h before being washed with 70 % ethanol and incubated with 50% sucrose solution in PBS overnight. Pancreata were then embedded in OCT (ThermoFisher; Waltham, MA, USA) and frozen and stored at -80°C. 10 μM sections were cut using a Leica CM3050 Cryostat at -20°C, and the slides were stored at -80°C until imaging. Sections were thawed briefly, washed twice with 1X PBS, blocked with 5% donkey serum in PBT, and incubated overnight at 4°C with the following primary antisera: guinea pig anti-insulin (Abcam; Waltham, MA, USA) and rabbit anti-glucagon (Santa Cruz; Dallas, TX, USA). Sections were then washed twice with PBS and incubated for 2 h at room temperature with species specific Cy2 and Cy5 conjugated secondary antibodies, endogenous tdTomato was bright enough to capture without antibody detection. Nuclear labelling was performed using DAPI (Molecular Probes). Images were captured on a Nikon A1 Confocal microscope at 40X oil immersion magnification, z-stacks were captured to ensure quantification of single planes of cells.

**Pseudoislets:** Pseudoislets were prepared using the protocol as described (*13*). Briefly, human islets were handpicked and then dispersed with trypsin (Thermo Scientific). Islet cells were counted and incubated with adenovirus expressing shRNA against human TFAM or a scramble control for 2 hr at MOI of 500. Cells were then seeded at 2000 cells per well in CellCarrier Spheroid Ultra-low attachment microplates (PerkinElmer) in enriched pseudoislet media as described (*13*). Cells were allowed to reaggregate for 7 days before being harvested for cryosections or RNA isolation, cDNA generation, and Sybr-based qRT-PCR measurement as previously described (*1*). Primer information is provided in Table S3.

**EndoC-βH3 cells:** Human EndoC-βH3 cells were grown in DMEM containing 5.6mM glucose, 2% BSA, 50 μM 2-mercaptoethanol, 10mM nicotinamide, 5.5 μg/mL transferrin, 6.7 ng/mL selenite, 100 units/mL penicillin, and 100 units/mL streptomycin (*14*). Cells were transfected using NanoJuice (Novagen) or treated with (-)-Epicatechin (EPI; Sigma) and collected 48hr (for RNA) or 72hr (for protein) later.

**Immunoblot analysis:** Isolated mouse islets, human pseudoislets or frozen liver tissue was homogenized in RIPA buffer containing protease and phosphatase inhibitors. Samples were centrifuged at full speed for 10min, 4 °C to pellet insoluble material and the supernatant was used for immunoblot analysis. Protein quantification was carried out using a Pierce MicroBCA kit (ThermoFisher; Waltham, MA, USA). Up to 25µg of protein lysate from islets and pseudoislets or 150 µg of protein lysate from liver were prepared in Laemmli buffer with DTT and denatured at 70°C for 10min, or 37°C for 30min for OXPHOS analysis. Samples were then run on a 4-15% Tris-glycine protein gel (Bio-Rad; Hercules, CA, USA) at 150V until separated. Samples were then transferred to a nitrocellulose membrane at 90V for 90min, membranes were blocked with 5% skim milk in 1X TBS + 0.05% Tween-20 and incubated overnight at 4 °C with primary antisera. Membranes were then washed 3X with 1X TBS+0.05% Tween-20 and incubated with species-specific HRP conjugated secondary antisera (Vector Labs).

**Perceval imaging:** ATP/ADP ratio was monitored using the PercevalHR biosensor (*15*). Islets were perfused with 16.7 mM glucose to stimulate ATP production in a recording chamber on an Olympus IX-73 inverted epifluorescence microscope (Olympus). PercevalHR was excited at 488 nm using a TILL Polychrome V monochromator (FEI), and a QuantEM:512SC cooled CCD camera (PhotoMetrics) was used to collect emission at 527 nm (*16*). Data were acquired and analyzed using Metafluor (Invitrogen) and plotted using Igor Pro (WaveMetrics Inc.).

**Liver functional analysis:** Serum ALT, LDH and cholesterol were measured as previously published (*17-19*). Liver glycogen was determined using a glycogen assay kit (Cayman Chemical) per manufacturer's instructions. Samples were normalized to tissue weight at time of dissection.

**Bulk RNA-Seq of liver:** Sequencing was performed by the Advanced Genomics Core at University of Michigan Medical School. Total RNA was isolated from frozen liver samples and DNase treated using commercially available kits (Omega Biotek and Ambion, respectively). Libraries were constructed and subsequently subjected to 151 bp paired-end cycles on the NovaSeq-6000 platform (Illumina). FastQC (v0.11.8) was used to ensure the quality of data. Reads were mapped to the reference genome GRCm38 (ENSEMBL), using STAR (v2.6.1b) and assigned count estimates to genes with RSEM (v1.3.1). Alignment options followed ENCODE standards for RNA-seq. FastQC was used in an additional post-alignment step to ensure that only high-quality data were used for expression quantitation and differential expression. Data were pre-filtered to remove genes with 0 counts in all samples. Differential gene expression analysis was performed using DESeq2, using a negative binomial generalized linear model (thresholds: linear fold change >1.5 or <-1.5, Benjamini-Hochberg FDR ($P_{adj}$) <0.05). Plots were generated using variations of DESeq2 plotting functions and other packages with Genialis (Boston, MA, USA).

**Nuclear isolation from mouse islets:** Isolated islets were handpicked, washed twice with PBS, and dissociated to single cells with 500 µL 0.25% trypsin at 37°C for 3min followed by vigorous pipetting and quenching with complete islet media. Single cells were washed twice with PBS and finally resuspended in 1X HBSS + 200 U/mL DNAse and incubated at 37°C for 30min. Cells were washed 4X with PBS to ensure removal of DNAse and resuspended in PBS + 1% BSA. Viability was checked using trypan blue, and only samples above 80% viability were carried forward. Cells were split evenly into 2 portions for differential snRNA and snATAC isolation protocols. For snRNA, cells were pelleted at 500 x *g* for 5min, 4°C, and lysed with 50 µL RNA-LB (10mM Tris-HCl pH7.4, 10mM NaCl, 3mM $MgCl_2$, and 1% NP-40 in sterile nuclease free water) by incubation on ice for 3min. Nuclei were pelleted at 500 x *g* for 5min, 4°C, washed in 500 µL wash buffer (1X

PBS, 1% BSA and 0.2U/µL RNAse inhibitor), spun at 500 x *g* for 5 min, 4°C and finally responded in 100 µL wash buffer. For snATAC, cells were pelleted at 500 x *g* for 5 min, 4°C and lysed in 50 µL ATAC-LB (10 mM Tris-HCl pH7.4, 10 mM NaCl, 3 mM $MgCl_2$, 0.1% NP-40, 0.1% Tween-20 and 0.01% digitonin in sterile nuclease free water) by gentle pipetting 5X and incubation on ice for 3 min. 1mL ATAC-wash buffer (10 mM Tris-HCl pH7.4, 10 mM NaCl, 3 mM $MgCl_2$ and 0.1% Tween-20 in sterile nuclease free water) was added and samples spun at 500 x *g* for 5min, 4°C. Nuclei were resuspended in nuclei buffer (10X Genomics). After all nuclei isolation protocols, nuclei were counted, and the Advanced Genomics Core at the University of Michigan processed the samples for sequencing.

**Single-nucleus RNA-sequencing (snRNA-seq) and quantification:** Approximately 14,000 single nuclei were loaded onto a 10X Genomics Chromium machine and resulting captured cells were sequenced via 10X Genomics Single Cell 3P v3 chemistry (10x Genomics, USA), yielding approximately 7k nuclei per lane. All libraries were analyzed using the Cell Ranger analysis pipeline version 3.0. Briefly, reads were aligned to mouse reference sequence mm10 and unique molecular identifiers (UMIs) were quantified using the Cell Ranger "count" software with default parameters. Count data from each single nucleus were normalized to the same sequencing depth and then aggregated using the "cellranger aggr" tool. Cell barcodes with fewer than 1000 detected genes and a total mitochondrial content > 20% (signifying potential apoptotic cells) were discarded from analyses.

**Single-nucleus ATAC-sequencing (snATAC-seq) and quantification:** Approximately 14,000 single nuclei were loaded and sequenced via 10X Genomics Single Cell ATAC v1 chemistry (10x Genomics, USA), yielding approximately 7K cells per lane. All libraries were sequenced individually on Illumina NovaSeq S4 flowcells and were analyzed using the Cell Ranger ATAC analysis pipeline version 1.2.0. Reads were similarly aligned to mouse reference sequence mm10 and fragments, open chromatin peaks were quantified using the Cell Ranger ATAC “count” software with default parameters. Fragment data from each single cell library were normalized to the same sequencing depth and then aggregated using the “cellranger aggr” tool. Resultant genomic alignment (BAM) files were filtered to contain only nucleosome free reads (<147 bp) using SAMtools version 1.8 (*20*).

**snRNA-seq doublet cell detection and removal:** To identify and remove potential multiplet droplets from snRNA-seq data, we used DoubletFinder (*21*) and Scrublet (*22*) to identify multiplet nuclei in an unbiased/non-cell type specific manner. For both methods, we assumed an expected doublet rate of 10% of total cells. Afterwards, we removed all multiplet nuclei identified by at least one of these two methods (union set of doublets).

**snRNA-seq cell clustering:** Nuclei were clustered using top 2000 most variable genes, which are determined by fitting a line to the relationship of the variance and mean using a local polynomial regression (loess). Gene expression values were standardized using the observed normalized mean and expected variance (given by the fitted line). Feature variance was then calculated on the standardized values after clipping to a maximum of the square root of the number of cells. After scaling and centering the feature values, principal component analysis (PCA) was performed on the top 2000 variable features. Following this, uniform manifold approximation and projection (UMAP) was applied on the top 20 principal components to reduce the data to two dimensions. Next, a shared nearest neighbor (SNN) graph was created by calculating the neighborhood (Jaccard index) overlap between each cell and its 20 (default) nearest neighbors. To identify cell clusters, a SNN modularity optimization-based clustering algorithm was used (*23*).

**Annotation of cell types in snRNA-seq:** Islet cell clusters were annotated based on expression of known cell-type markers —*Ins1/2* (beta), *Gcg* (alpha), *Sst* (delta). The remaining cells were annotated using pathway databases and annotations from the *clusterProfiler* (version 3.14.0 (*24*)) R package.

**Differential gene expression analysis:** Differential expression analyses was performed using the FindMarkers function (with default parameters) and the wilcox test option in the R package Seurat (*23*) to identify genes that are enriched in each cell type. In each comparison, protein coding genes and long intergenic non-coding RNAs (lincRNAs) with one or more UMIs in at least 10% of either cell type population being compared was used. Differentially expressed genes with FDR < 5% were regarded as significant results.

**Pathway analysis of differentially expressed genes:** Differentially expressed genes for each cell-type were functionally annotated using the R package *clusterProfiler* (version 3.14.0 (*24*)). The Kyoto Encyclopedia of Genes and Genomes (KEGG), gene ontology (GO), and WikiPathways databases were used to determine associations with particular biological processes, diseases, and molecular functions. The top pathways with an FDR-adjusted p-value <5% were summarized in the results.

**Integration of snRNA-seq and snATAC-seq data:** Integrative clustering and analysis of single nuclei transcriptomes and single nucleus epigenomes was performed using the R package Seurat (*23*). First, gene activity scores were derived from the resultant snATAC-seq peak count-matrix using the CreateGeneActivityMatrix function with default parameters. Next, single nuclei with < 5,000 total read counts were discarded from analyses. The resultant single nuclei and gene activity scores were log normalized and scaled. Using the processed snRNA-seq data (also analyzed with Seurat), we identified anchors between the snATAC-seq gene activity score matrix and snRNA-seq gene expression matrix following the methodology described in (*23*). After identifying anchors between the datasets, cell-type labels from the snRNA-seq dataset were transferred to the snATAC-seq dataset and a prediction and confidence score was assigned for each cell.

**Aggregation of snATAC-seq profiles into pseudo-bulk profiles:** For each cell-type, snATAC-seq profiles were aggregated into single bulk ATAC-seq profiles using using SAMtools version 1.8 (*20*) and the "merge" command. Merged BAM files were sorted and indexed using SAMtools and provided as input to identify open chromatin peaks. Peaks were called using MACS version 2.1.0.20151222 and the following parameters "-g 'mm' --nomodel --call-summits --qval 0.01 --keep-dup all -B -f BAMPE". Cell-type specific open chromatin peaks were identified using BEDtools version 2.27.0(*25*) and the "bedtools intersect -v" command.

**Transcription factor motif enrichment analysis:** The *findMotifsGenome.pl* (HOMER version 4.6 (*26*)) script with parameters "hg19 -size 200" was used to determine TF motifs enriched in ATAC-seq peaks for KO beta cells relative to control beta cells (and vice versa). As an example, to identify TF motifs enriched in KO beta cell-specific peaks, we provided these peaks as the foreground, and the control beta cell-specific peaks as the background.

**Statistics**

In all figures, data are presented as means ± SEM, and error bars denote SEM, unless otherwise noted in the legends. Outlier tests (ROUT method (*27*)) were routinely performed in GraphPad Prism. Statistical comparisons were performed using unpaired two-tailed Student's *t*-tests, one-way or two-way ANOVA, followed by Tukey's or Sidak's post-hoc test for multiple comparisons, as appropriate (GraphPad Prism). A $P$ value < 0.05 was considered significant.

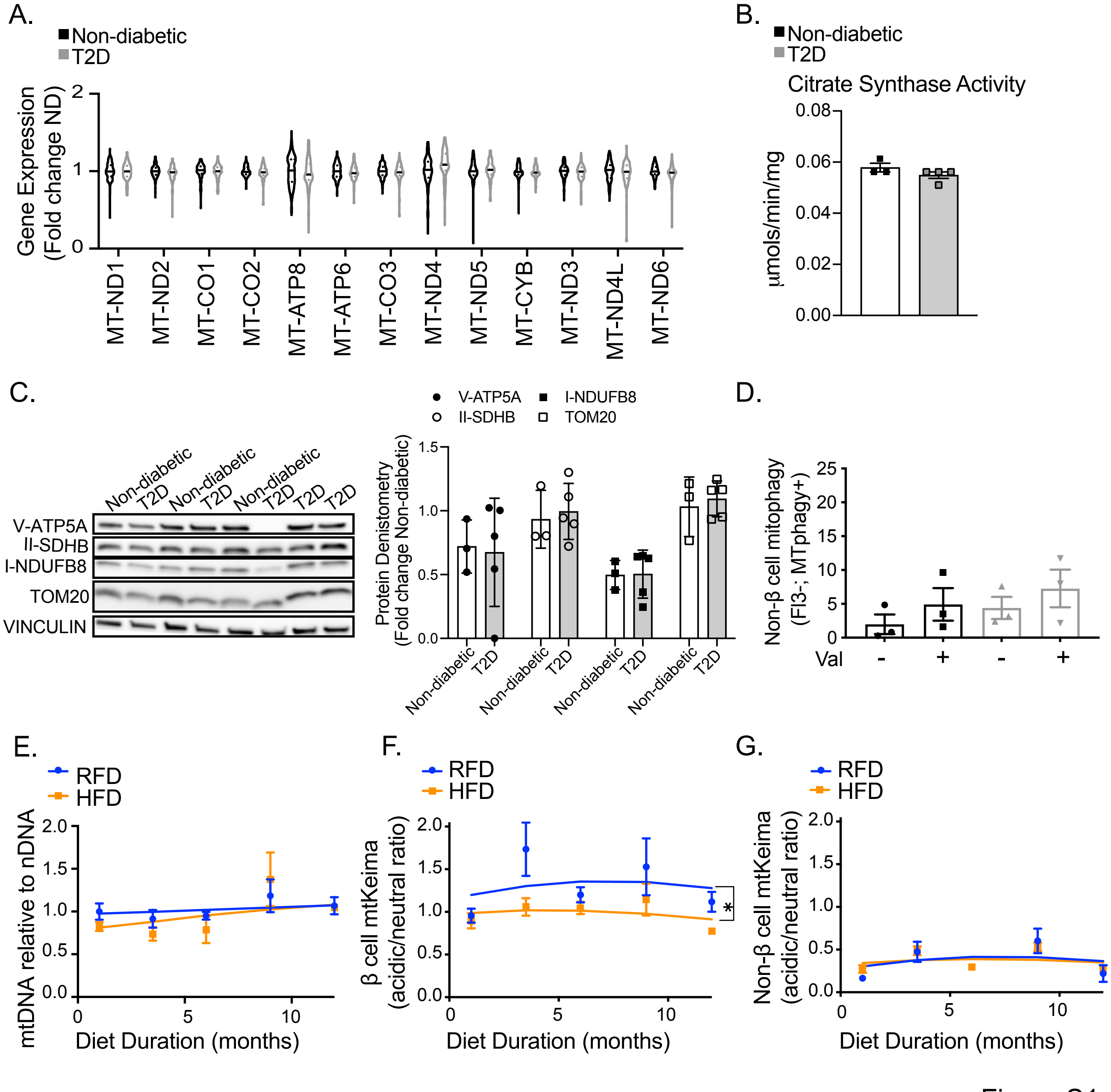
A.
Non-diabetic
T2D
Gene Expression (Fold change ND)
MT-ND1
MT-ND2
MT-CO1
MT-CO2
MT-ATP8
MT-ATP6
MT-CO3
MT-ND4
MT-ND5
MT-CYB
MT-ND3
MT-ND4L
MT-ND6
B.
Non-diabetic
T2D
Citrate Synthase Activity
μmols/min/mg
C.
V-ATP5A
II-SDHB
I-NDUFB8
TOM20
VINCULIN
Non-diabetic
T2D
Protein Denistometry (Fold change Non-diabetic)
D.
Non-β cell mitophagy (Fl3-; MTphagy+)
Val
E.
RFD
HFD
mtDNA relative to nDNA
Diet Duration (months)
F.
β cell mtKeima (acidic/neutral ratio)
G.
Non-β cell mtKeima (acidic/neutral ratio)

Figure S1

**Figure S1. Mitochondrial quality control defects appear to be intrinsic to β-cells in T2D.** (A) Violin plot showing levels of mitochondrially encoded transcripts from single α-cell analysis of cells from human non-diabetic (ND) donors or those with T2D. *n* = 3-5/group. (B) Citrate synthase activity of non-diabetic or T2D donor islets. (C) Western blot and quantitation from non-diabetic and T2D donors for various OXPHOS proteins and TOM20. Vinculin serves as a loading control. *n* = 3-5/group. (D) Flow cytometry quantification of percentage of MTPhagy+ non-β-cells from non-diabetic donors or those with T2D. *n* = 3-4/group. (E) Quantification of mtDNA:nuclear DNA ratio from islets of mt-Keima mice at various time points of HFD feeding. *n* = 4-6/group. (F) Flow cytometry quantification of ratio of acidic:neutral localized mitochondria in β-cells from RFD- or HFD-fed mt-Keima mice at various time points. *$P < 0.05$ effect of diet by two-way ANOVA. *n* = 4-6/group. (G) Flow cytometry quantification of ratio of acidic:neutral localized mitochondria in non-β-cells from RFD- or HFD-fed mt-Keima mice at various time points. n = 4-6/group.

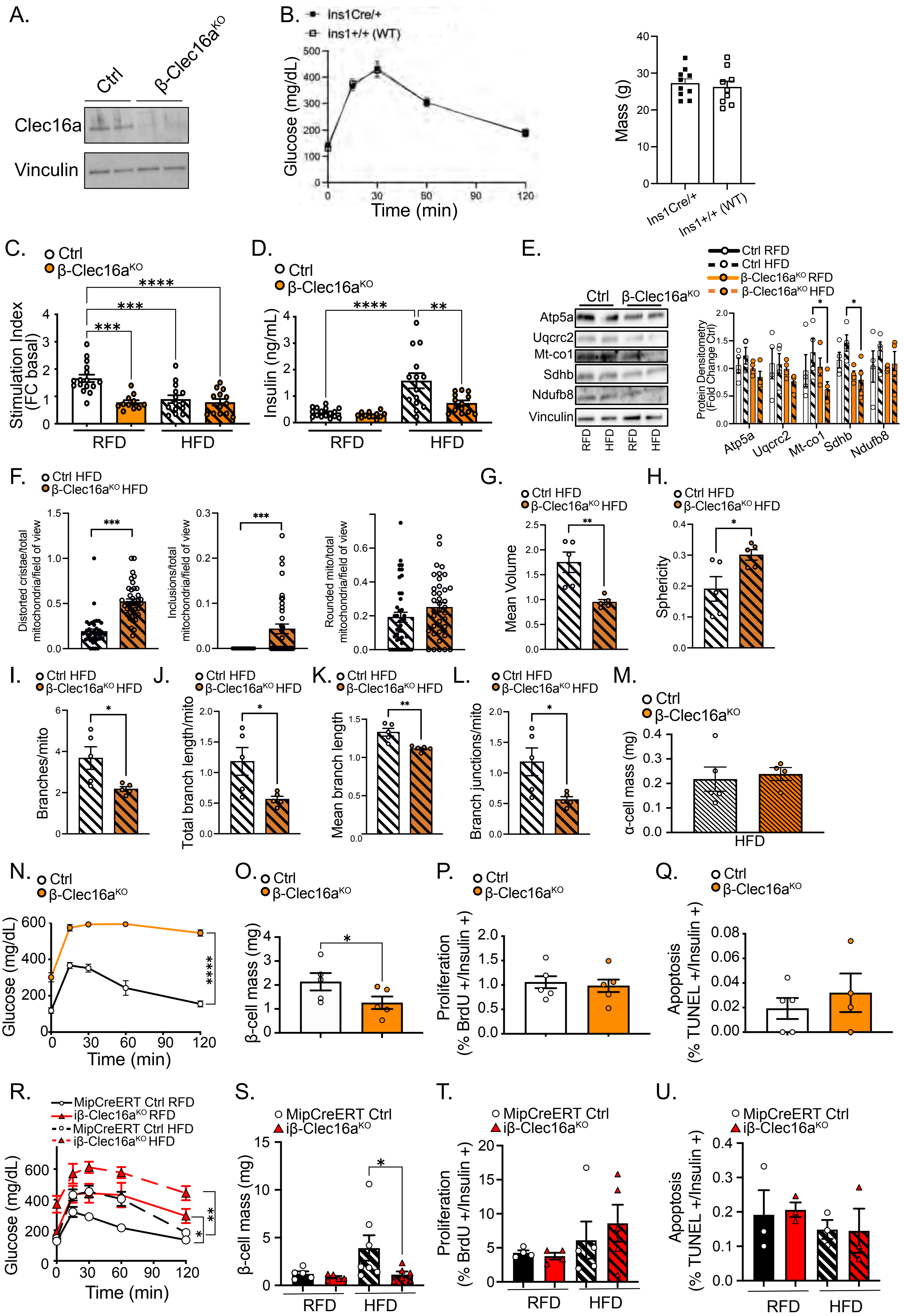
A.
Ctrl
β-Clec16a^KO
Clec16a
Vinculin
B.
Ins1Cre/+
Ins1+/+ (WT)
Glucose (mg/dL)
Time (min)
Mass (g)
C.
Ctrl
β-Clec16a^KO
Stimulation Index (FC basal)
RFD
HFD
D.
Insulin (ng/mL)
E.
Atp5a
Uqcrc2
Mt-co1
Sdhb
Ndufb8
Vinculin
Ctrl RFD
Ctrl HFD
β-Clec16a^KO RFD
β-Clec16a^KO HFD
Protein Densitometry (Fold Change Ctrl)
F.
Ctrl HFD
β-Clec16a^KO HFD
Distorted cristae/total mitochondria/field of view
Inclusions/total mitochondria/field of view
Rounded mito/total mitochondria/field of view
G.
Mean Volume
H.
Sphericity
I.
Branches/mito
J.
Total branch length/mito
K.
Mean branch length
L.
Branch junctions/mito
M.
α-cell mass (mg)
N.
Glucose (mg/dL)
Time (min)
O.
β-cell mass (mg)
P.
Proliferation (% BrdU +/Insulin +)
Q.
Apoptosis (% TUNEL +/Insulin +)
R.
MipCreERT Ctrl RFD
iβ-Clec16a^KO RFD
MipCreERT Ctrl HFD
iβ-Clec16a^KO HFD
Glucose (mg/dL)
Time (min)
S.
MipCreERT Ctrl
iβ-Clec16a^KO
β-cell mass (mg)
T.
Proliferation (% BrdU +/Insulin +)
U.
Apoptosis (% TUNEL +/Insulin +)

Figure S2

**Figure S2. Reductions in β-cell mass in adult mice following mitophagy deficiency are exacerbated by aging and obesity and not due to developmental defects.** (A) Representative WB demonstrating expression of Clec16a in islets isolated from 16-week-old Ctrl or β-Clec16a$^{KO}$ mice. Vinculin serves as a loading control. $n$ = 4/group. (B) Change in blood glucose levels during an ITT. $n$ = 9-15 mice per group. (C) Blood glucose concentration measured during an IPGTT of Ins1Cre/+ or Ins1+/+ following 15-weeks high fat diet (HFD). $n$ = 9-10 mice per group. (D) Weights of Ins1Cre/+ or Ins1+/+ mice following 15-weeks high fat diet (HFD). $n$ = 9-10 mice per group. (E) *In vivo* insulin secretion presented as stimulation index. $n$ = 12-16 mice per group. ***$P$ < 0.001, ****$P$ < 0.0001 by one-way ANOVA, Tukey's multiple comparison post-test. (F) Serum insulin levels following a 6hr fast. $n$ = 12-16 mice per group. **$P$ < 0.01, ****$P$ < 0.0001 by one-way ANOVA, Tukey's multiple comparison post-test. (G) Representative western blot and quantitation from Ctrl or β-Clec16a$^{KO}$ islets fed RFD or HFD for OXPHOS proteins. Vinculin serves as a loading control. $n$ = 4/group. *$P$ < 0.05 by two-way ANOVA, Tukey's multiple comparison post-test. (H) Quantification of mitochondrial morphology in Ctrl and β-Clec16a$^{KO}$ from TEM images. $n$ = 2-3/group. At least 40 fields of view were quantitated per sample. (I-N) β-cell mitochondrial morphology and network analysis of deconvolution immunofluorescence Z-stack images stained for SDHA (and insulin) from pancreatic sections of Ctrl and β-Clec16a$^{KO}$ RFD or HFD mice. (O) Pancreatic α-cell mass from 12-week HFD-fed Ctrl or β-Clec16a$^{KO}$ mice. $n$ = 4-5 animals per group. (P-S) Results from 1-year old Ctrl or β- Clec16a$^{KO}$ mice fed RFD. (P) Blood glucose concentrations during an IPGTT. $n$ = 5-6 mice per group. ****$P$ < 0.0001 by two-way ANOVA effect of genotype. (Q) Pancreatic β-cell mass. $n$ = 5 mice per group. *$P$ < 0.05 by Student's unpaired $t$-test. (R) β-cell proliferation as measured by % BrdU incorporated Insulin$^{+}$ cells. (S) β-cell apoptosis as measured by TUNEL$^{+}$/Insulin$^{+}$ cells. (T-W) Results from tamoxifen inducible Clec16a$^{KO}$ (iβ-Clec16a$^{KO}$) or Ctrl (MipCreERT) mice, following tamoxifen injection in 6-week-old animals. (T) Blood glucose concentrations during an IPGTT in MipCreERT or iβ-Clec16a$^{KO}$ mice on RFD or HFD for 18weeks. $n$ = 4-7 mice/group. *$P$ < 0.05, **$P$ < 0.01 by two-way ANOVA effect of genotype on respective diets. (U) Pancreatic β-cell mass $n$ = 5-7 mice/group; *$P < 0.05$ by one-tailed Student's unpaired $t$-test. (V) β-cell proliferation as measured by % BrdU incorporated $Insulin^+$ cells. (W) β-cell apoptosis as measured by $TUNEL^+/Insulin^+$ cells.

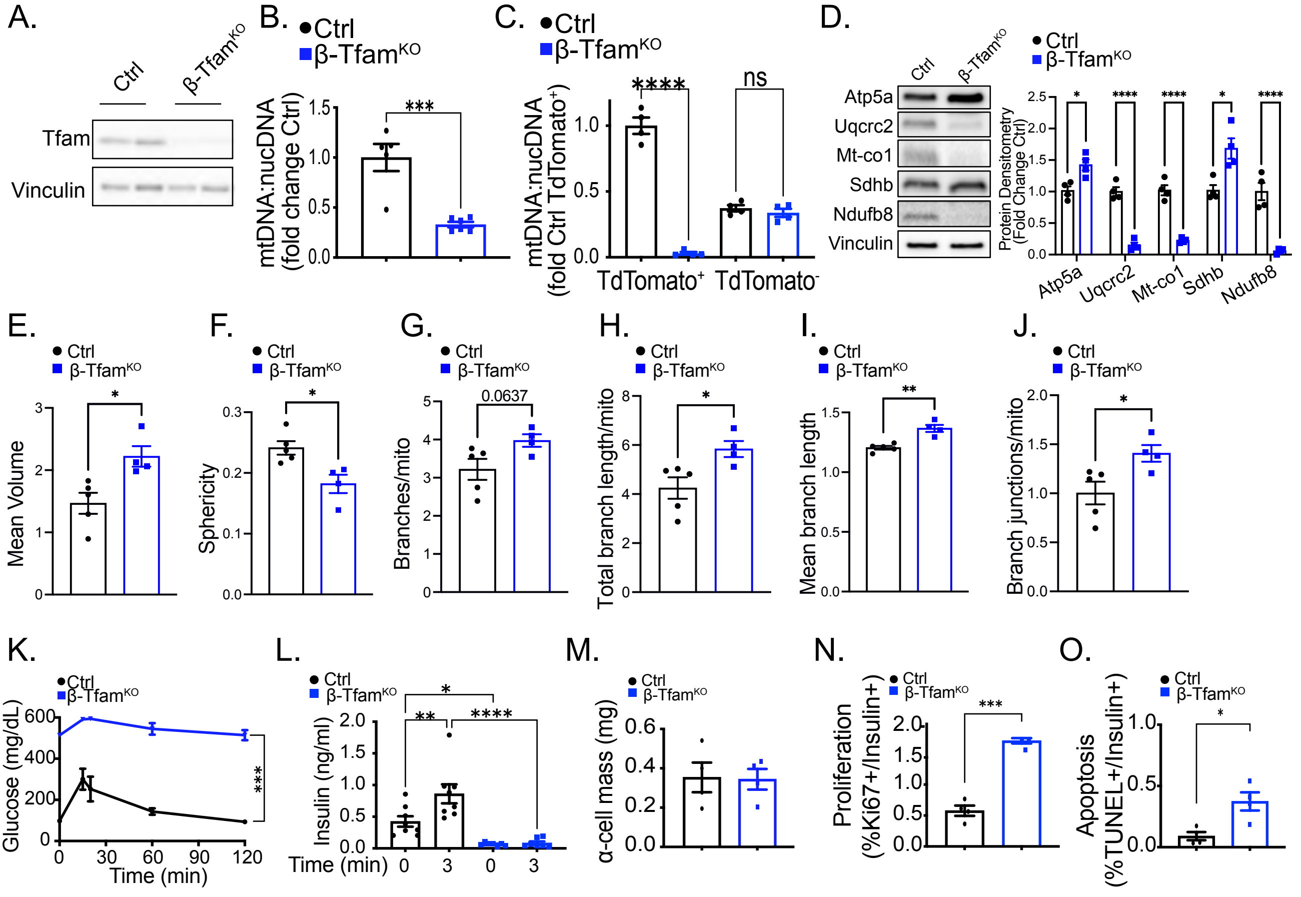

A.
Ctrl
β-Tfam^KO
Tfam
Vinculin
B.
Ctrl
β-Tfam^KO
mtDNA:nucDNA (fold change Ctrl)
***
C.
Ctrl
β-Tfam^KO
mtDNA:nucDNA (fold Ctrl TdTomato+)
****
ns
TdTomato+
TdTomato-
D.
Ctrl
β-Tfam^KO
Atp5a
Uqcrc2
Mt-co1
Sdhb
Ndufb8
Vinculin
Protein Densitometry (Fold Change Ctrl)
E.
Mean Volume
F.
Sphericity
G.
Branches/mito
0.0637
H.
Total branch length/mito
I.
Mean branch length
J.
Branch junctions/mito
K.
Glucose (mg/dL)
Time (min)
L.
Insulin (ng/ml)
Time (min)
M.
α-cell mass (mg)
N.
Proliferation (%Ki67+/Insulin+)
O.
Apoptosis (%TUNEL+/Insulin+)


Figure S3

**Figure S3. Characterization of mice bearing β-cell specific Tfam deficiency reveals mtDNA depletion, alterations in OXPHOS subunit expression, increases in mitochondrial density, and worsening glucose intolerance with age.** (A) Representative WB demonstrating expression of Tfam in isolated islets from 10-week-old Ctrl or β-$Tfam^{KO}$ mice. Vinculin serves as a loading control. *n* = 4/group. (B) Quantification of mtDNA versus nuclear DNA ratio from islets of Ctrl or β-$Tfam^{KO}$ mice. *n* = 5-6 mice per group. ****P* < 0.001 by Student's unpaired *t*-test. (C) Quantification of mtDNA versus nuclear DNA from live FACS sorted islet cells of Ctrl or β-$Tfam^{KO}$ mice. *n* = 4 mice per group. *****P* < 0.0001 by one-way ANOVA, Tukey's multiple comparison post-test. (D) Representative western blot and quantitation from Ctrl or β-$Tfam^{KO}$ islets for OXPHOS proteins. Vinculin serves as a loading control. *n* = 3-4/group. **P* < 0.05, *****P* < 0.0001 by Student's unpaired *t*-test. (E-J) β-cell mitochondrial morphology and network analysis of deconvolution immunofluorescence Z-stack images stained for SDHA (and insulin) from pancreatic sections of Ctrl and β-$Tfam^{KO}$ mice by MitoAnalyzer. *n* = 4-5/group. (K) Blood glucose concentrations during an IPGTT of Ctrl or β-$Tfam^{KO}$ mice at 14-weeks of age. *n* = 4-6 animals per group. ****P* < 0.001 by two-way ANOVA effect of genotype. (L) Serum insulin measured during *in vivo* glucose-stimulated insulin release in 10-week-old Ctrl or β-$Tfam^{KO}$ mice. *n* = 7-8 animals per group. **P* < 0.05, ***P* < 0.01, *****P* < 0.0001 by one-way ANOVA, Tukey's multiple comparison post-test. (M) Pancreatic α-cell mass from 12-week-old Ctrl or β-$Tfam^{KO}$ mice. *n* = 4 animals per group. (N) β-cell replication measured as the % of Ki67+/Insulin+ cells in 12-week-old mice. *n* = 4 mice per group. ****P* < 0.001 by Student's unpaired *t*-test. (O) β-cell apoptosis measured as the % of TUNEL+/Insulin+ cells. *n* = 4-11 mice per group. **P* < 0.05 by Student's unpaired *t*-test.

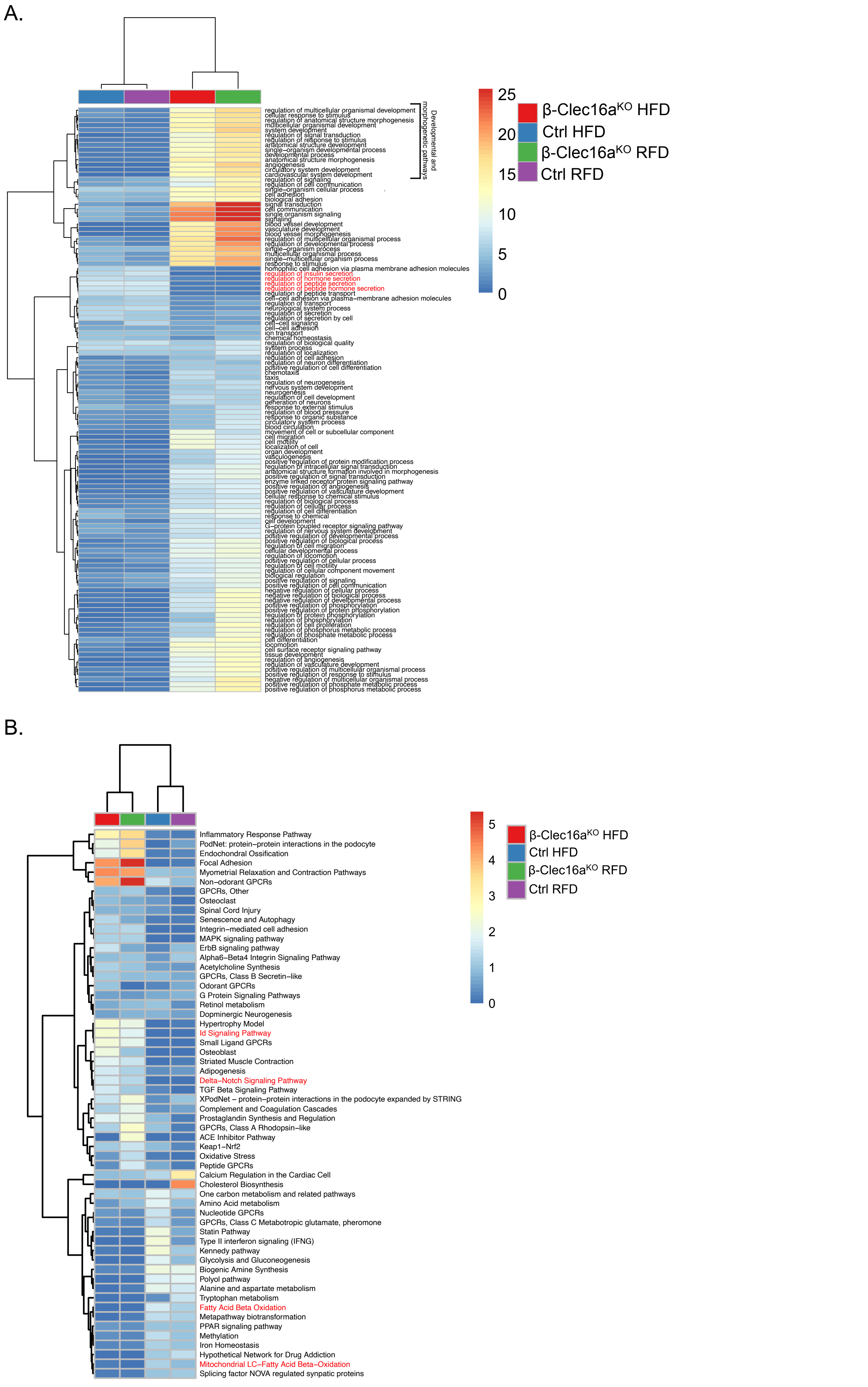
A.
regulation of multicellular organismal development
cellular response to stimulus
regulation of anatomical structure morphogenesis
multicellular organismal development
system development
regulation of signal transduction
regulation of response to stimulus
anatomical structure development
single-organism developmental process
developmental process
anatomical structure morphogenesis
angiogenesis
circulatory system development
cardiovascular system development
regulation of signaling
regulation of cell communication
single-organism cellular process
cell adhesion
biological adhesion
signal transduction
cell communication
single organism signaling
signaling
blood vessel development
vasculature development
blood vessel morphogenesis
regulation of multicellular organismal process
regulation of developmental process
single-organism process
multicellular organismal process
single-multicellular organism process
response to stimulus
homophilic cell adhesion via plasma membrane adhesion molecules
regulation of insulin secretion
regulation of hormone secretion
regulation of peptide secretion
regulation of peptide hormone secretion
regulation of peptide transport
cell-cell adhesion via plasma-membrane adhesion molecules
regulation of transport
neurological system process
regulation of secretion
regulation of secretion by cell
cell-cell signaling
cell-cell adhesion
ion transport
chemical homeostasis
regulation of biological quality
system process
regulation of localization
regulation of cell adhesion
regulation of neuron differentiation
positive regulation of cell differentiation
chemotaxis
taxis
regulation of neurogenesis
nervous system development
neurogenesis
regulation of cell development
generation of neurons
response to external stimulus
regulation of blood pressure
response to organic substance
circulatory system process
blood circulation
movement of cell or subcellular component
cell migration
cell motility
localization of cell
organ development
vasculogenesis
positive regulation of protein modification process
regulation of intracellular signal transduction
anatomical structure formation involved in morphogenesis
positive regulation of signal transduction
enzyme linked receptor protein signaling pathway
positive regulation of angiogenesis
positive regulation of vasculature development
cellular response to chemical stimulus
regulation of biological process
regulation of cellular process
regulation of cell differentiation
response to chemical
cell development
G-protein coupled receptor signaling pathway
regulation of nervous system development
positive regulation of developmental process
positive regulation of biological process
regulation of cell migration
cellular developmental process
regulation of locomotion
positive regulation of cellular process
regulation of cell motility
regulation of cellular component movement
biological regulation
positive regulation of signaling
positive regulation of cell communication
negative regulation of cellular process
negative regulation of biological process
negative regulation of developmental process
positive regulation of phosphorylation
positive regulation of protein phosphorylation
regulation of protein phosphorylation
regulation of phosphorylation
regulation of cell proliferation
regulation of phosphorus metabolic process
regulation of phosphate metabolic process
cell differentiation
locomotion
cell surface receptor signaling pathway
tissue development
regulation of angiogenesis
regulation of vasculature development
positive regulation of multicellular organismal process
positive regulation of response to stimulus
negative regulation of multicellular organismal process
positive regulation of phosphate metabolic process
positive regulation of phosphorus metabolic process
Developmental and morphogenetic pathways
25
20
15
10
5
0
β-Clec16a^KO HFD
Ctrl HFD
β-Clec16a^KO RFD
Ctrl RFD
B.
Inflammatory Response Pathway
PodNet: protein-protein interactions in the podocyte
Endochondral Ossification
Focal Adhesion
Myometrial Relaxation and Contraction Pathways
Non-odorant GPCRs
GPCRs, Other
Osteoclast
Spinal Cord Injury
Senescence and Autophagy
Integrin-mediated cell adhesion
MAPK signaling pathway
ErbB signaling pathway
Alpha6-Beta4 Integrin Signaling Pathway
Acetylcholine Synthesis
GPCRs, Class B Secretin-like
Odorant GPCRs
G Protein Signaling Pathways
Retinol metabolism
Dopminergic Neurogenesis
Hypertrophy Model
Id Signaling Pathway
Small Ligand GPCRs
Osteoblast
Striated Muscle Contraction
Adipogenesis
Delta-Notch Signaling Pathway
TGF Beta Signaling Pathway
XPodNet - protein-protein interactions in the podocyte expanded by STRING
Complement and Coagulation Cascades
Prostaglandin Synthesis and Regulation
GPCRs, Class A Rhodopsin-like
ACE Inhibitor Pathway
Keap1-Nrf2
Oxidative Stress
Peptide GPCRs
Calcium Regulation in the Cardiac Cell
Cholesterol Biosynthesis
One carbon metabolism and related pathways
Amino Acid metabolism
Nucleotide GPCRs
GPCRs, Class C Metabotropic glutamate, pheromone
Statin Pathway
Type II interferon signaling (IFNG)
Kennedy pathway
Glycolysis and Gluconeogenesis
Biogenic Amine Synthesis
Polyol pathway
Alanine and aspartate metabolism
Tryptophan metabolism
Fatty Acid Beta Oxidation
Metapathway biotransformation
PPAR signaling pathway
Methylation
Iron Homeostasis
Hypothetical Network for Drug Addiction
Mitochondrial LC-Fatty Acid Beta-Oxidation
Splicing factor NOVA regulated synpatic proteins
5
4
3
2
1
0
β-Clec16a^KO HFD
Ctrl HFD
β-Clec16a^KO RFD
Ctrl RFD

Figure S4

**Figure S4. RNA sequencing highlights defects in terminal β-cell identity and activation of a precursor state following mitophagy deficiency.** (A-B) Heatmap representations from Ctrl (blue or purple) or β-Clec16a$^{KO}$ (red or green) mice fed HFD or RFD as denoted, (A) WikiPathways biological terms analysis from islets with downregulation of hormone processing, insulin secretion and hormone secretion (red highlighted terms). (B) Heatmap representation of WikiPathways GO-term analysis with upregulation of abnormal mature cell differentiation signaling (Id, Delta-Notch pathways) and downregulation of mitochondrial function (reduced β-oxidation, and mitochondrial LC-fatty acid oxidation) (red highlighted terms).

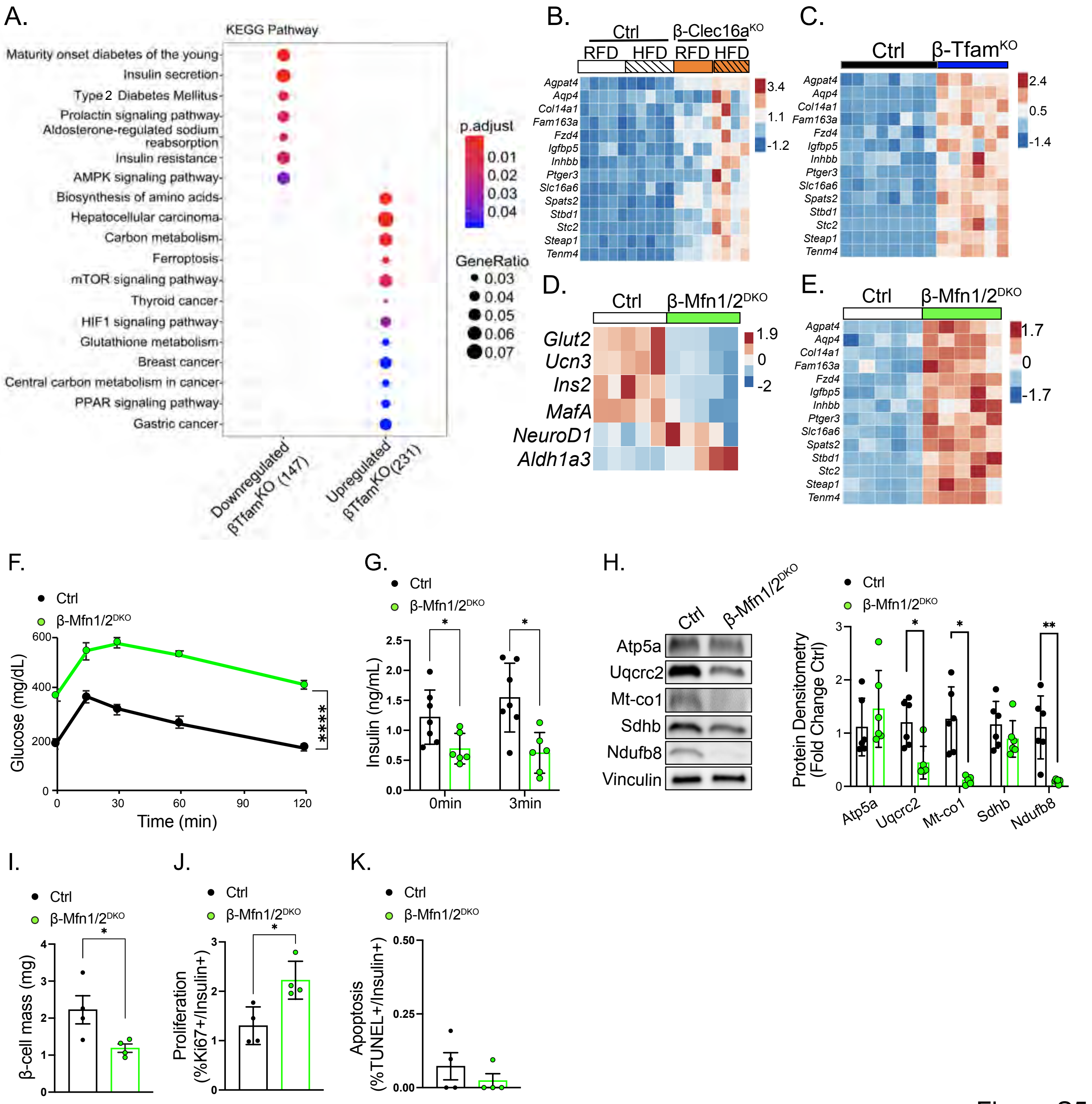
A.
KEGG Pathway
Maturity onset diabetes of the young
Insulin secretion
Type 2 Diabetes Mellitus
Prolactin signaling pathway
Aldosterone-regulated sodium reabsorption
Insulin resistance
AMPK signaling pathway
Biosynthesis of amino acids
Hepatocellular carcinoma
Carbon metabolism
Ferroptosis
mTOR signaling pathway
Thyroid cancer
HIF1 signaling pathway
Glutathione metabolism
Breast cancer
Central carbon metabolism in cancer
PPAR signaling pathway
Gastric cancer
Downregulated βTfamKO (147)
Upregulated βTfamKO(231)
p.adjust
0.01
0.02
0.03
0.04
GeneRatio
0.03
0.04
0.05
0.06
0.07
B.
Ctrl
β-Clec16aKO
RFD HFD RFD HFD
Agpat4 Aqp4 Col14a1 Fam163a Fzd4 Igfbp5 Inhbb Ptger3 Slc16a6 Spats2 Stbd1 Stc2 Steap1 Tenm4
3.4
1.1
-1.2
C.
Ctrl
β-TfamKO
2.4
0.5
-1.4
D.
Ctrl
β-Mfn1/2DKO
Glut2 Ucn3 Ins2 MafA NeuroD1 Aldh1a3
1.9
0
-2
E.
Ctrl
β-Mfn1/2DKO
1.7
0
-1.7
F.
Ctrl
β-Mfn1/2DKO
Glucose (mg/dL)
Time (min)
****
G.
Insulin (ng/mL)
0min
3min
H.
Ctrl
β-Mfn1/2DKO
Atp5a
Uqcrc2
Mt-co1
Sdhb
Ndufb8
Vinculin
Protein Densitometry (Fold Change Ctrl)
I.
β-cell mass (mg)
J.
Proliferation (%Ki67+/Insulin+)
K.
Apoptosis (%TUNEL+/Insulin+)

Figure S5

**Figure S5. RNA sequencing highlights that loss of mtDNA content and mitochondrial fusion lead to impaired β-cell maturity.** (A) KEGG pathway analysis from bulk RNAseq depicting downregulated or upregulated pathways overrepresented in islets from β-$Tfam^{KO}$ mice. (B-C) Heatmaps showing upregulation of immature genes in islets from Ctrl and β-$Clec16a^{KO}$ mice fed HFD or RFD (B) or islets from Ctrl and β-$Tfam^{KO}$ mice (C). (D-E) Heatmaps generated from bulk RNAseq of Ctrl or β-$Mfn1/2^{DKO}$ islets displaying levels of selected mature and immature genes. (F) Blood glucose concentrations during an IPGTT of Ctrl or β-$Mfn1/2^{DKO}$ mice on HFD at 10-weeks of age. $n$ = 4 animals per group. ****$P < 0.0001$ by Student's unpaired *t*-test. (G) Serum insulin measured during *in vivo* glucose-stimulated insulin release in 10-week-old Ctrl or β-$Mfn1/2^{DKO}$ mice on HFD. $n$ = 7-8 animals per group. *$P < 0.05$ by Student's unpaired *t*-test. (H) Representative western blot and quantitation from islets of Ctrl or β-$Mfn1/2^{DKO}$ on HFD for OXPHOS proteins. Vinculin serves as a loading control. $n$ = 6/group. *$P < 0.05$; **$P < 0.01$ Student's unpaired *t*-test. (I) Pancreatic β-cell mass from 10-week-old Ctrl or β-$Mfn1/2^{DKO}$ mice on HFD. n = 4 mice per group. *$P < 0.05$ by Student's unpaired *t*-test. (J) β-cell replication measured as the % of Ki67+/Insulin+ cells in 10-week-old mice. $n$ = 4 mice per group. *$P < 0.05$ by Student's unpaired *t*-test. (K) β-cell apoptosis measured as the % of TUNEL+/Insulin+ cells. $n$ = 4 mice per group.

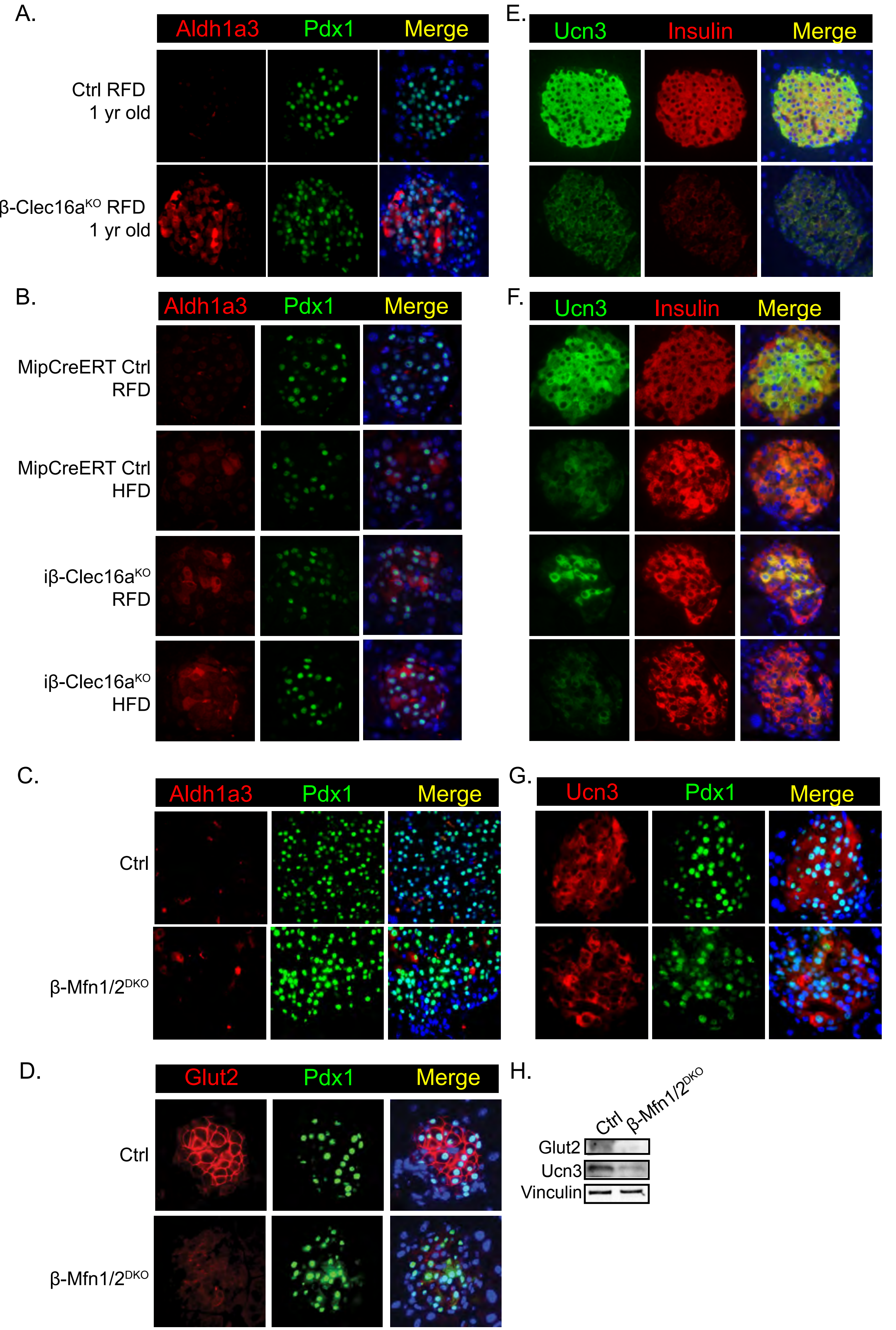
A.
Aldh1a3
Pdx1
Merge
Ctrl RFD
1 yr old
β-Clec16aKO RFD
1 yr old
B.
Aldh1a3
Pdx1
Merge
MipCreERT Ctrl
RFD
MipCreERT Ctrl
HFD
iβ-Clec16aKO
RFD
iβ-Clec16aKO
HFD
C.
Aldh1a3
Pdx1
Merge
Ctrl
β-Mfn1/2DKO
D.
Glut2
Pdx1
Merge
Ctrl
β-Mfn1/2DKO
E.
Ucn3
Insulin
Merge
F.
Ucn3
Insulin
Merge
G.
Ucn3
Pdx1
Merge
H.
Ctrl
β-Mfn1/2DKO
Glut2
Ucn3
Vinculin

Figure S6

**Figure S6. Reductions in β-cell maturity are exacerbated by aging and obesity following loss of mitophagy and are also observed following impaired mitochondrial fusion.** (A-G) Representative immunofluorescence images of dedifferentiation and maturity markers in Ctrl or β-Clec16a$^{KO}$ islets at 1 yr of age; Ctrl or iβ-Clec16a$^{KO}$ islets at 24 weeks of age following 18 weeks of deletion and either RFD or HFD feeding; and Ctrl or β-Mfn1/2$^{DKO}$ islets. *n* = 4-6/group for all studies. (A-C) Immunostaining with antibodies against Aldh1a3 (red), Pdx1 (green) and DAPI (blue). (D) Immunostaining with antibodies against Glut2 (red), Pdx1 (green) and DAPI (blue) and (E-G) Ucn3 (green), Insulin (red) and DAPI (blue). (H) Protein expression of Glut2 and Ucn3 by WB in of islets from Ctrl or β-Mfn1/2$^{DKO}$ mice on HFD. *n* = 3-6 per group.

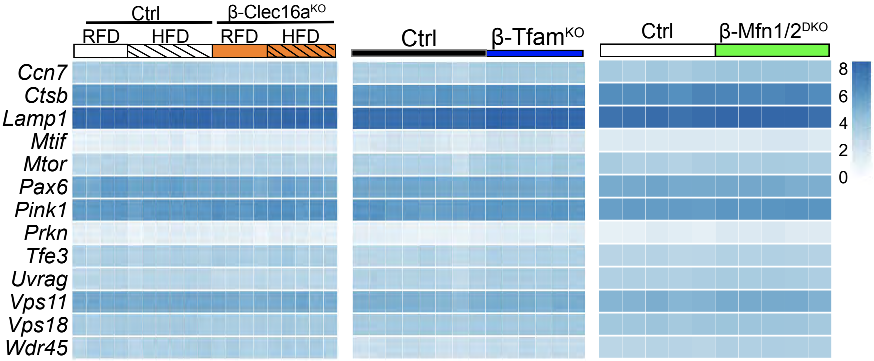
Ctrl
β-Clec16a^KO
RFD
HFD
RFD
HFD
Ctrl
β-Tfam^KO
Ctrl
β-Mfn1/2^DKO
Ccn7
Ctsb
Lamp1
Mtif
Mtor
Pax6
Pink1
Prkn
Tfe3
Uvrag
Vps11
Vps18
Wdr45
8
6
4
2
0

Figure S7

**Figure S7. Parkin and mTOR transcriptional pathways are not changed following mitochondrial quality control defects.** Heat maps representing known Clec16a targets including Parkin and mTOR pathway genes from bulk RNAseq from Ctrl and β-Clec16a$^{KO}$ mice fed HFD or RFD, Ctrl and β-Tfam$^{KO}$, and Ctrl or β-Mfn1/2$^{DKO}$, mice. Represented as transcripts per million (TPM).

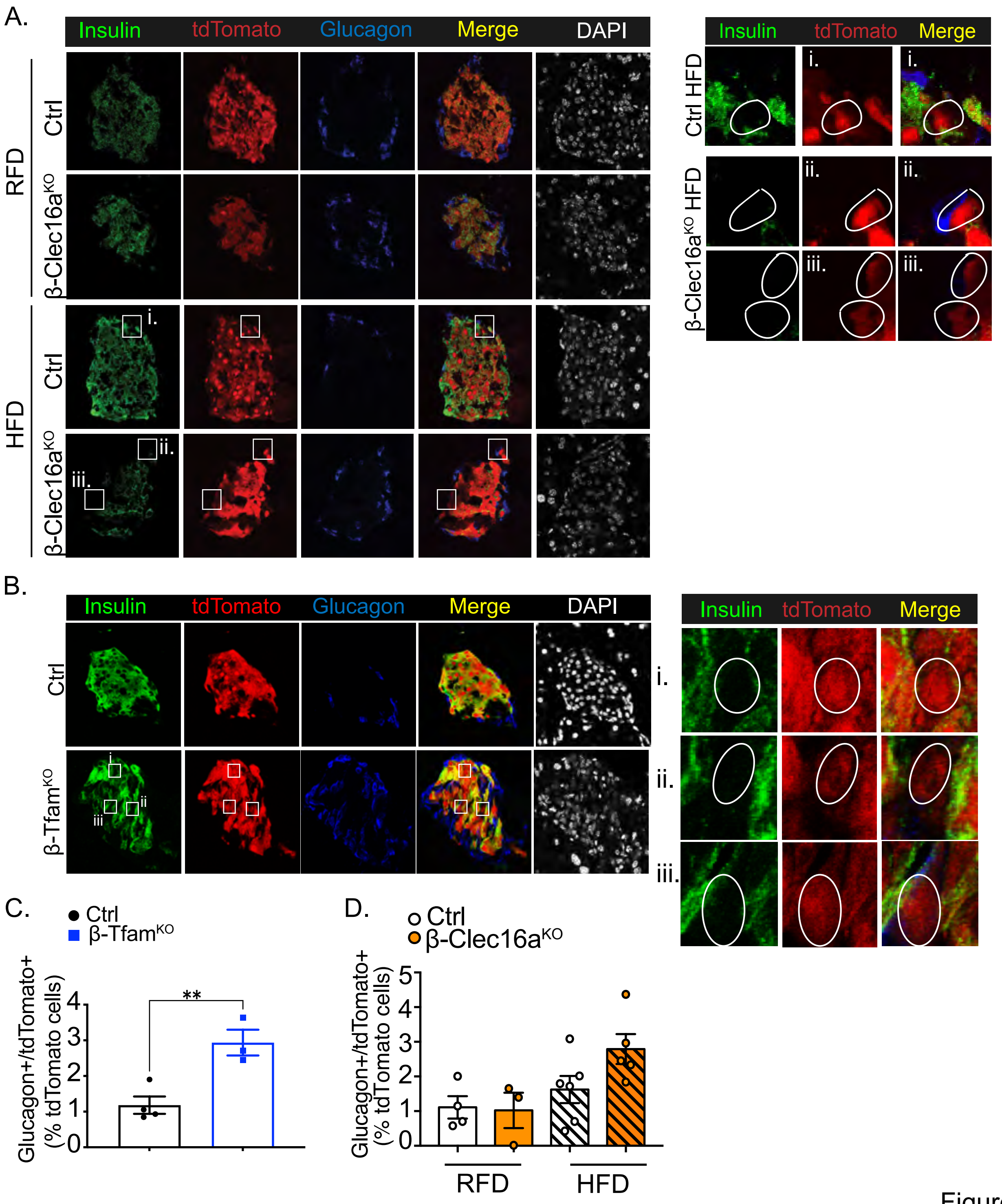
A.
Insulin
tdTomato
Glucagon
Merge
DAPI
RFD
Ctrl
β-Clec16a$^{KO}$
HFD
Ctrl
β-Clec16a$^{KO}$
i.
ii.
iii.
Ctrl HFD
β-Clec16a$^{KO}$ HFD
B.
Insulin
tdTomato
Glucagon
Merge
DAPI
Ctrl
β-Tfam$^{KO}$
i
ii
iii
i.
ii.
iii.
C.
Ctrl
β-Tfam$^{KO}$
Glucagon+/tdTomato+ (% tdTomato cells)
**
0
1
2
3
4
D.
Ctrl
β-Clec16a$^{KO}$
Glucagon+/tdTomato+ (% tdTomato cells)
0
1
2
3
4
5
RFD
HFD

Figure S8

**Figure S8. Lineage tracing studies implicate mitochondrial quality control regulation of β-cell identity.** (A) Representative confocal immunofluorescence images showing staining with anti-Insulin (green), anti-Glucagon (blue) and endogenous tdTomato (red) in *Ins1*-Cre;Rosa26-tdTomato and β-Clec16a$^{KO}$;Rosa26-tdTomato mice. Shown right are magnified images of i, ii, iii white squares. (B) Representative confocal immunofluorescence images showing staining with anti-Insulin (green), anti-Glucagon (blue) and endogenous tdTomato (red) in *Ins1*-Cre;Rosa26-tdTomato and β-Tfam$^{KO}$;Rosa26-tdTomato mice. Shown right are magnified images of i, ii, iii white squares. (C) Quantification of Glucagon-positive tdTomato-positive cells in *Ins1*-Cre;Rosa26-tdTomato and β-Clec16a$^{KO}$;Rosa26-tdTomato mice. $n$ = 3-6 mice per group. (D) Quantification of Glucagon-positive tdTomato-positive cells in *Ins1*-Cre;Rosa26-tdTomato and β-Tfam$^{KO}$;Rosa26-tdTomato mice. $n$ = 3-4 mice per group. **$P$ < 0.01 by Student's unpaired $t$-test.

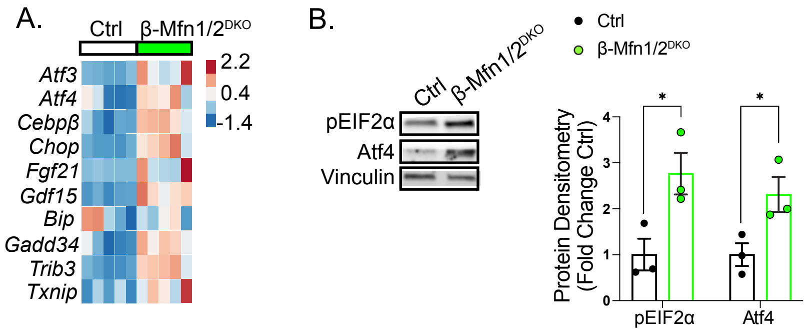
A.
Ctrl
β-Mfn1/2DKO
Atf3
Atf4
Cebpβ
Chop
Fgf21
Gdf15
Bip
Gadd34
Trib3
Txnip
2.2
0.4
-1.4
B.
Ctrl
β-Mfn1/2DKO
pEIF2α
Atf4
Vinculin
Ctrl
β-Mfn1/2DKO
Protein Densitometry (Fold Change Ctrl)
*
*
0
1
2
3
4
pEIF2α
Atf4

Figure S9

**Figure S9: Impaired mitochondrial fusion elicits the ISR.** (A) Heatmap generated from bulk RNA sequencing of RFD Ctrl and β-Mfn1/2$^{DKO}$ islets displaying ISR gene expression. *n* = 5 mice per group. (B) Representative WB and quantitation demonstrating expression of pEIF2α and Atf4 protein of islets from Ctrl or β-Mfn1/2$^{DKO}$ mice on HFD. Vinculin serves a loading control. *n* = 3 mice per group. *$P < 0.05$ by Student's unpaired *t*-test.

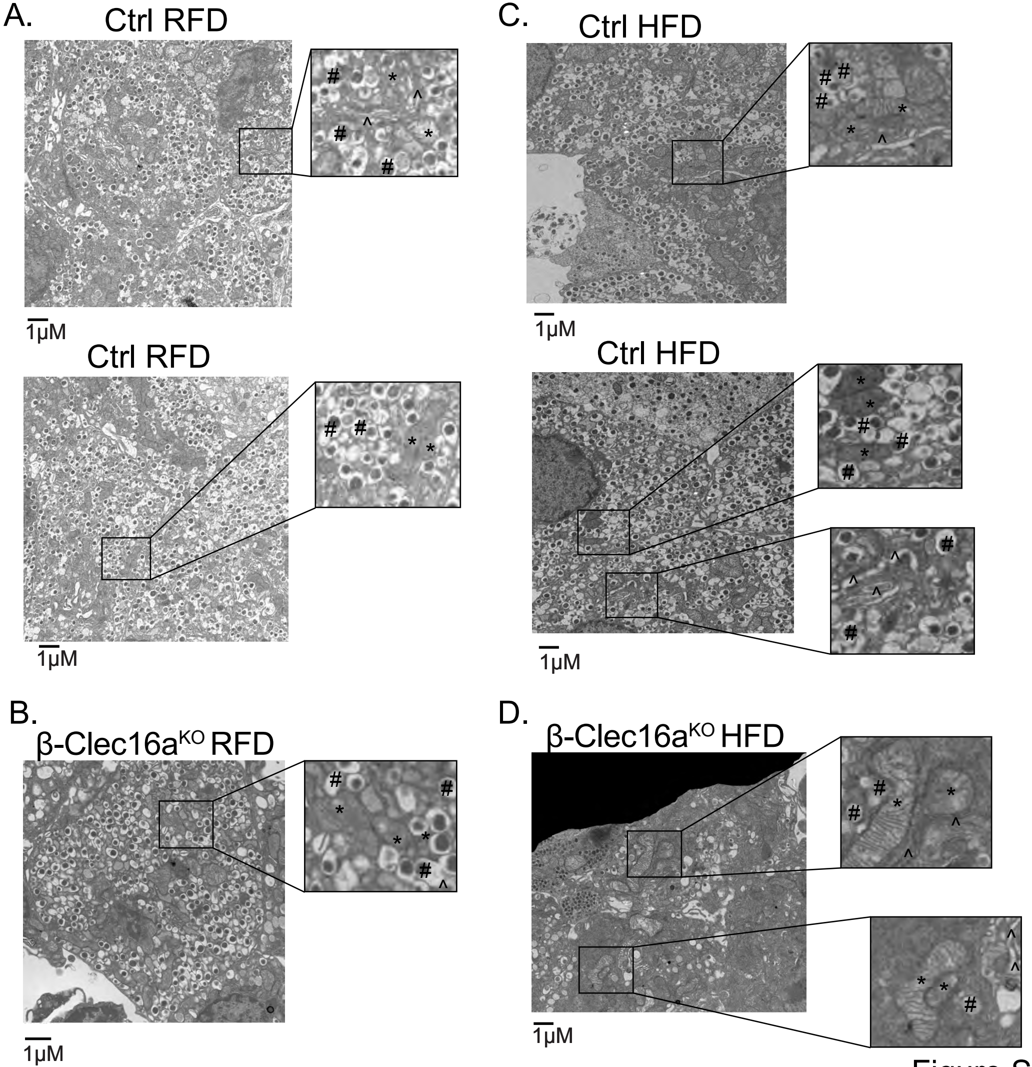
A.
Ctrl RFD
1µM
Ctrl RFD
1µM
B.
β-Clec16a^KO RFD
1µM
C.
Ctrl HFD
1µM
Ctrl HFD
1µM
D.
β-Clec16a^KO HFD
1µM

Figure S10

**Figure S10. Loss of mitophagy does not elicit overt defects in ER morphology.** (A-D) Representative transmission electron microscopy images from islets from Ctrl (A,C) or β-Clec16a$^{KO}$ (B,D) mice fed RFD or HFD as labeled (*n* = 3/group). B-cells are identified by the presence of insulin granules (#); mitochondria indicated with *, and ER indicated with ^.

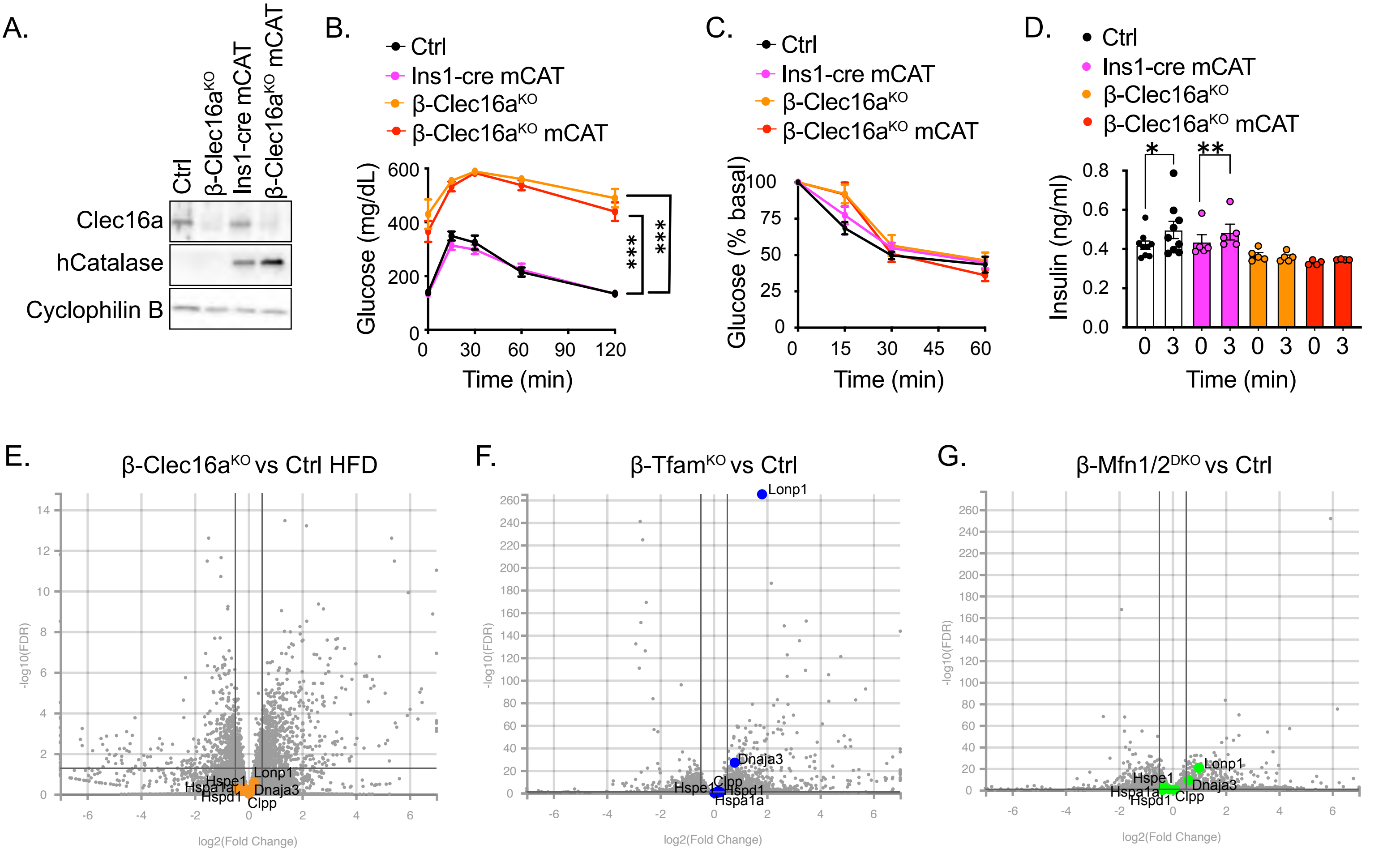
A.
Ctrl
β-Clec16aKO
Ins1-cre mCAT
β-Clec16aKO mCAT
Clec16a
hCatalase
Cyclophilin B
B.
Ctrl
Ins1-cre mCAT
β-Clec16aKO
β-Clec16aKO mCAT
Glucose (mg/dL)
Time (min)
***
***
C.
Ctrl
Ins1-cre mCAT
β-Clec16aKO
β-Clec16aKO mCAT
Glucose (% basal)
Time (min)
D.
Ctrl
Ins1-cre mCAT
β-Clec16aKO
β-Clec16aKO mCAT
Insulin (ng/ml)
Time (min)
*
**
E.
β-Clec16aKO vs Ctrl HFD
-log10(FDR)
log2(Fold Change)
Lonp1
Hspe1
Hspa1a
Hspd1
Dnaja3
Clpp
F.
β-TfamKO vs Ctrl
Lonp1
Dnaja3
Hspe1
Clpp
Hspd1
Hspa1a
G.
β-Mfn1/2DKO vs Ctrl
Lonp1
Hspe1
Dnaja3
Hspa1a
Hspd1
Clpp

Figure S11

**Figure S11. Reducing mitochondrial ROS does not rescue hyperglycemia following defective β-cell mitophagy.** (A) Protein expression of Clec16a and human Catalase (hCatalase) by western blot in β-Clec16a$^{KO}$ and β-Clec16a$^{KO}$-mCAT islets as well as Ctrl and Ins1-Cre-mCAT controls. Cyclophilin B serves as a loading control. (B-D) Results from Ctrl, Ins1-Cre-mCAT, β-Clec16a$^{KO}$ or β-Clec16a$^{KO}$-mCAT mice fed HFD for 12 weeks. (B) Blood glucose concentrations measured during an IPGTT. ***$P$ < 0.01 by two-way ANOVA effect of β-Clec16a$^{KO}$ with or without mCAT overexpression. (C) Blood glucose levels during an ITT. (D) Serum insulin levels measured during *in vivo* glucose-stimulated insulin release in 12-week HFD-fed mice of the groups indicated. *$P$ < 0.05, **$P$ < 0.01 by Student's unpaired $t$-test. (E-G) Volcano plot depicting differential RNA expression in RNAseq studies from islets from HFD-fed β-Clec16a$^{KO}$ €, β-Tfam$^{KO}$ (F), and β-Mfn1/2$^{DKO}$ (G) mice ($n$ = 3-8 mice) compared to their respective littermate controls with UPR$^{mt}$ gene targets highlighted. Significantly differentially expressed genes demarcated by -log10 FDR > or < 2 and log2 fold change (FC) > or < 0.5.

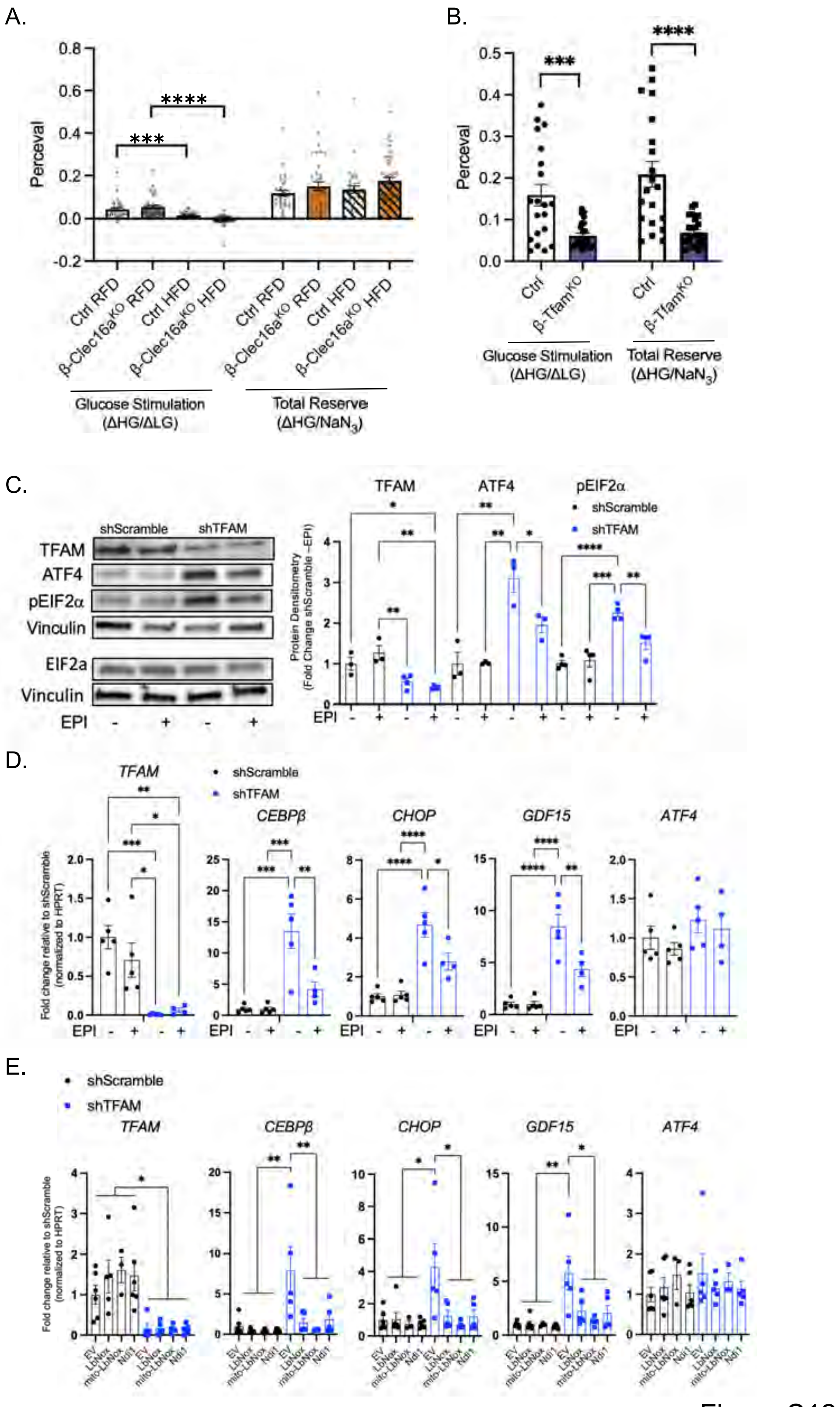
A.
Perceval
Ctrl RFD
β-Clec16a^KO RFD
Ctrl HFD
β-Clec16a^KO HFD
Glucose Stimulation (ΔHG/ΔLG)
Total Reserve (ΔHG/NaN3)
B.
Ctrl
β-Tfam^KO
C.
shScramble
shTFAM
TFAM
ATF4
pEIF2α
Vinculin
EIF2a
EPI
Protein Densitometry (Fold Change shScramble −EPI)
D.
TFAM
CEBPβ
CHOP
GDF15
Fold change relative to shScramble (normalized to HPRT)
E.
EV
LbNox
mito-LbNox
Ndi1

Figure S12

**Figure S12. ETC/OXPHOS system defects contribute to ISR activation following mitochondrial genome instability in β-cells.** (A) ATP/ADP ratio as measured by PercevalHR fluorescence in islets from RFD- or HFD-fed Ctrl or β-Clec16a$^{KO}$ mice stimulated *ex vivo* with high glucose (HG) vs low glucose (LG) levels or treated with $NaN_3$. ***$P$ < 0.001, ****$P$ < 0.0001 by one-way ANOVA, Tukey's multiple comparison post-test. (B) ATP/ADP ratio as measured by PercevalHR fluorescence in Ctrl or β-Tfam$^{KO}$ islets stimulated *ex vivo* with HG vs LG levels or treated with $NaN_3$. ***$P$ < 0.001, ****$P$ < 0.0001 by Student's unpaired *t*-test. (C) Representative WB and quantitation demonstrating expression of TFAM, ATF4, pEIF2α, and EIF2α protein from shScramble or shTFAM EndoC-βH3 cells with or without EPI. Vinculin serves a loading control. *n* = 3-4 group. *$P$ < 0.05, **$P$ < 0.01, ***$P$ < 0.001, ****$P$ < 0.0001 by one-way ANOVA, Tukey's multiple comparison post-test. (D) qPCR analysis of *TFAM* and ISR target genes from shScramble or shTFAM EndoC-βH3 cells with or without EPI. *n* = 5 per group; *$P$ < 0.05, **$P$ < 0.01, ***$P$ < 0.001, ****$P$ < 0.0001 by one-way ANOVA, Tukey's multiple comparison post-test. € qPCR analysis of *TFAM* and ISR target genes from shScramble or shTFAM EndoC-βH3 cells transfected with empty vector (EV), LbNox, mito-LbNox, or Ndi1. *n* = 5-6 per group; *$P$ < 0.05, **$P$ < 0.01 by one-way ANOVA, Tukey's multiple comparison post-test.

A.

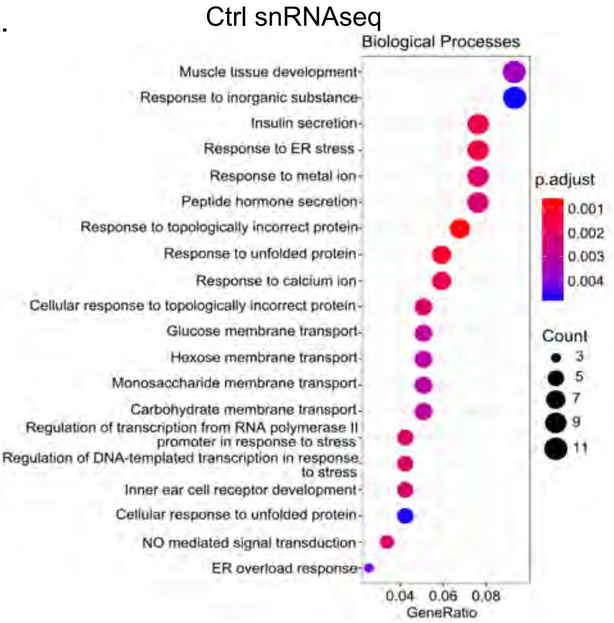


B.

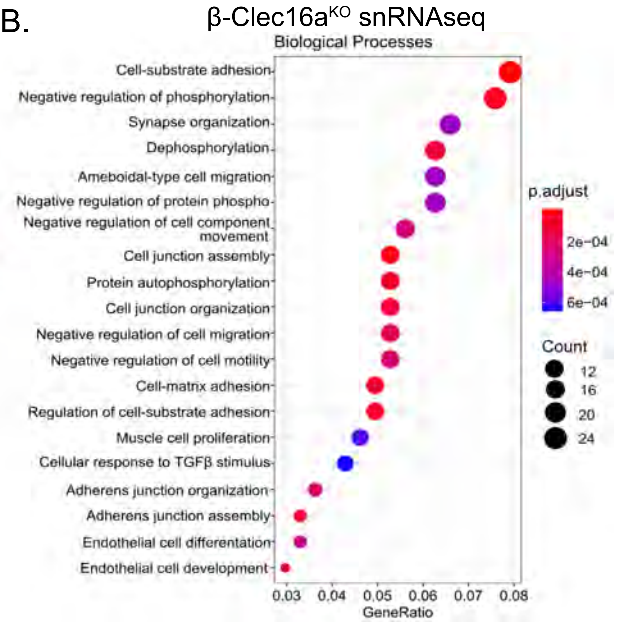


C.

### GREAT predictions: analysis Ctrl specific regions

| Ontology | Term Name | Binom rank | Binom raw p-value | Binom FDR Q-Val | Binom fold enrichment |
|---|---|---|---|---|---|
| GO Biological processes | Phagosome-lysosome fusion | 17 | $1.65955\times10^{-9}$ | $1.27786\times10^{-6}$ | 19.5752 |
| | Vacuolar transport | 38 | $8.04426\times10^{-7}$ | $2.77103\times10^{-4}$ | 2.5396 |
| | Vacuole organization | 42 | $1.19289\times10^{-6}$ | $3.71784\times10^{-4}$ | 2.4592 |
| | Vesicle fusion | 58 | $1.09165\times10^{-5}$ | $2.46374\times10^{-3}$ | 2.4243 |
| | Organelle membrane fusion | 65 | $1.37256\times10^{-5}$ | $2.76413\times10^{-3}$ | 2.3950 |
| | Positive regulation of protein localization to membrane | 86 | $4.40808\times10^{-5}$ | $6.70950\times10^{-3}$ | 2.2476 |
| | Lysosomal transport | 87 | $4.43274\times10^{-5}$ | $6.66949\times10^{-3}$ | 2.4047 |
| | Positive regulation of glucose metabolic process | 195 | $4.46698\times10^{-4}$ | $2.99860\times10^{-2}$ | 2.3844 |
| | Regulation of mitophagy | 196 | $4.58525\times10^{-4}$ | $3.06229\times10^{-2}$ | 2.7972 |
| | Positive regulation calcium ion-dependent exocytosis | 214 | $5.69271\times10^{-4}$ | $3.48213\times10^{-2}$ | 2.9989 |
| | Regulation of hydrogen peroxide-induced cell death | 217 | $5.87078\times10^{-4}$ | $3.54141\times10^{-2}$ | 2.6273 |
| | Regulation of response to reactive oxygen species | 243 | $8.47382\times10^{-4}$ | $4.56470\times10^{-2}$ | 2.3817 |
| Mouse Phenotype | Abnormal cellular replicative senescence | 3 | $4.27033\times10^{-7}$ | $1.36295\times10^{-3}$ | 2.9149 |
| | Delayed cellular replicative senescence | 8 | $5.79861\times10^{-6}$ | $6.94021\times10^{-3}$ | 10.6798 |

D.

### GREAT predictions: analysis of β-Clec16a$^{KO}$ specific regions

| Ontology | Term Name | Binom rank | Binom raw p-value | Binom FDR Q-Val | Binom fold enrichment |
|---|---|---|---|---|---|
| GO Biological processes | Cellular response to carbohydrate stimulus | 201 | $3.3679\times10^{-20}$ | $2.19318\times10^{-18}$ | 2.0103 |
| | Cellular response to monosaccharide stimulus | 235 | $1.70267\times10^{-17}$ | $9.48474\times10^{-16}$ | 2.0276 |
| | Cellular response to hexose stimulus | 241 | $5.83611\times10^{-17}$ | $3.16991\times10^{-15}$ | 2.0182 |
| | Axo-dendritic transport | 270 | $1.15510\times10^{-15}$ | $5.60009\times10^{-14}$ | 2.4480 |
| | Axonal transport | 282 | $4.44435\times10^{-15}$ | $2.06300\times10^{-13}$ | 2.6881 |
| | Retrograde axonal transport | 293 | $1.20488\times10^{-14}$ | $5.38289\times10^{-13}$ | 4.3619 |
| | Positive regulation of transcription from RNA polymerase II promoter involved in cellular response to chemical stimulus | 470 | $4.25571\times10^{-10}$ | $1.18526\times10^{-8}$ | 2.0915 |
| | Negative regulation of epithelial to mesenchymal transition | 474 | $4.99883\times10^{-10}$ | $1.38048\times10^{-8}$ | 2.1833 |
| | Positive regulation of nucler-transcribed mRNA catabolic process, deadenylation-dependent decay | 610 | $2.44932\times10^{-8}$ | $1.49603\times10^{-6}$ | 2.4695 |
| | Positive regulation of nuclear-transcribed mRNA poly(A) tail shortening | 676 | $9.73723\times10^{-8}$ | $1.88551\times10^{-6}$ | 3.0060 |
| Mouse Phenotype | Abnormal pancreas development | 313 | $1.79172\times10^{-13}$ | $5.48107\times10^{-12}$ | 2.0386 |
| | Abnormal pancreatic beta cell differentiation | 371 | $6.59305\times10^{-12}$ | $1.70158\times10^{-10}$ | 2.1148 |
| | Abnormal nitric oxide homeostasis | 404 | $3.94434\times10^{-11}$ | $9.34827\times10^{-10}$ | 2.1981 |
| | Increased circulating iron level | 425 | $1.27681\times10^{-10}$ | $2.87658\times10^{-9}$ | 2.1049 |
| | Abnormal fibroblast migration | 239 | $1.91674\times10^{-17}$ | $7.67899\times10^{-15}$ | 2.1269 |
| | Decreased fibroblast cell migration | 204 | $3.95059\times10^{-18}$ | $1.85426\times10^{-16}$ | 2.2820 |

Figure S13

**Figure S13. Single nuclei RNA- and ATAC-sequencing identifies unique β-cell populations with distinct transcriptomic and chromatin landscapes following impaired mitochondrial quality control.** (A) Biological process pathway analysis showing upregulated pathways unique to the Ctrl β-cell population from snRNA sequencing. (B) Biological process pathway analysis showing upregulation pathways unique to the β-cell population from β-Clec16a$^{KO}$ mice by snRNAseq. (C) GREAT pathway analysis of the unique open chromatin peaks from the Ctrl β-cell population. (D) GREAT pathway analysis of the unique open chromatin peaks from the β-cell population from β-Clec16a$^{KO}$ mice.

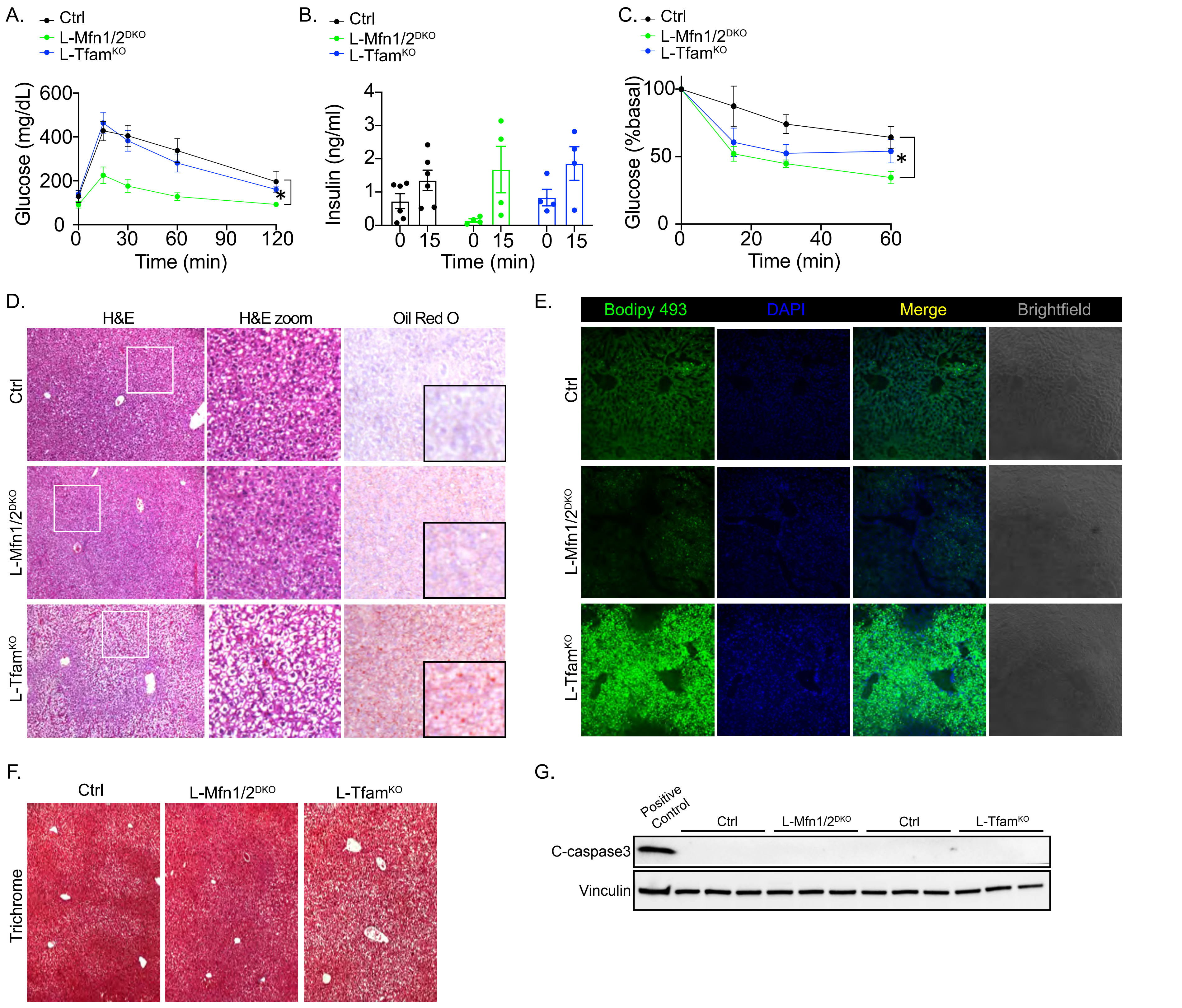
A.
Ctrl
L-Mfn1/2DKO
L-TfamKO
Glucose (mg/dL)
Time (min)
B.
Insulin (ng/ml)
C.
Glucose (%basal)
D.
H&E
H&E zoom
Oil Red O
E.
Bodipy 493
DAPI
Merge
Brightfield
F.
Trichrome
G.
Positive Control
C-caspase3
Vinculin

Figure S14

**Figure S14. Mitochondrial quality control perturbations in the liver affect insulin sensitivity, glycemic control, and/or lipid accumulation.** (A-C) Metabolic results from AAV8-Tbg$^{cre}$ Ctrl, L-Mfn1/2$^{DKO}$ or L-Tfam$^{KO}$ mice, *n* = 4-6 mice/group for all studies. (A) Blood glucose concentrations during an IPGTT 4 weeks post-AAV injection. **P* < 0.05 by two-way ANOVA effect of L-Mfn1/2$^{DKO}$ vs Ctrl. (B) Serum insulin levels measured during an IPGTT 4 weeks post-AAV injection at 0- or 15-min post glucose bolus. (C) Change in blood glucose levels during an ITT 3 weeks post-AAV injection. **P* < 0.05 by two-way ANOVA effect of L-Mfn1/2$^{DKO}$ vs Ctrl. (D) H&E and Oil Red O staining of liver from Ctrl, L-Mfn1/2$^{DKO}$ or L-Tfam$^{KO}$ liver sections. (E) Representative immunofluorescence images of lipid accumulation in liver from Ctrl, L-Mfn1/2$^{DKO}$ or L-Tfam$^{KO}$ liver sections; lipid (Bodipy 493) in green, nuclei (DAPI) in blue, and whole section brightfield image (grey). (F) Trichrome staining of liver from Ctrl, L-Mfn1/2$^{DKO}$ or L-Tfam$^{KO}$ liver sections. (G) WB for cleaved Caspase 3 in Ctrl, L-Mfn1/2$^{DKO}$ or L-Tfam$^{KO}$ liver. *n* = 3-6 mice per group. WB includes CDDO treated Min6 cells as a positive control. Vinculin serves as a loading control.

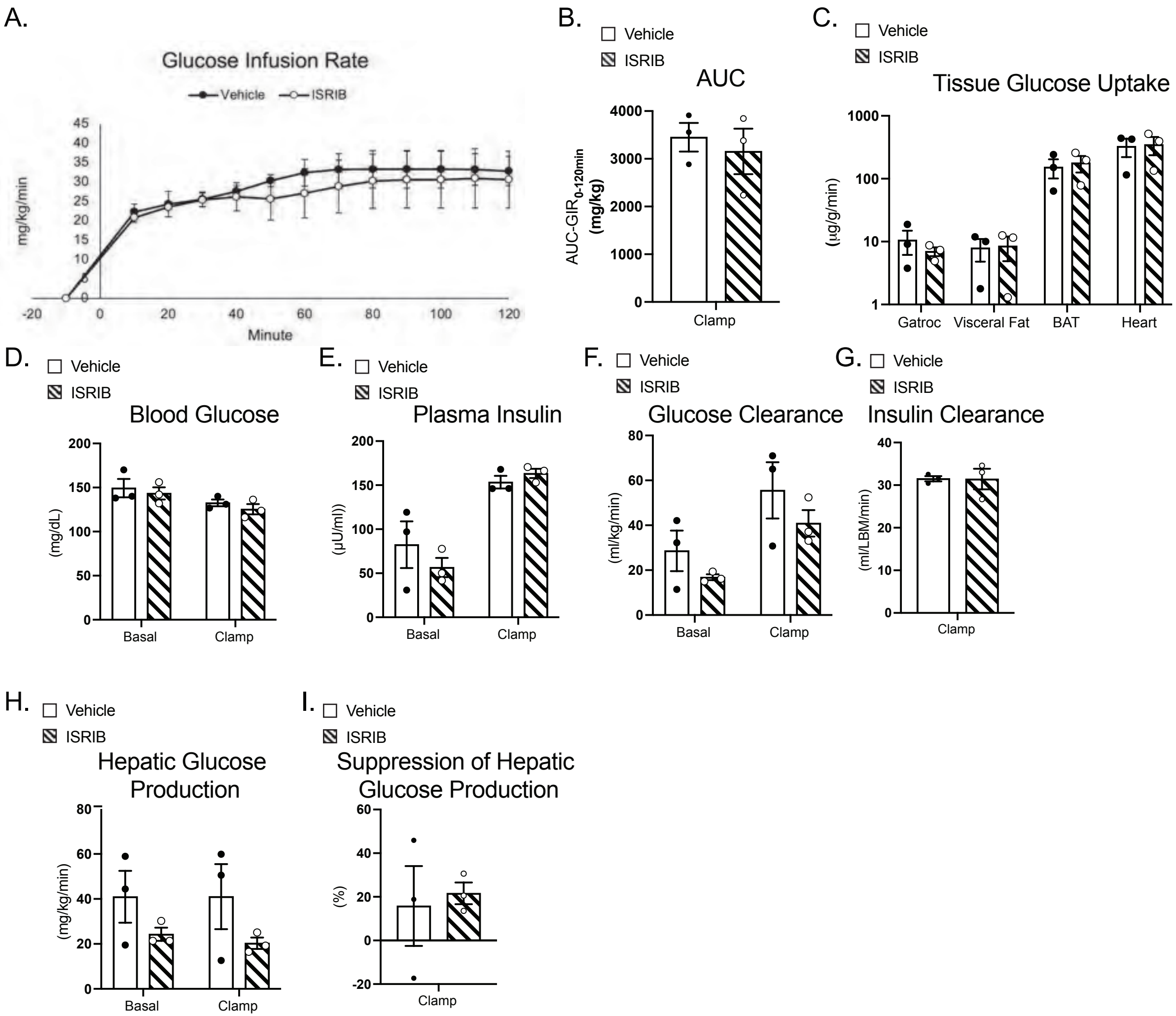
A.
Glucose Infusion Rate
Vehicle
ISRIB
mg/kg/min
Minute
B.
AUC
AUC-GIR0-120min (mg/kg)
Clamp
C.
Tissue Glucose Uptake
(µg/g/min)
Gatroc
Visceral Fat
BAT
Heart
D.
Blood Glucose
(mg/dL)
Basal
Clamp
E.
Plasma Insulin
(µU/ml)
F.
Glucose Clearance
(ml/kg/min)
G.
Insulin Clearance
(ml/LBM/min)
H.
Hepatic Glucose Production
(mg/kg/min)
I.
Suppression of Hepatic Glucose Production
(%)

Figure S15

**Figure S15. ISRIB does not alter multiple parameters of blood glucose regulation during hyperinsulinemic-euglycemic clamp studies.** (A-B) Glucose infusion rate (A) and AUC (B) in Vehicle or ISRIB treated control mice during hyperinsulinemic-euglycemic clamp. (C) Glucose uptake measured using portal vein infused tracers in various tissues. (D) Basal and clamped blood glucose levels in Vehicle or ISRIB treated control mice. (E) Basal and clamped plasma insulin levels in Vehicle or ISRIB treated control mice. (F) Basal and clamped glucose clearance rates in Vehicle or ISRIB treated control mice. (G) Clamped insulin clearance rates in Vehicle or ISRIB treated control mice. (H) Basal and clamped hepatic glucose production rates in Vehicle or ISRIB treated control mice. (I) Clamped hepatic glucose production suppression rates (%) in Vehicle or ISRIB treated control mice.

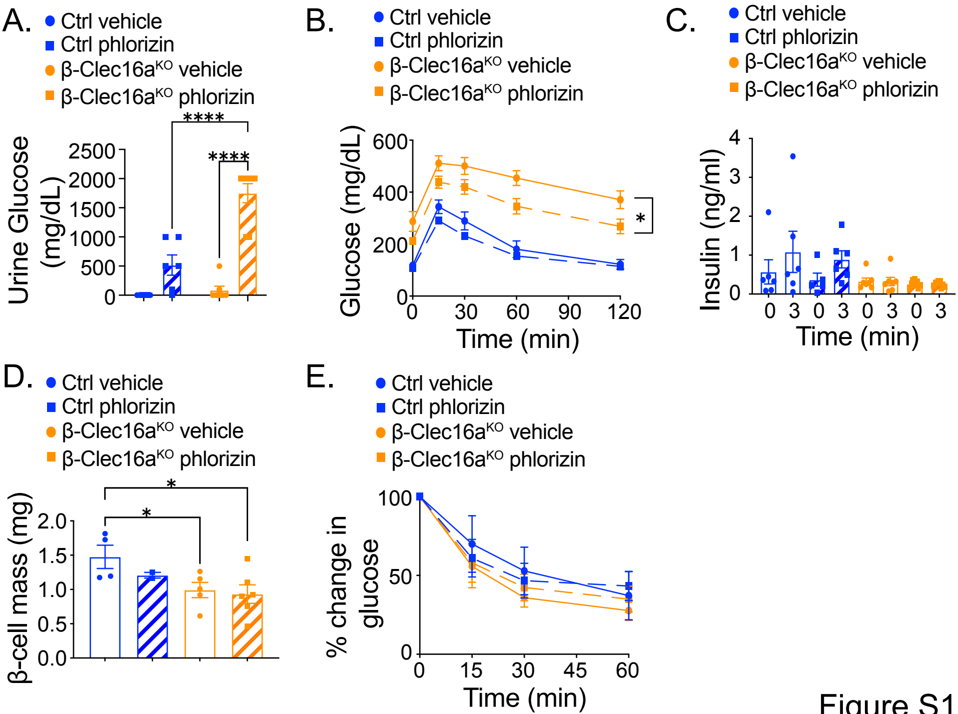

A.
Ctrl vehicle
Ctrl phlorizin
β-Clec16aKO vehicle
β-Clec16aKO phlorizin
Urine Glucose (mg/dL)
2500
2000
1500
1000
500
0
****
****
B.
Glucose (mg/dL)
600
400
200
0
Time (min)
0
30
60
90
120
*
C.
Insulin (ng/ml)
4
3
2
1
0
Time (min)
0 3 0 3 0 3 0 3
D.
β-cell mass (mg)
2.0
1.5
1.0
0.5
0.0
*
*
E.
% change in glucose
100
50
0
Time (min)
0
15
30
45
60


Figure S16

**Figure S16. Reducing hyperglycemia does not improve β-cell function/mass following mitophagy deficiency.** (A-E) Ctrl or β-Clec16a$^{KO}$ mice fed HFD for 12-weeks treated with vehicle or phlorizin via mini-osmotic pump implantation for 6-weeks. *n* = 4-6/group for all studies. (A) Urine glucose concentrations measured 5 weeks after pump implantation. ****$P < 0.0001$ by two-way ANOVA, Tukey's multiple comparison post-test. (B) Blood glucose concentrations during an IPGTT. *$P < 0.05$ by two-way ANOVA effect of drug within genotype. (C) Serum insulin levels measured during *in vivo* glucose-stimulated insulin release. (D) Pancreatic β-cell mass. *$P < 0.05$ by Student's unpaired *t*-test. (E) Blood glucose levels during an ITT.

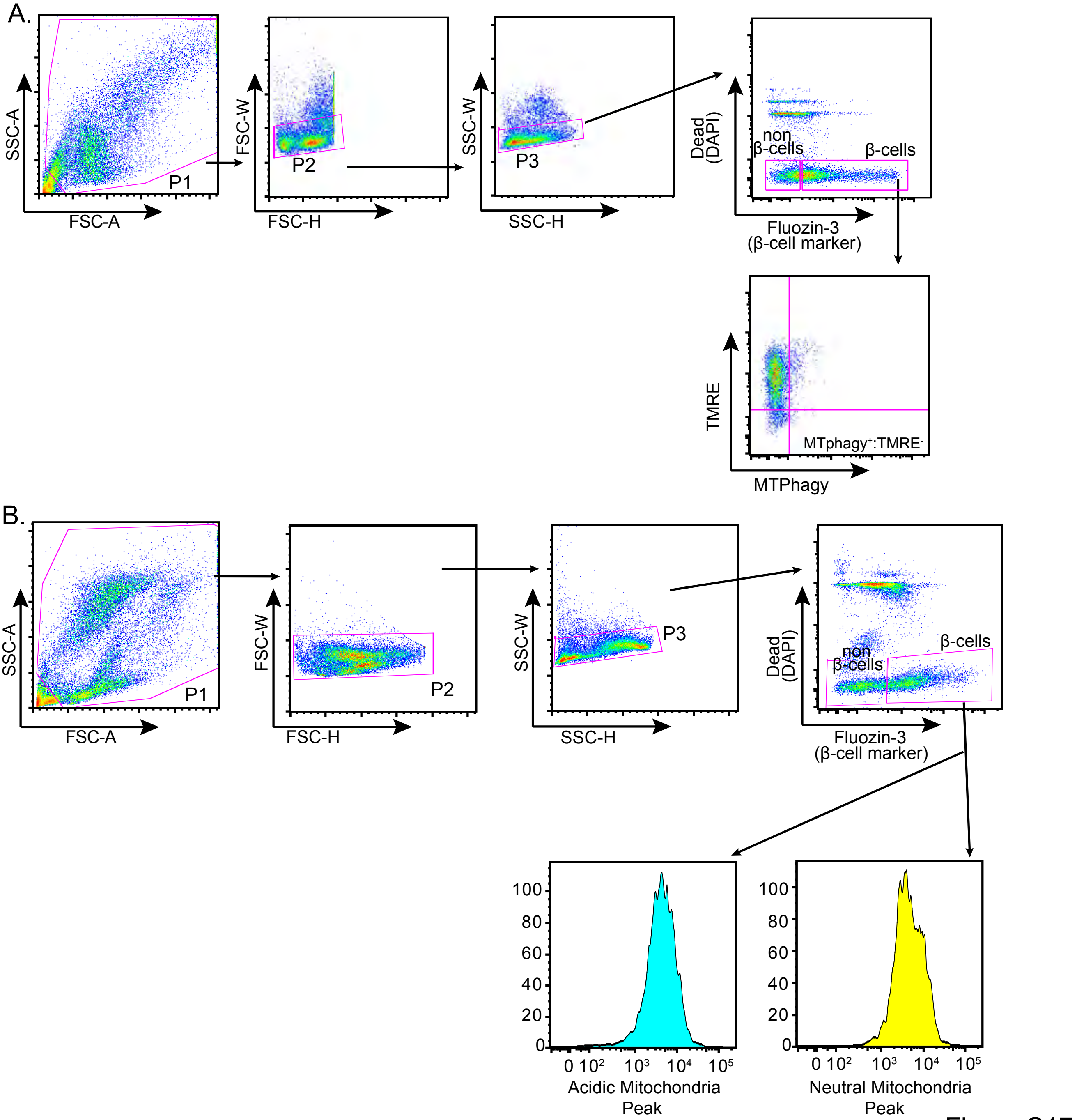
A.
SSC-A
FSC-A
P1
FSC-W
FSC-H
P2
SSC-W
SSC-H
P3
Dead (DAPI)
non β-cells
β-cells
Fluozin-3 (β-cell marker)
TMRE
MTPhagy
MTphagy⁺:TMRE⁻
B.
SSC-A
FSC-A
P1
FSC-W
FSC-H
P2
SSC-W
SSC-H
P3
Dead (DAPI)
non β-cells
β-cells
Fluozin-3 (β-cell marker)
100
80
60
40
20
0
0 10² 10³ 10⁴ 10⁵
Acidic Mitochondria Peak
100
80
60
40
20
0
0 10² 10³ 10⁴ 10⁵
Neutral Mitochondria Peak

Figure S17

**Figure S17. Flow cytometry gating strategy.** (A) Representative flow cytometry plots from dispersed human islets, showing doublet removal (P2 and P3), identification of live (DAPI negative) cells gated by Fluozin-3 staining (to identify $Zn^{2+}$ enriched β-cell populations), and followed by measurement of $MTphagy^{HI}$:$TMRE^{LO}$ populations, which designates cells undergoing mitophagy. (B) Representative flow cytometry plots from dispersed murine islets from mt-Keima transgenic mice, showing doublet removal (P2 and P3), identification of live (DAPI negative) gated by Fluozin-3 staining, and measurement of acidic to neutral mitochondria histograms, used to quantify cells undergoing mitophagy.

**Table S1:** Gene expression, reported by log2(FC) and FDR, for shared up- and down-regulated genes in β-Clec16a$^{KO}$, Tfam$^{KO}$, and Mfn1/2$^{DKO}$ islets compared to respective littermate controls.

| Gene Name | ENSEMBL mouse | Clec16a HFD log2(FC) | FDR | TFAM log2(FC) | FDR | Mfn1/2 log2(FC) | FDR |
|---|---|---|---|---|---|---|---|
| Abtb2 | ENSMUSG00000032724 | 1.14 | 2.43E-02 | 0.79 | 1.54E-07 | 1.21 | 1.66E-09 |
| Adm | ENSMUSG00000030790 | 3.48 | 5.92E-06 | 2.68 | 4.32E-06 | 2.44 | 1.34E-10 |
| Adm2 | ENSMUSG00000054136 | 4.40 | 1.62E-06 | 4.08 | 5.66E-62 | 1.04 | 1.22E-03 |
| Agap1 | ENSMUSG00000055013 | 0.80 | 3.19E-04 | 0.97 | 4.93E-47 | 0.91 | 4.77E-33 |
| Agpat4 | ENSMUSG00000023827 | 1.14 | 6.33E-06 | 0.84 | 7.68E-05 | 1.65 | 1.87E-24 |
| Akna | ENSMUSG00000039158 | 1.20 | 1.29E-03 | 2.76 | 3.24E-104 | 1.54 | 6.24E-18 |
| Angptl6 | ENSMUSG00000038742 | 2.54 | 2.30E-05 | 3.22 | 8.72E-136 | 0.90 | 2.03E-05 |
| Ap1s2 | ENSMUSG00000031367 | -0.78 | 6.21E-03 | -0.96 | 1.52E-07 | -0.70 | 2.67E-09 |
| Areg | ENSMUSG00000029378 | 6.86 | 1.35E-09 | 7.12 | 2.61E-14 | 5.94 | 1.61E-252 |
| Arhgef2 | ENSMUSG00000019467 | 0.40 | 6.47E-03 | 0.81 | 2.43E-04 | 0.81 | 4.65E-01 |
| Arl14ep | ENSMUSG00000027122 | 0.89 | 2.94E-05 | 1.35 | 5.64E-65 | 0.75 | 9.24E-19 |
| Astn1 | ENSMUSG00000026587 | 1.86 | 9.33E-06 | 1.13 | 2.22E-06 | 1.13 | 9.25E-08 |
| Atf3 | ENSMUSG00000026628 | 1.79 | 4.60E-04 | 3.88 | 8.21E-46 | 1.78 | 9.64E-05 |
| B3gnt9 | ENSMUSG00000069920 | -0.72 | 3.02E-03 | -0.89 | 6.14E-06 | -1.77 | 2.13E-16 |
| Bambi-ps1 | ENSMUSG00000081219 | -1.85 | 3.16E-03 | -0.80 | 4.85E-03 | -2.34 | 2.52E-30 |
| Brinp2 | ENSMUSG00000004031 | 1.44 | 2.23E-06 | 1.33 | 2.55E-20 | 1.06 | 3.82E-06 |
| C2cd4b | ENSMUSG00000091956 | -0.93 | 1.77E-07 | -0.84 | 9.92E-08 | -0.59 | 4.14E-03 |
| Car10 | ENSMUSG00000056158 | -0.53 | 1.14E-02 | -0.60 | 4.02E-05 | -0.67 | 9.67E-09 |
| Cebpb | ENSMUSG00000056501 | 0.98 | 6.30E-03 | 1.63 | 1.70E-45 | 0.72 | 2.92E-04 |
| Coro6 | ENSMUSG00000020836 | 0.96 | 8.81E-03 | 1.35 | 6.90E-04 | 2.10 | 9.82E-08 |
| Cryl1 | ENSMUSG00000021947 | -1.35 | 1.85E-06 | -0.81 | 1.60E-09 | -0.73 | 8.03E-08 |
| Ddit3 | ENSMUSG00000025408 | 0.76 | 1.05E-02 | 2.79 | 2.58E-79 | 1.81 | 1.93E-19 |
| Dmpk | ENSMUSG00000030409 | -0.86 | 2.35E-07 | -0.94 | 1.84E-09 | -0.60 | 1.17E-04 |
| Dnajc24 | ENSMUSG00000027166 | -1.04 | 2.64E-03 | -0.74 | 7.18E-07 | -0.87 | 5.58E-14 |
| Dock5 | ENSMUSG00000044447 | 0.92 | 1.89E-03 | 2.76 | 3.02E-123 | 1.51 | 3.83E-16 |
| Extl1 | ENSMUSG00000028838 | 1.76 | 1.92E-02 | 3.78 | 1.16E-28 | 1.65 | 2.49E-05 |
| Fam166a | ENSMUSG00000026969 | -0.91 | 1.44E-03 | -1.48 | 5.56E-12 | -1.25 | 6.29E-06 |
| Fgf21 | ENSMUSG00000030827 | 2.79 | 1.85E-02 | 3.07 | 7.76E-14 | 2.49 | 3.57E-04 |
| Fut1 | ENSMUSG00000008461 | 3.85 | 2.22E-06 | 4.32 | 4.52E-82 | 2.50 | 1.83E-70 |
| G0s2 | ENSMUSG00000009633 | 1.80 | 1.40E-03 | 1.24 | 1.65E-12 | 2.96 | 5.57E-18 |
| Gbp11 | ENSMUSG00000092021 | 1.42 | 1.18E-02 | 2.81 | 9.17E-05 | 2.68 | 1.04E-23 |
| Gdf15 | ENSMUSG00000038508 | 4.67 | 4.76E-06 | 2.91 | 7.44E-40 | 0.72 | 3.76E-07 |
| Gipr | ENSMUSG00000030406 | -1.05 | 5.85E-04 | -1.27 | 2.29E-20 | -0.89 | 2.68E-06 |
| Gpr135 | ENSMUSG00000043398 | 1.58 | 5.91E-05 | 1.74 | 8.90E-26 | 1.88 | 1.55E-09 |
| Gpt2 | ENSMUSG00000031700 | 1.18 | 1.83E-04 | 1.52 | 5.30E-06 | 1.85 | 7.76E-45 |
| Hfe | ENSMUSG00000006611 | 0.90 | 1.73E-04 | 1.94 | 7.50E-47 | 0.84 | 4.74E-06 |

| | | | | | | | |
|---|---|---|---|---|---|---|---|
| Hrk | ENSMUSG00000046607 | 5.52 | 5.07E-04 | 3.46 | 2.39E-08 | 3.36 | 1.23E-21 |
| Igfbp5 | ENSMUSG00000026185 | 1.60 | 4.02E-08 | 0.77 | 1.15E-03 | 1.18 | 4.17E-10 |
| Inhbb | ENSMUSG00000037035 | 1.42 | 6.79E-04 | 1.00 | 6.38E-04 | 0.91 | 4.50E-04 |
| Ins2 | ENSMUSG00000000215 | -0.45 | 3.01E-02 | -0.95 | 5.87E-19 | -1.09 | 2.39E-13 |
| Itpkb | ENSMUSG00000038855 | -0.76 | 5.77E-10 | -0.79 | 1.44E-14 | -0.81 | 1.81E-13 |
| Izumo1 | ENSMUSG00000064158 | 3.59 | 5.15E-03 | 2.03 | 6.51E-06 | 2.08 | 6.24E-12 |
| Kcna2 | ENSMUSG00000040724 | 1.00 | 8.68E-03 | 1.13 | 6.08E-06 | 1.52 | 2.49E-08 |
| Lhfpl2 | ENSMUSG00000045312 | 1.39 | 2.21E-05 | 2.87 | 1.28E-35 | 2.09 | 1.13E-33 |
| Lmo1 | ENSMUSG00000036111 | 1.90 | 3.08E-06 | 3.13 | 3.88E-54 | 2.29 | 6.39E-47 |
| Loxl2 | ENSMUSG00000034205 | -0.45 | 2.77E-02 | -0.70 | 1.40E-08 | -0.87 | 4.28E-20 |
| Mafa | ENSMUSG00000047591 | -0.63 | 1.09E-02 | -0.66 | 2.32E-03 | -1.34 | 1.60E-10 |
| Mei4 | ENSMUSG00000043289 | 3.15 | 1.18E-02 | 3.33 | 5.30E-12 | 1.62 | 3.50E-03 |
| Metrn | ENSMUSG00000002274 | 1.17 | 4.86E-03 | 1.06 | 1.28E-05 | 0.99 | 9.37E-05 |
| Mob3b | ENSMUSG00000073910 | 0.86 | 3.17E-02 | 1.30 | 3.20E-11 | 1.09 | 4.25E-04 |
| Ngf | ENSMUSG00000002881 | 0.45 | 2.44E-02 | 0.48 | 5.73E-04 | 1.94 | 2.34E-03 |
| Niban1 | ENSMUSG00000026483 | 3.32 | 2.21E-08 | 3.00 | 9.54E-70 | 1.32 | 2.21E-10 |
| Npas3 | ENSMUSG00000021010 | 0.87 | 3.80E-02 | 0.77 | 4.45E-06 | 0.87 | 8.33E-06 |
| Nupr1 | ENSMUSG00000030717 | 1.38 | 1.54E-02 | 4.89 | 1.45E-51 | 3.29 | 2.01E-05 |
| Olfm4 | ENSMUSG00000022026 | -0.66 | 1.53E-02 | -0.73 | 4.35E-07 | -0.64 | 2.14E-03 |
| Opcml | ENSMUSG00000062257 | 3.00 | 9.41E-07 | 1.26 | 1.34E-02 | 2.73 | 1.68E-05 |
| Or2t48 | ENSMUSG00000050818 | 4.61 | 2.25E-06 | 5.33 | 2.71E-83 | 3.77 | 3.15E-36 |
| Paqr3 | ENSMUSG00000055725 | 1.18 | 7.62E-04 | 1.61 | 7.26E-12 | 1.77 | 9.52E-29 |
| Pfkfb2 | ENSMUSG00000026409 | -0.62 | 3.55E-02 | -0.86 | 2.06E-21 | -0.86 | 2.43E-19 |
| Pfn4 | ENSMUSG00000020639 | 1.08 | 5.86E-03 | 1.08 | 4.36E-15 | 1.17 | 3.90E-11 |
| Phactr1 | ENSMUSG00000054728 | -0.79 | 3.26E-04 | -1.02 | 9.67E-11 | -1.25 | 2.72E-26 |
| Plcxd1 | ENSMUSG00000064247 | 1.15 | 1.36E-02 | 1.82 | 3.37E-15 | 0.96 | 1.54E-03 |
| Prss1 | ENSMUSG00000062751 | 2.86 | 1.09E-02 | 2.44 | 3.82E-02 | 7.04 | 4.30E-03 |
| Ptger3 | ENSMUSG00000040016 | 1.79 | 5.17E-03 | 2.67 | 2.36E-15 | 2.45 | 9.44E-16 |
| Rassf2 | ENSMUSG00000027339 | 1.06 | 2.38E-02 | 2.10 | 4.58E-28 | 1.75 | 1.39E-20 |
| Rcc2 | ENSMUSG00000040945 | 1.02 | 3.08E-05 | 0.81 | 4.78E-24 | 0.66 | 9.91E-13 |
| Scarf2 | ENSMUSG00000012017 | 2.76 | 1.72E-05 | 1.41 | 2.27E-11 | 1.17 | 9.14E-18 |
| Scd3 | ENSMUSG00000025202 | 0.90 | 9.11E-03 | 2.08 | 4.92E-32 | 1.26 | 4.69E-10 |
| Sesn2 | ENSMUSG00000028893 | 0.91 | 1.73E-04 | 1.20 | 2.62E-23 | 0.82 | 1.53E-28 |
| Sgip1 | ENSMUSG00000028524 | 2.87 | 1.18E-06 | 2.17 | 6.95E-12 | 3.57 | 1.55E-54 |
| Sh2b2 | ENSMUSG00000005057 | 1.34 | 3.51E-02 | 1.01 | 3.09E-04 | 0.89 | 6.18E-03 |
| Slc16a6 | ENSMUSG00000041920 | 0.93 | 3.96E-03 | 0.91 | 2.14E-22 | 0.89 | 1.58E-20 |
| Slc1a7 | ENSMUSG00000008932 | 3.97 | 3.57E-03 | 6.50 | 6.04E-27 | 3.59 | 5.12E-12 |
| Slc2a2 | ENSMUSG00000027690 | -1.27 | 1.22E-02 | -0.99 | 1.07E-10 | -1.59 | 1.37E-24 |
| Slc6a9 | ENSMUSG00000028542 | 1.19 | 1.97E-02 | 2.12 | 2.92E-32 | 0.93 | 1.81E-06 |

| | | | | | | | |
|---|---|---|---|---|---|---|---|
| Slc7a3 | ENSMUSG00000031297 | 5.43 | 3.38E-12 | 3.48 | 4.75E-32 | 1.60 | 1.10E-03 |
| Slitrk6 | ENSMUSG00000045871 | -1.08 | 3.52E-04 | -1.00 | 1.70E-15 | -1.38 | 3.03E-34 |
| Snx33 | ENSMUSG00000032733 | 0.85 | 1.75E-04 | 0.84 | 5.55E-16 | 0.90 | 9.73E-25 |
| Sorcs2 | ENSMUSG00000029093 | 1.29 | 5.10E-03 | 0.82 | 7.43E-05 | 0.64 | 2.32E-04 |
| Stard5 | ENSMUSG00000046027 | 1.10 | 3.83E-06 | 2.39 | 2.04E-83 | 1.61 | 2.60E-22 |
| Stbd1 | ENSMUSG00000047963 | 1.98 | 6.60E-06 | 4.31 | 1.28E-105 | 2.60 | 1.54E-15 |
| Stc2 | ENSMUSG00000020303 | 4.88 | 1.71E-07 | 4.68 | 1.36E-46 | 4.40 | 1.11E-57 |
| Steap1 | ENSMUSG00000015652 | 1.52 | 5.08E-04 | 3.47 | 3.27E-153 | 1.65 | 1.25E-08 |
| Sytl4 | ENSMUSG00000031255 | -1.17 | 6.57E-03 | -0.63 | 4.61E-11 | -1.21 | 6.77E-21 |
| T2 | ENSMUSG00000058159 | -2.42 | 2.35E-05 | -0.90 | 4.22E-04 | -0.95 | 8.24E-06 |
| Tenm4 | ENSMUSG00000048078 | 2.42 | 3.55E-06 | 3.45 | 1.68E-109 | 2.50 | 1.95E-51 |
| Th | ENSMUSG00000000214 | -1.48 | 2.49E-13 | -0.81 | 2.97E-06 | -0.96 | 3.26E-05 |
| Tmem215 | ENSMUSG00000046593 | -1.06 | 9.43E-06 | -1.16 | 3.75E-37 | -1.38 | 2.40E-20 |
| Trib1 | ENSMUSG00000032501 | -0.65 | 1.26E-02 | -1.36 | 2.51E-11 | -0.84 | 4.50E-24 |
| Trib3 | ENSMUSG00000032715 | 3.58 | 6.75E-08 | 5.15 | 5.04E-42 | 1.80 | 4.86E-16 |
| Trim66 | ENSMUSG00000031026 | 3.26 | 6.84E-08 | 2.72 | 3.22E-39 | 1.99 | 4.33E-84 |
| Trp53cor1 | ENSMUSG00000085912 | 1.13 | 1.10E-03 | 1.69 | 9.97E-20 | 0.71 | 1.18E-12 |
| Trpm5 | ENSMUSG00000009246 | -2.28 | 2.94E-02 | -1.43 | 2.34E-04 | -1.15 | 4.14E-09 |
| Ubap1l | ENSMUSG00000086228 | 3.21 | 2.59E-03 | 5.70 | 5.72E-93 | 1.93 | 6.46E-12 |
| Ucn3 | ENSMUSG00000044988 | -1.05 | 2.04E-08 | -0.89 | 1.17E-23 | -1.37 | 1.97E-20 |
| Unc5b | ENSMUSG00000020099 | 1.95 | 2.41E-05 | 1.62 | 1.91E-17 | 1.52 | 1.58E-19 |
| Usp51 | ENSMUSG00000067215 | -0.66 | 2.58E-02 | -0.77 | 2.93E-06 | -0.73 | 3.26E-11 |
| Vil1 | ENSMUSG00000026175 | -0.48 | 3.55E-03 | -0.63 | 4.60E-09 | -0.65 | 3.29E-11 |
| Vmn2r3 | ENSMUSG00000091572 | 6.31 | 2.01E-04 | 9.63 | 3.27E-37 | 6.19 | 6.85E-76 |
| Vrtn | ENSMUSG00000071235 | 5.10 | 3.29E-03 | 9.62 | 7.95E-40 | 3.98 | 1.18E-23 |
| Wipf3 | ENSMUSG00000086040 | 1.27 | 1.46E-05 | 1.98 | 1.45E-48 | 1.82 | 9.60E-18 |
| Zfp189 | ENSMUSG00000039634 | 0.83 | 2.52E-02 | 1.53 | 2.41E-45 | 0.79 | 2.37E-09 |

**Table S2:** Detailed donor information for human islets used for all studies within the manuscript.

| Unique Islet Prep Identifer | Islet Isolation Center | Age | BMI | Sex | Cause of Death |
|---|---|---|---|---|---|
| ADHH356 | IIDP | 52 | 25.9 | Female | Cerebrovascular/stroke |
| ADID386 | IIDP | 30 | 22.4 | Male | Head trauma |
| AEIY348 | Southern California Islet Cell Resource Center | 24 | 38.1 | Male | Cerebrovascular/stroke |
| AEJQ100 | The Scharp-Lacy Research Institute | 56 | 13.8 | Male | Cerebrovascular/stroke |
| AEJT193 | University of Wisconsin | 51 | 29.0 | Male | Head trauma |
| AELK219 | University of Pennsylvania | 37 | 38.1 | Female | Cerebrovascular/stroke |
| AFFC135 | IIDP | 32 | 28.5 | Male | Head trauma -gunshot wound suicide |
| ARJR491 | University of Miami | 38 | 33.0 | Male | Head trauma |
| H1081 | IIDP | 44 | 32.5 | Male | Undisclosed |
| HP-22030-01 | Prodo labs | 36 | 25.6 | Male | Anoxic event |
| HP-22034-01 | Prodo labs | 49 | 32.5 | Male | Anoxic event |
| HP-22052-01 | Prodo labs | 52 | 25.4 | Male | Anoxic event |
| HP-22065-01 | Prodo labs | 61 | 22.4 | Male | Anoxic event |
| HP-22187-01 | Prodo labs | 47 | 27.9 | Male | Anoxic event |
| HP14149-01 | Prodo labs | 59 | 21.5 | Male | Anoxia/cardiovascular arrest |
| HP14178-01 | Prodo labs | 25 | 36.9 | Female | Cerebrovascular/stroke |
| HP14199-01 | Prodo labs | 34 | 27.5 | Female | Cerebrovascular/stroke |
| HP14205-01 | Prodo labs | 52 | 22.5 | Male | Cerebrovascular/stroke |
| HP15135-01 | Prodo labs | 29 | 23.0 | Male | ICH |
| HP15219-01 | Prodo labs | 25 | 22.8 | Male | Head trauma |
| HP15298-01T2D | Prodo labs | 42 | 43.0 | Female | ICH |
| HP16012-01T2D | Prodo labs | 48 | 43.7 | Male | Anoxic event with CPR |
| HP16023-01T2D | Prodo labs | 51 | 24.4 | Male | Stroke |
| HP16272-01 | The Scharp-Lacy Research Institute | 52 | 29.8 | Male | Head trauma |
| HP17117-01 T2D | Prodo labs | 51 | 35.6 | Male | Cerebral vascular accident/brain death |
| HP1720901T2D | Prodo labs | 52 | 42.8 | Female | Stroke |
| HP17225-01T2D | The Scharp-Lacy Research Institute | 59 | 27.7 | Male | Cerebrovascular/stroke |
| HP17300-01T2D | Prodo labs | 35 | 27.7 | Male | Stroke |
| HP18054-01 | The Scharp-Lacy Research Institute | 40 | 25.3 | Male | Head trauma |
| HP20259-01T2D | Prodo labs | 59 | 25.1 | Male | Head trauma |
| ID1189 | IIDP | 15 | 23.0 | Male | Head trauma |
| ID1199 | IIDP | 50 | 28.5 | Female | Cerebrovascular/stroke |
| ID1276 | IIDP | 19 | 20.0 | Male | Head trauma |
| ID1330 | IIDP | 53 | 22.0 | Female | Cerebrovascular/stroke |
| R385 | ADI Islet Core, Alberta | 47 | 31.0 | Male | Neurological |
| R386 T2D | ADI Islet Core, Alberta | 43 | 35.8 | Female | Neurological |
| R426 | ADI Islet Core, Alberta | 33 | 31.9 | Female | Neurological |
| R427 | ADI Islet Core, Alberta | 52 | 35.8 | Male | Medical Assistance in Dying |
| R434 | ADI Islet Core, Alberta | 50 | 24.2 | Female | Neurological |
| R436 | ADI Islet Core, Alberta | 57 | 39.7 | Female | Medical Assistance in Dying |
| R451 | ADI Islet Core, Alberta | 56 | 24.0 | Female | Neurological |
| R510 | ADI Islet Core, Alberta | 53 | 21.3 | Female | Undisclosed |

**Table S3:** Demographic and metabolic comparison of human islet donors from mtDNA content studies in Fig. 1A.

| | T2D; n=9 | ND; n=24 | |
|---|---|---|---|
| **Age** | 48.89 | 41.42 | |
| Stdev | 9.37 | 12.82 | |
| | | t-test | 0.1331 |
| **BMI** | 33.98 | 27.05 | |
| Stdev | 7.42 | 4.78 | |
| | | t-test | **0.0010 |
| **HbA1C** | 6.73 | 5.34 | |
| Stdev | 0.65 | 0.19 | |
| | | t-test | ***1.3855E-08 |
| **Sex** | | | |
| Males | n=6 (66.67%) | n=15 (62.50%) | |
| Females | n=3 (33.33%) | n=9 (37.50%) | |

**Table S4:** Reported medications for human islet donors with T2D for studies in Fig. 1A.

| Islet Prep Identifier | Age | BMI | Sex | A1C | Medication |
|---|---|---|---|---|---|
| HP15298-01T2D | 42 | 43.0 | Female | 6.5 | No medication |
| HP16012-01T2D | 48 | 43.7 | Male | 6.6 | Medication |
| HP16023-01T2D | 51 | 24.4 | Male | 6.9 | No medication |
| HP17117-01 T2D | 51 | 35.6 | Male | 7.1 | 4y oral medications |
| HP1720901T2D | 52 | 42.8 | Female | 6.6 | Newly diagnosed |
| HP17225-01T2D | 59 | 27.7 | Male | 6.5 | Metformin-6y / T2D-10y |
| HP17300-01T2D | 35 | 27.7 | Male | 6.3 | Metformin-3y / T2D-3y |
| AELK219 | 37 | 38.1 | Female | 8.2 | 0-5y oral medications |
| SAMN12634037 | 63 | 31.7 | Male | 5.9 | Metformin-5y / T2D 6-10y |

**Table S5:** Mouse and human primer sequences.

Mouse

| Gene | Forward | Reverse |
|---|---|---|
| Bip | TTCAGCCAATTATCAGCAAACTCT | TTTTCTGATGTATCCTCTTCACCAGT |
| Cebpb | GCAATCCGGATCAAACGTG | AACAACCCCGCAGGAACAT |
| Hprt | GGCCAGACTTTGTTGGATTTG | TGCGCTCATCTTAGGCTTTGT |
| mt9/mt11 | GAGCATCTTATCCACGCTTCC | GGTGGTACTCCCGCTGTAAA |
| Ndufv | CTTCCCCACTGGCCTCAAG | CCAAAACCCAGTGATCCAGC |
| Trib3 | TCAGCAACTGTGAGAGGACGA | TGCTTGTCCCACAGAGAGTCA |
| Ucn3 | AGCTGCAACCCTGAACAGTCA | TCAGCATCGCTCCCTGTAAGT |

Human

| Gene | Forward | Reverse |
|---|---|---|
| ATF4 | ACGTTGCCATGATCCCTCAGT | TGGGCTCATACAGATGCCACT |
| BIP | TGAAGCCCGTCCAGAAAGTGT | ATTCGAGTCGAGCCACCAACA |
| CEBPB | TGGGAATCTTTTCCGTTTCAAGCA | CTGCCCCCAAAAGGCTTTGTA |
| CHOP | GCGCATGAAGGAGAAAGAACAGG | TCGATTTCCTGCTTGAGCCGT |
| FGF21 | GGGAGTCAAGACATCCAGGT | GGCTTCGGACTGGTAAACAT |
| GAPDH | AATCCCATCACCATCTTCCA | TGGACTCCACGACGTACTCA |
| GDF15 | GACCCTCAGAGTTGCACTCC | GCCTGGTTAGCAGGTCCTC |
| GLUT1 | GGACAGGCTCAAAGAGGTTATG | AGGAGGTGGGTGGAGTTAAT |
| HPRT | GATTTTATCAGACTGAAGAGC | TCCAGTTAAAGTTGAGAGATC |
| INS | GAAGCGTGGCATTGTGGAACA | GCTGCGTCTAGTTGCAGTAGTTC |
| MAFA | TGAGCGGAGAACGGTGATTTCTAAGG | GGAACGGAGAACCACGTTCAACGTA |
| MT-CO1 | CGCCACACTCCACGGAAGCA | CGGGGCATTCCGGATAGGCC |
| MT-CO2 | AGAACCAGGCGACCTGCGAC | CCCCCGGTCGTGTAGCGGTG |
| MT-CO3 | CACTGGCCCCCAACAGGCAT | AGTATCAGGCGGCGGCTTCGA |
| MT-ND3 | TTACGAGTGCGGCTTCGACC | ACTCATAGGCCAGACTTAGG |
| MT-ND5 | TCGAATAATTCTTCTCACCC | TAGTAATGAGAAATCCTGCG |
| PDX1 | TACTGGATTGGCGTTGTTTGTGGC | AGGGAGCCTTCCAATGTGTATGGT |
| TFAM | ATGGCGTTTCTCCGAAGCAT | TCCGCCCTATAAGCATCTTGA |

**Table S6:** Antibody information.

| Antibody | Company | Catalogue Number |
|---|---|---|
| AFP | R&D | MAB1368-SP |
| Aldh1a3 | Novus | NBP2-15339 |
| ATF4 | Pierce/Thermo | PA5-19521 |
| C-peptide | DSHB | GN-ID4 |
| Clec16a | n/a | Developed in Soleimanpour lab |
| Cyclophilin B | Invitrogen | PA1-027A |
| Cyp2e1 | Abcam | ab28146 |
| EIF2a | Cell Signaling | 2103S |
| Glucagon | Sigma | G2654 |
| Glut2 | Millipore | 07-1402-I |
| Insulin | Abcam | ab7842 |
| Mfn1 | Abcam | ab126575 |
| Mfn2 | Abcam | ab56889 |
| Mup3 | Novus | NBP2-61648 |
| OXPHOS (human) | Abcam | ab110411 |
| OXPHOS (mouse) | Abcam | ab110413 |
| Pdx1 | Abcam | ab47308 |
| pEIF2a | Cell Signaling | 5199S |
| Psat1 | ProteinTech | 10501-1-AP |
| TFAM (human) | PhosphoSolutions | 1999-hTFAM |
| Tfam (mouse) | PhosphoSolutions | 2001-Tfam |
| Tom20 | Cell Signaling | 42406S |
| Top2a | Abcam | ab52934 |
| Ucn3 | n/a | Gift of Wylie Vale and Joan Vaughan, The Salk Institute for Biological Sciences |
| Vinculin | Millipore | CP74-100UG |
